# NSP4 mutation T492I drives rapid evolution of SARS-CoV-2 toward Omicron

**DOI:** 10.1101/2024.07.05.602217

**Authors:** Xiaoyuan Lin, Zhou Sha, Chunlin Zhang, Julia M. Adler, Ricardo Martin Vidal, Christine Langner, Beibei Fu, Yan Xiong, Meng Tan, Chen Jiang, Hao Zeng, Xiaokai Zhang, Qian Li, Jingmin Yan, Xiaoxue Lu, Shiwei Wang, Xuhu Mao, Dusan Kunec, Jakob Trimpert, Haibo Wu, Quanming Zou, Zhenglin Zhu

## Abstract

T492I, a mutation encountered in SARS-CoV-2 nonstructural protein 4 (NSP4), enhances viral replication and alters nonstructural protein cleavage, causing potential evolutionary impacts. Through comprehensive comparative analyses based on evolve-and-resequence experiments of SARS-CoV-2 wild-type and Delta strains with or without T492I, we demonstrate that NSP4 T492I not only increases the mutation rate, but also accelerates the emergence of many mutations characteristic for Omicron variants. Accordingly, viral populations that evolved from ancestors with T492I, show Omicron-biased selective forces and increases in viral replication, infectivity, immune evasion capacity, potentials for cross-species transmission and receptor-binding affinity. Aside from enhanced replication, we observed stronger epistasis regarding viral replication and infectivity in T492I than in S N501Y and NSP6 ΔSGF; this facilitates the regulation of mutation types, which can drive fast evolution of Omicron specific mutations. Our results highlight the role of an important replication-enhancing mutation in regulating the evolutionary rates and mutational trends of SARS-CoV-2.

## INTRODUCTION

The SARS-CoV-2 Omicron strain is the latest variant of concern (VOC), giving rise to the fourth wave of the COVID-19 pandemic. Variants of this lineage are still predominant around the world ^1–3^. Compared to previously emerged VOCs and variants of interest (VOIs), Omicron is heavily mutated ^3^, resulting in considerable conformational alterations, which cause increased transmissibility and immune evasion capability compared with Alpha and Delta ^4, 5^. In the same time, mutations encountered in Omicron variants are thought to result in attenuated pathology ^6^ as well as changes in the binding affinity to host cell enzymes ^7, 8^. Subvariants of the Omicron lineage exhibit substantially impaired cell-cell fusion capacity and tend to use the endosomal entry pathway, rather than the plasma membrane entry pathway mediated by transmembrane protease serine 2 (TMPRSS2) predominantly observed in earlier variants ^9–11^. This change is possibly associated with the tropism of Omicron toward the upper airway epithelium and away from lung tissue ^12^. Collectively, these adaptations may have caused the sudden emergence and fixation of Omicron variants within about two months, which is faster than in any previous VOCs or VOIs ^13–15^. However, the origin of the Omicron variant remains obscure. This VOC contains a number of mutations rarely seen in previous VOCs or VOIs, particularly changes observed in the spike protein (S) distinguish this variant from previously circulating variants ^16^. Identification of Omicron’s origins is of more than academic importance and may help to prevent the pandemic spread of new variants possibly emerging in the future ^15^.

Previous work suggests that the S protein, the nucleocapsid protein (N), the non-structural protein 4 (NSP4) and NSP6 facilitate the functional adaption of Omicron variants ^17–20^. The T492I mutation within the non-structural protein 4 (NSP4) in SARS-CoV-2 is associated with increased replication capacity and accelerated viral replication ^18^. Thus, variants bearing T492I theoretically undergo more replication cycles and introduce more mutations to the genome than the wild-type virus within a transmission cycle. T492I may endow more opportunities to develop adaptive mutations (such as S D614G and N R203K/G304R) ^19–21^ to support the transmission advantage and adaptability of SARS-CoV-2. Mutations may shape future evolution by epistasis ^22^, modulating the effects of mutations at other sites ^23^, such as the epistatic shifts caused by the spike N501Y substitution in the effects of receptor-binding domain (RBD) mutations ^14, 23–25^. Our previous *in silico* analysis of the highly transmissible Delta sub-variant 21J (with 492I) and the less transmissible variants 21A+21I (with T492) predicted an additive or possibly synergistic ^26^ effect of T492I in fitness ^18^. Moreover, T492I increases the cleavage efficiency of the viral main protease NSP5 by enhancing enzyme−substrate binding, resulting in an increased production of most NSPs processed by NSP5. The NSPs are known to constitute the viral replication and transcription complex, interact with host proteins during the early coronavirus replication cycle, and initiate the biogenesis of replication organelles ^27, 28^. Consequently, the T492I mutation, as an adaptive mutation involved in replication, transcription and the capacity of immune evasion, may cause emergence and increased selection of other adaptive mutations. Furthermore, as T492I is fixed both in Delta and Omicron lineages, this mutation may also be a major contributor to the evolutionary processes causing the swift replacement of Delta by Omicron.

To evaluate the hypotheses formulated above, we performed evolve and resequence experiments of replicate SARS-CoV-2 populations propagated by serial passaging on Calu-3 cells in a time course of 45 and 90 days (15 and 30 transmission cycles), respectively. We resequenced and compared the populations evolved from the wild-type strain and those from the T492I mutant. We also performed the same experiment for the VOC Delta and compared the evolved populations resulting from Delta ancestors with T492I and that without T492I. The results show that the T492I mutation not only induced an elevated evolutionary rate, but also an accumulation of dominant mutations, mostly mutations characteristic for the Omicron lineage. By assaying viral subgenomic RNA and genomic RNA, we further demonstrated that the populations evolved from ancestors with T492I (492I runs) show an increase of viral replication and infection, compared with the populations evolved from the ancestors without T492I (T492 runs). We also demonstrated that 429I runs have an increased immune evasion capacity and receptor-binding affinity compared to T492 runs by the detection of interferon production, ELISAs (enzyme-linked immunosorbent assay) and SPR (surface plasmon resonance) experiments. Through evaluating the performance of virus in the cells expressing the ACE2 orthologs of different species, we found that the populations evolved from the 492I runs have higher cross-species infection potentials than those of the T492 runs. Experiments conducted in cell lines demonstrated that the combined effect of NSP4 T492I and other adaptive mutations in infection and viral replication is stronger than that for S N501Y and NSP6 ΔSGF, suggesting strong epistatic effects of T492I. Population genomic analyses showed biased nucleotide mutation types and positive selection toward Omicron specific mutations induced by T492I. Biased nucleotide mutation types were found to result from the up-regulation of APOBEC (apolipoprotein B mRNA editing enzyme catalytic subunit) and down-regulation of ADAR (adenosine deaminase acting on RNA enzymes) in viruses harboring T492I. In conclusion, our findings demonstrate the capability of T492I to accelerate the evolution of SARS-CoV-2 and suggest an important role of this mutation in evolution of Omicron variants. We further identify forces that drive this development, including the enhancement of replication, epistasis effects and mutation type changes induced by T492I.

## RESULTS

### T492I induces rapid and Omicron-biased evolution

To evaluate the effects of T492I on the evolution of SARS-CoV-2, we constructed four variants as ancestors for our evolution experiments: the T492I mutant (aWT-I), which is based on the wild-type (WT) strain (GenBank accession No. MT020880); the wild-type strain (aWT-T); the Delta variant (EVAg: 009V-04187, aDelta-I); and the reversely mutated (I492T) Delta variant (aDelta-T). We chose aDelta-I and aDelta-T as ancestors in the evolution experiment to evaluate the effects of T492I on strains with adaptive mutations, such as Delta mutations. Experimental evolution was carried out over an incubation course of 90 days (30 transmission events) with three independent replicates performed in parallel (R1, R2 and R3) (Figure 1A), resulting in corresponding evolved populations (eWT-I, eWT-T, eDelta-I and eDelta-T). To develop a better understanding of the evolutionary process and confirm the findings, we also independently performed evolve-and-resequence experiments on the four constructed variants in a shorter experiment of 45 days (15 transmission events). RNA from the evolved viruses was extracted and analyzed via whole-genome sequencing (WGS). Through variant detection of the sequencing data via a Bayesian statistical framework ^29^ (requiring a frequency > 0.01), we identified 753 and 611 mutations in populations that underwent experimental evolution for 90 days (90-day runs) and those subject to experimental evolution for 45 days (45-day runs), respectively. In both the 90-day and 45-day runs, eWT-I accumulated more mutations than eWT-T did, and the same effect was observed when 45-day eDelta-I and 45-day eDelta-T were compared (Figure 1B). The 90-day eWT-I has a median mutation rate (0.000116 per base per transmission event, nt^-1^ T^-1^), which is 4.36-fold higher than that of the 90-day eWT-T (0.000027 nt^-1^ T^-1^), and the 90-day eDelta-I has a median mutation rate (0.000133 nt^-1^ T^-1^), which is 2.63-fold higher than that of the 90-day eDelta-T (0.000051 nt^-1^ T^-1^). Similarly, the 45-day 492I runs (eWT-I and eDelta-I) have a 3∼4-fold higher mutation rate than the 45-day T492 runs (eWT-T and eDelta-T, Figure 1C). This finding demonstrates the increase in SARS-CoV-2 mutation rates conveyed by T492I. Accordingly, global surveys of full-length SARS-CoV-2 genomes also revealed a greater number of mutations in the 492I variants than in the T492 variants (Figure 1D and Supplementary Figure S1A) before the emergence of VOCs (from April 2020 to November 2020).

**Figure 1.**
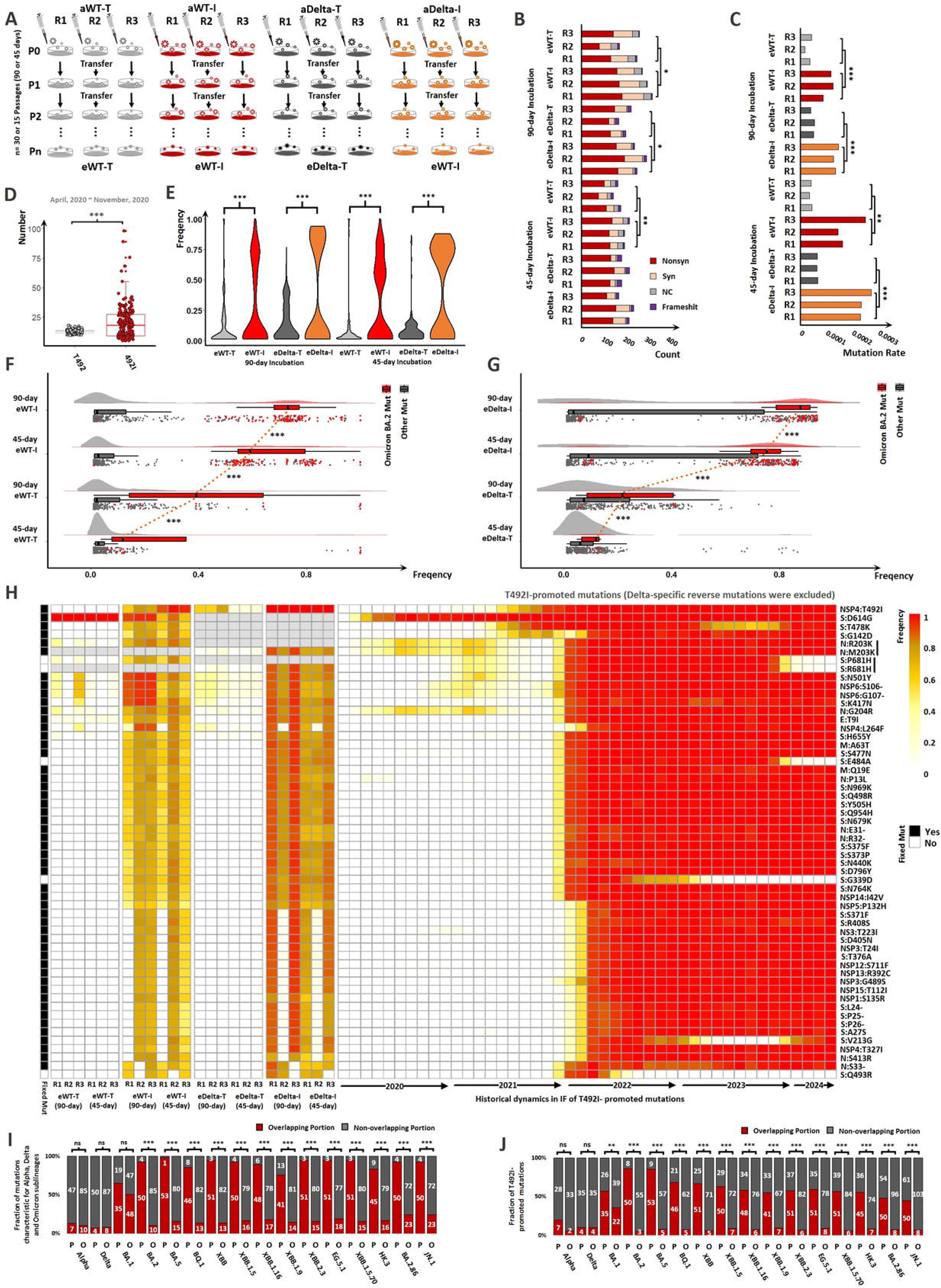
Evidence suggesting Omicron-biased evolution induced by T492I. (A) Graphic overview of evolve-and-resequence experiments. The ancestors are the wild-type strain bearing T492I (aWT-I), the wild-type strain (aWT-T), the Delta variant (aDelta-I) and the Delta variant without T492I (aDelta-T), and the evolved populations are eWT-I, eWT-T, eDelta-I and eDelta-T, respectively. Each run has three replicates (R1, R2 and R3). (B) Counts of nonsynonymous (Nonsyn), synonymous (Syn), noncoding region (NC) and frameshift mutations (Frameshift) in different evolved populations. The statistics for the comparison of mutation counts between the 492I and T492 runs was performed via ANOVA tests. (C) Comparison of mutation rates in different evolved populations. (D) Comparison of the number of amino acid substitutions between the 492I variants and the T492 variants from April 2020 to November 2020 on the basis of global SARS-CoV-2 epidemiological data. (E) Distributions of mutation frequencies in different evolved populations. (F, G) are the comparison in the distribution of mutation frequencies between eWT-I and eWT-T (F) and that between eDelta-I and eDelta-T (G). Raincloud, box and scatter plots display the distributions of the mutations characteristic for Omicron BA.2 (red) and other mutations (grey). (H) Heatmap displays the frequencies of the T492I-promoted mutations (Delta-specific reverse mutations were excluded). Other Delta-specific mutations in eWT-T and eWT-I are colored by gray, and so do the WT-specific mutations in eDelta-T and eDelta-I. The historical global IFs of these mutations is shown on the right. The column ‘Fixed’ on the left denotes the mutations already fixed to date. In (I), P denotes the fractions of T492I-promoted mutations (Delta-specific reverse mutations were excluded) in the mutations characteristic for the Alpha, Delta and Omicron sublineages (VOC mutations), and O denotes the fractions of other high-frequency mutations (with a >50% frequency in a replicate) in VOC mutations. In (J), P denotes the fractions of VOC mutations in T492I-promoted mutations, and O denotes those in other high-frequency mutations. The overlapping portions are marked by red. The statistics for the comparisons between P and O was performed by Chi-squared tests. ‘*’ denotes a P-value < 0.1, ‘**’ denotes a P-value <0.05, and ‘***’ denotes a P-value <0.01.

For both the 90-day– and 45-day-evolved populations, the mutations we identified were mostly nonsynonymous (Binomial Test, P-value < 2.2e-16, Figure 1B) and single nucleotide polymorphisms (SNPs, Binomial Test, P-value < 2.2e-16, Supplementary Figure S1B). The 492I runs have a greater fraction of high-frequency nonsynonymous mutations than the T492 runs, both in the 90-day and 45-day replicates (Figures 1E-G and S1C). A biased distribution toward high-frequency mutations was also found for synonymous and noncoding region mutations (Supplementary Figure S1D, E). The higher frequencies of the mutations observed in the 90-day 492I runs than in in the 45-day 492I runs suggest an evolutionary dynamic induced by T492I (Supplementary Figure S1F). There were 139 high-frequency nonsynonymous mutations with a > 0.5 frequency in one or more replicates of the 90-day and 45-day runs (Supplemental Table S1). Among these high frequency mutations, 78 were identified with a significantly greater frequency in the 492I runs than in the T492 runs (please refer to the Methods description for details), suggesting that the occurrence of and selection for these mutations was promoted by T492I. All of these T492I-promoted mutations were present in at least one of the Omicron sublineages, most of which (95%) were still dominant around the world (April, 2024, Figure 1F-H and Supplementary Figure S1G). Similarly, T492I appeared to promote synonymous mutations as well as mutations in the noncoding region characteristic for the Omicron variant (Supplementary Figure S1H, I and Supplemental Tables S2, S3). On the basis of the viral quasispecies reconstructed by TenSQR ^30^ (Data S1), the predicted dominant strains in the 492I replicates harbored a greater percentage of dominant-to-date mutations than did the T492 replicates, both for the 90-day and 45-day experiments (Supplementary Figure S1J, K). These findings suggest T492I-induced evolution toward adaptiveness and the emergence of Omicron variants. When the 22 Delta-to-WT reverse mutations were excluded (Supplementary Figure S1L), 54 mutations remained. The fraction of Omicron sublineage mutations in these 54 T492I-promoted mutations was generally greater than the fraction of other high-frequency mutations (Figure 1I), and the fraction of these 54 T492I-promoted mutations in Omicron sublineage mutations was also greater than those of other high frequency mutations (Figure 1J). Similar fraction differences were not found for the VOCs Alpha and Delta. This confirmed the Omicron-biased evolution induced by T492I. The fraction of T492I-promoted mutations in the mutations of the Omicron BA.2 (86%) and BA.5 (85%) was greater than those of other Omicron sublineages, suggesting an evolutionary bias toward early Omicron lineages induced by T492I.

### Omicron-biased selective force induced by T492I

We performed sliding window analyses of selection signatures by ANGSD ^31^ and SweeD ^32^ to evaluate the impacts of T492I on the evolutionary driving force. The results revealed that the evolved populations of the 492I runs had more regions with high nucleotide diversities (π) and deviated-from-zero Tajima’s D than did the T492 runs (Figure 2A, B and Data S1), which was consistent with the increased mutation rate observed in the 492I runs. π and Tajima’s D both positively correlate the frequencies of mutations (Supplementary Figure S2A, B). Accordingly, there was increased genetic differentiation (Fst) between the 492I and T492 runs in the genomic regions with high-frequency mutations (Figure 2C, Data S1). For the 492I runs, the genomic regions with fixed Omicron mutations presented a greater π and a greater deviation from zero for Tajima’s D than did those without fixed Omicron mutations (Figure 2D, E). These results suggest Omicron-biased evolutionary forces induced by T492I. Positive selection signatures, peaks of the composite likelihood ratio (CLR) ^33, 34^, were identified in the evolved populations of the 492I runs (Figure 2F, G and Supplementary Figure S2C, D) but were absent in the evolved populations of the T492 runs, possibly due to the limited number of identified SNPs after removing monomorphic cases. For the 492I runs, mutations characteristic for Omicron sublineages biasedly emerged in genomic regions with evidence for positive selection (a high CLR with a P-value <0.05 according to the ranking tests, Figure 2H). The identification of selection signatures via another population genomics analysis tool (CLEAR ^35^) revealed that, compared with those of the T492 runs, the evolved populations of the 492I runs presented a greater accumulation of Omicron mutations that were positively selected (Figure 2I). The 492I runs also had more Omicron mutations in the genomic regions with estimated positive selection strength than the T492 runs (Figure 2J). The estimated effective population sizes in the 492I runs were smaller than those in the T492 runs (Supplementary Figure S2E). This suggests positive directional selection of Omicron mutations induced by T492I. The 45-day Delta-I runs showed more positions with a positive Tajima’s D and fewer positions with a minus Tajima’s D than the 90-day Delta-I runs (Kolmogorov-Smirnov tests, P-value = 1.371e-05), inferring a transition in the degree of selective sweep from the 45-day state (intermediate) to the 90-day state. Directional selection may have effects similar to those of balancing selection (Tajima’s D>0) if the selected mutations are present at intermediate frequencies ^36^.

**Figure 2.**
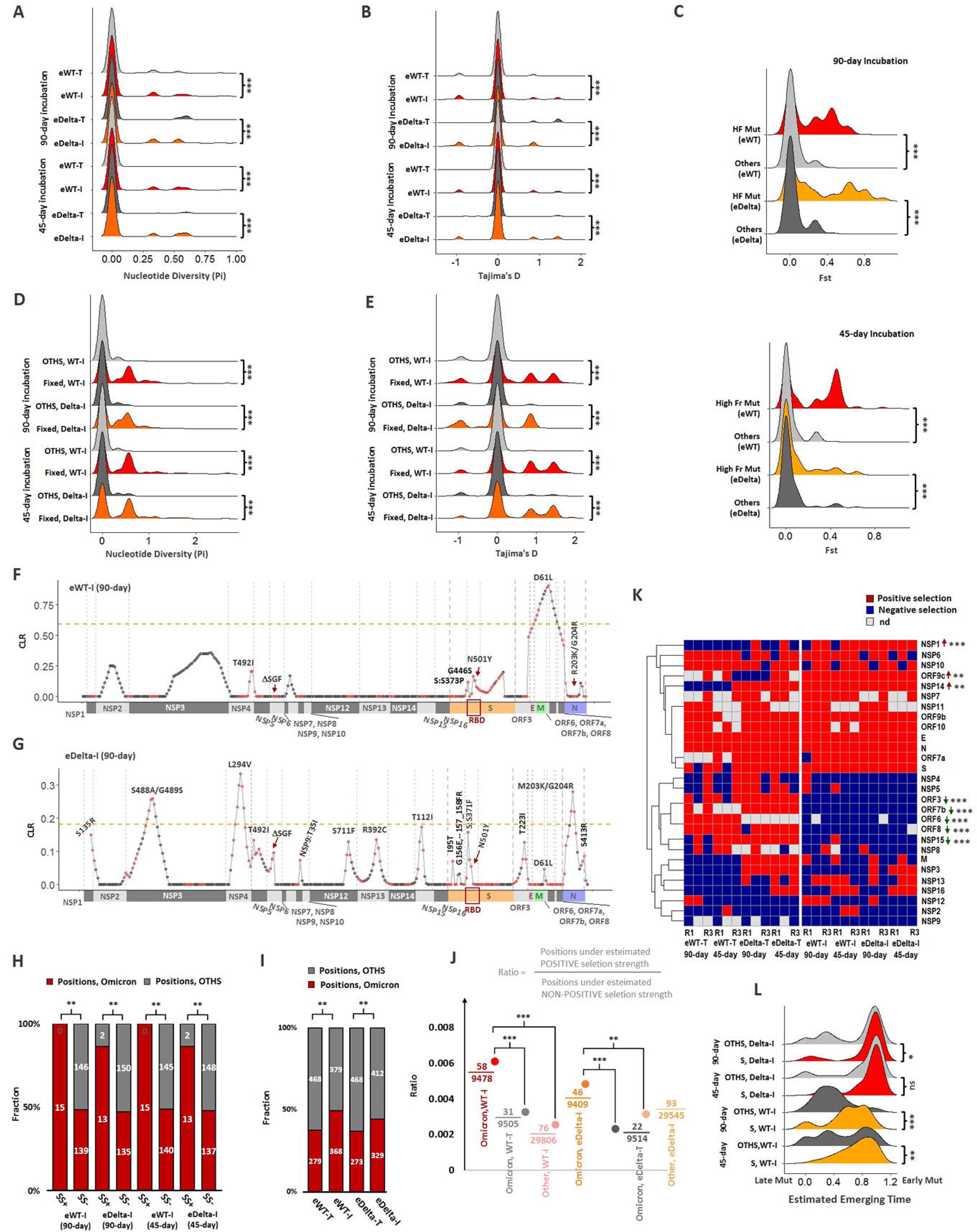
Selective forces induced by T492I. (A-E) show the distributions of selection signatures on the basis of sliding window analyses across runs. (A, B) Comparison of the distributions of the nucleotide diversity (A) and Tajima’s D (B) between the T492 and 492I runs. (C) Comparison of the genetic differentiation (Fst) between the genomic regions with high frequency mutations (HF Mut) and those without high frequency mutations (others), both for the 90-day (upper) and 45-day runs (lower). (D, E) Comparisons of the distributions of the nucleotide diversity (D) and Tajima’s D (E) between the genomic regions with fixed Omicron mutations (Fixed) and those without fixed Omicron mutations (OTHS). For (A-E), comparisons were performed via Kolmogorov-Smirnov tests. (F, G) Sliding window views display the CLR peaks and the thresholds (orange dotted line), for the 90-day eWT-I (F) and eDelta-I runs (G). The points colored in red denote the positions with high frequency mutations. (H) Comparisons of the fractions of Omicron mutations (Positions, Omicron) between the positions with positive selection signatures (SS+) and those without positive selection signatures (SS-). ‘Positions, OTHS’ denotes the positions without Omicron mutations. (I) Comparisons of the fractions of Omicron BA.2 mutations (Positions, Omicron) within the positions with a significantly high likelihood (*H*) across different runs. (J) Comparisons of the ratio of the positions with an estimated positive selection strength and those with an estimated nonpositive selection strength across the positions with Omicron BA.2 mutations in eDelta-I (Omicron, eDelta-I), eDelta-T (Omicron, eDelta-T), eWT-I (Omicron, eWT-I) and eWT-T (Omicron, eWT-T) and those without Omicron mutations in eDelta-I (Other, Delta-I) and eWT-I (Other, WT-I). The statistics in (H-J) were performed via chi-square tests. (K) Estimated selection of the proteins in different runs. ‘nd’ denotes not detectable, due to the lack of segregating sites. By Fisher’s exact test, the proteins with a significantly greater percentage of positive selection cases in the 492I runs than in the T492 runs are marked by upward red arrows, and those with a significantly greater percentrage of negative selection are marked by green downward arrows. (L) Comparisons of the distributions of the estimated emergence times of the mutations in the spike protein (S) and other proteins (OTHS) for the 90-day and 45-day 492I runs (Delta-I and WT-I). ‘Early Mut’ and ‘Late Mut’ denote the mutations that evolved early and those that evolved late in the run, respectively. Statistics were performed via Kolmogorov-Smirnov tests. ‘*’ denotes a P-value < 0.1, ‘**’ denotes a P-value < 0.05, and ‘***’ denotes a P-value <0.01.

Furthermore, we calculated the ratios of the nonsynonymous Pi and synonymous Pi (PiN/PiS) across SARS-CoV-2 proteins. The proteins with a significant >1 PiN/PiS and a significant <1 PiN/PiS were considered positively selected and negatively selected, respectively. The results (Figure 2K) revealed that the proteins NSP1, NSP14 and ORF9c were more positively selected in the 492I runs than in the T492 runs. These three proteins are involved in immune evasion ^37–39^ and the variations observed here suggest viral adaptations to the host innate immune response ^40^. The proteins NSP15, ORF3, ORF6, ORF7b and ORF8 were negatively selected in the 492I runs compared with the T492 runs. Overall, the 492I populations presented more negatively selected proteins and fewer positively selected proteins than the T492 populations (chi-square tests, P-value = 0.001855), suggesting the overall functionality of the T492I-promoted mutations. This was confirmed by a lower Tajima’s D in the 492I strains of the 90-day runs than in the 45-day runs. On the basis of the *in silico* reconstructed viral quasispecies referred to above, we predicted the emergence time of the mutations in the dominant strain. The results revealed that nonspike mutations generally emerged later than spike mutations did (Figure 2L), both in the 90-day eDelta-492I runs and the 45-day eDelta-492I runs. This finding agrees with previous evidence of the highest accumulation of spike mutations among all SARS-CoV-2 proteins ^41, 42^. The 90-day eDelta-I runs harbored more late emerged mutations than the 90-day eWT-I runs did, and similar effects were observed in eDelta-I and eWT-I for the 45-day runs (Supplementary Figure S2F). This may be due to the late emergence of Delta-to-WT reverse mutations (Supplementary Figure S2G).

### T492I induces an evolution toward Omicron-like phenotypes

We predicted phenotypic changes induced by T492I-promoted mutations using published experimental data ^4, 43, 44^. Based on experimental infection results for normalized pseudo particles and CaCo-2 cells ^43^, 11 out of 28 T492I-promoted spike mutations were found to promote infection (Supplementary Figure S2H). Most T492I-promoted spike mutations show a promoting effect on the expression and processing of the spike proteins, although no significant correlation was found between the frequencies of mutations in the 492I populations and the distances of mutations to the receptor binding domain (Supplementary Figure S2J). The T492I-promoted spike mutations are mostly associated with increases in the levels of serum neutralization capability (Supplementary Figure S2H), and show potentials to induce higher cross-species infectivity than other mutations (Supplementary Figure S2I). The T492I-promoted non-spike mutations N R203K/G204R and NSP6 ΔSGF were likewise associated with enhancement in viral replication, infectivity and immune evasion capability, according to previous records ^19, 27, 45, 46^. These findings suggest potential Omicron-like phenotypes induced by the T492I-promoted mutations.

Following the prediction results described above, we tested and compared the replication and infectivity of ancestral populations and the populations resulting from our 90-day evolution trial in human lung epithelial cells (Calu-3, Figure 3A-I). The results show that eWT-I and eDelta-I (the evolved populations from the 492I runs) produced higher levels of extracellular viral RNA at 24 and 36 hours post infection (hpi) than eWT-T and eDelta-T (the evolved populations from the T492 runs), respectively (Figure 3B). This was similarly true for ancestral populations when compared among each other (aWT-I vs aWT-T and aDelta-I vs aDelta-T, Figure 3A). By calculating the fold changes of detected viral genome copies before and after evolution, we found that the T492I mutation endowed both WT and Delta strains with stronger replication capabilities after 30 passages (Figure 3C). Further, viral infectivity was measured by plaque assay and viral subgenomic RNA assay. The results showed that the evolved populations from the 492I run produced significantly higher infectious titers (Figure 3D, E) and E sgRNA loads (Figure 3G, H) compared to those of the control (the T492 run). We also found that the mutation T492I conferred increased infectivity to both the WT and Delta strains after the evolution process (Figure 3F, I).

**Figure 3.**
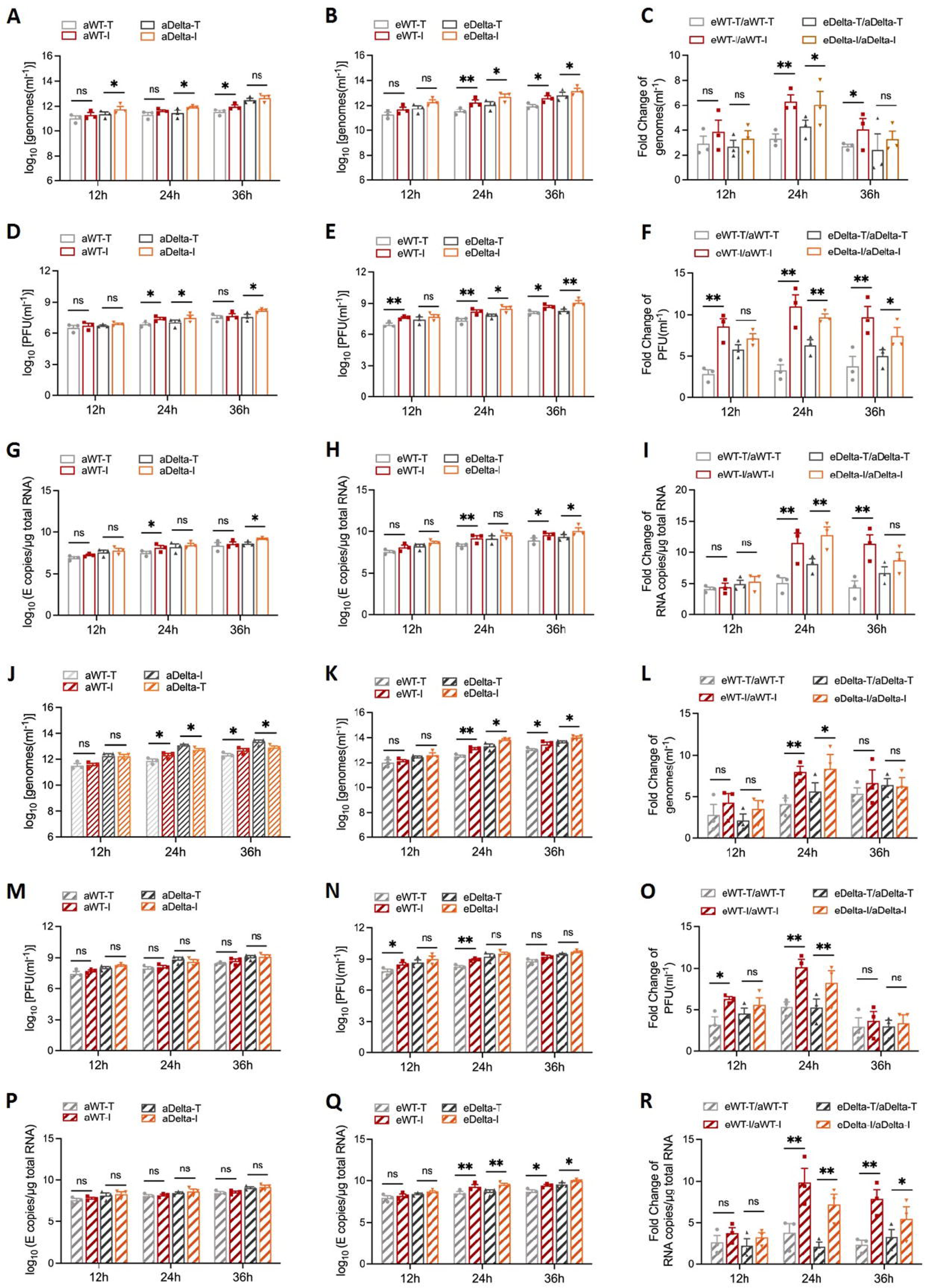
Evidence suggesting enhanced increases of viral replication and infectivity in 492I populations. (A-I) Calu-3 cells were infected with aWT-T, aWT-I, aDelta-I or aDelta-T virus at a MOI of 0.01. Genomic RNA levels (A, B), PFU titres (D, E), and E sgRNA loads (G, H) were detected via plaque assays and qRT-PCR. Fold changes in genomic RNA levels (C), PFU titres (F), and E sgRNA loads (I) were calculated from the ratios of evolved populations to ancestor strains. The experiments were performed in triplicate. (J-R) Genomic RNA levels (J, K), PFU titres (M, N), and E sgRNA loads (P, Q) were evaluated in the Vero E6 cell line. Fold changes (L, O, R) were calculated from the ratios of evolved populations to ancestor strains. The experiments were performed in triplicate. ‘*’ denotes a P-value < 0.1, ‘**’ denotes a P-value < 0.05, and ‘ns’ denotes ‘not significant’.

Furthermore, replication and infectivity tests were performed on the Vero E6 cells. In this experiment, similar trends were observed for fold changes in extracellular viral RNA (Figure 3J-L), infectious titers (Figure 3M-O), and E sgRNA loads (Figure 3P-R) before and after evolution. In particular, there are no significant differences in PFU titers (Figure 3M) and E sgRNA loads (Figure 3P) between either aWT-I and aWT-T or aDelta-I and aDelta-T. This is consistent with our previous observations ^18^. However, eWT-I produced significantly higher PFU titers (Figure 3N) and E sgRNA loads (Figure 3Q) than eWT-T, and so did eDelta-I compared to eDelta-T. These results demonstrate an enhancement of replication and infectivity induced by T492I in the evolution of SARS-CoV-2.

A previous study from our lab suggested a potential association between the T492I mutation and immune evasion capacity of SARS-CoV-2 ^18^. We subsequently tested the production of interferon (IFN)-β, IFN-λ, and interferon stimulated gene 56 (ISG56) in the pre– and post-evolution variants. As a result, the T492I mutation showed inhibitory effects on the mRNA level of IFN-β in both pre– and post-evolution wild type group (Figure 4A, B), but the inhibitory effect was stronger when the T492I mutation strains evolved over time (Figure 4C). Although there were no significant differences in the IFN-β level between T492 and 492I virus in the pre-evolution Delta group (Figure 4A, B), the production of IFN-β was significantly higher in the 492I virus of post-evolution Delta group (Figure 4C), suggesting that the WT and Delta strains bearing T492I acquired enhanced immune evasion capabilities in the 90-day evolution. These were further confirmed by examining the production IFN-λ and ISG56 at 24 and 36 hpi in Calu-3 cells (Figure 4D-I). Omicron mutations enhance the viral replication, infectivity and immune evasion capability of SARS-CoV-2 ^47–49^. The identified enlarged phenotypic alteration of the evolved populations in the 492I runs confirmed the Omicron-biased evolution induced by T492I.

**Figure 4.**
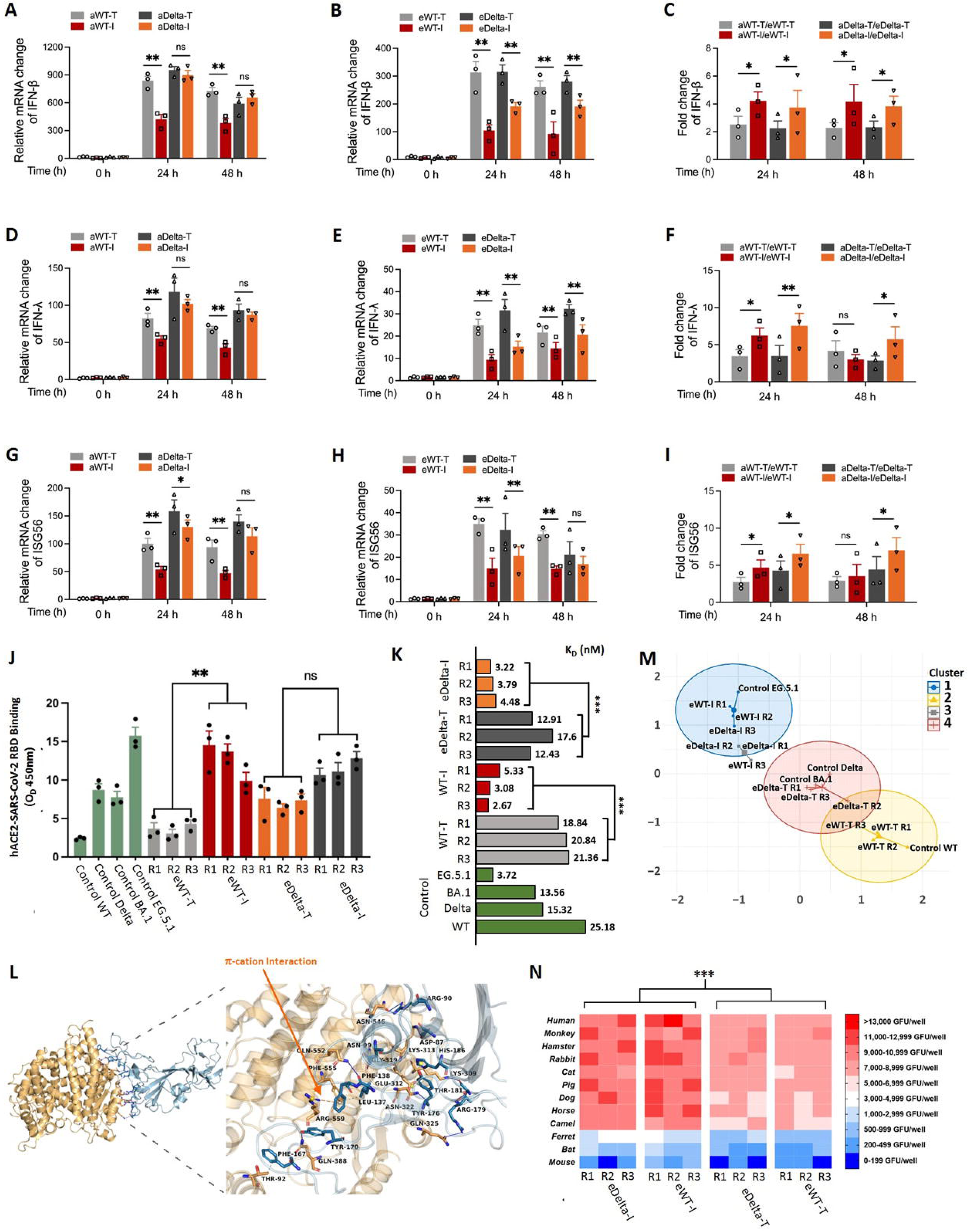
Evidence suggesting enhanced increases of immune evasion capability, RBD-hACE2 binding affinity and cross-species infection potentials after T492I-influenced evolution. (A-I) Calu-3 cells were infected with aWT-T, aWT-I, aDelta-I or aDelta-T virus at an MOI of 0.01. At 12, 24, and 48 h after infection, total RNA extracted from the cells was evaluated via real-time qRT-PCR. The relative changes in the IFN-β (A, B), IFN-λ (D, E), and ISG56 (G, H) mRNA levels were normalized to that of the GAPDH mRNA. Fold changes in the IFN-β (C), IFN-λ (F), and ISG56 (I) mRNA levels were calculated from the ratios of evolved populations to ancestor strains. The experiments were performed in triplicate. (J) The binding ability of the RBD and hACE2 was detected via ELISA. The RBD recombinant proteins were added to hACE2-coated plates, the mixture was incubated at room temperature for 1 h, and the absorbance was read at 405 nm. (K) The values of K_D_ in different groups resulting from the SPR experiments. Statistics was performed via ANOVA tests. (L) Representative molecular docking results of the hACE2/evolved RBD complex. The orange dotted line indicates the π-cation interaction that formed by ARG-559 of hACE2 and PHE-138 of the eWT-I R1 RBD. hACE2, orange; RBD, blue; hydrophobic interaction, gray dotted line; hydrogen bond, blue line; π-stacking (parallel), green dotted line; π-cation interaction, orange dotted line; salt bridge, yellow dotted line. (M) Results of phylogenetic-based clustering analyses based on the O_D_ and K_D_ values from the Enzyme-Linked Immunosorbent Assay (ELLSA) and SPR experiments. (N) The spike-expressing plasmid of evolved viruses, the packing plasmid, and the mNeonGreen reporter vector were cotransfected into HEK-293T cells to generate spike-bearing LVpps. The infection performance of LVpps bearing the spike of evolved viruses in Calu-3 cell lines stably expressing ACE2 orthologs (human, monkey, hamster, rabbit, cat, pig, dog, horse, camel, ferret, bat and mouse) is shown as a heatmap. Statistics was performed via Wilcoxon tests. ‘*’ denotes a P-value < 0.1, ‘**’ denotes a P-value < 0.05, and ‘ns’ denotes ‘not significant’.

### Increased RBD-hACE2 binding affinity and cross-species infection potentials induced by T492I

To further investigate the infection capacity of the evolved viruses, we tested the binding affinity of the evolved populations to human angiotensin-converting enzyme 2 (hACE2). According to the viral quasispecies reconstructed from our sequencing data, the predicted predominant strains of all replicates (R1, R2 or R3) were selected as representative strains (Supplemental Table S4), and their receptor-binding domains (RBDs) ^50^ were synthesized and subjected to a hACE2 binding ELISAs (Enzyme-Linked Immunosorbent Assay). The ancestor strains, WT and Delta, and the epidemic strains, BA.1 and EG.5.1, were used as controls. The results show that eWT-I has a stronger binding ability to hACE2 than eWT-T, and eDelta-I did not significantly increase the hACE2 affinity after 30 passages, compared with eDelta-T (Figure 4J). This data implies a potential association between the T492I-induced evolution and hACE2 affinity of the spike RBD. Next, we quantitatively analyzed the hACE2 affinities of these strains through surface plasmon resonance (SPR) experiments. Consistent with the results obtained from ELISA, strains of the 492I runs had significantly stronger hACE2 affinities than their controls (Figure 4K and Supplementary Figure S3). Through molecular docking, we found that the RBD-hACE2 binding interface of the eWT-I strains formed a noncovalent π-cation interaction, thereby further improving receptor affinity (Figure 4L and Supplementary Figure S4A-J and Supplemental Table S5). These results provide structural evidence to support the conduciveness of the T492I mutation to strengthening the affinity between RBD and hACE2 during the evolution. Based on the O_D_ (optical density) and K_D_ (dissociation rate constant) from ELISA and SPR experiments, phylogenetic-based clustering analyses show that the evolved populations of eWT-I and eDelta-I are clustered with EG.5.1, an Omicron sublineage predominant in 2023 (Figure 4M). These findings also confirm the T492I-induced Omicron-bias.

For validation of the predicted cross-species infectivity induced by T492I, we generated 12 stable Calu-3 cell lines with exogenous expression of various ACE2 orthologs and infected them with different lentiviral-pseudotyped particles (LVpp) bearing the RBDs of the reconstructed dominant strains to determine the host tropism of SARS-CoV-2 variants. By analyzing the infection performance, we found that most ACE2 orthologs could efficiently support virus entry, except those expressing the ACE2 of ferret, bat and mouse. Interestingly, the mutation T492I conferred drastically increased cross-species infectivity to the evolved viruses (Figure 4N), suggesting an enhanced cross-species infectivity induced by the T492I mutation.

### T492I induces biased mutation types by regulating RNA-editing enzymes

The emergence of Omicron is associated with shifts in the relative rate of mutations ^51^. Following up on this, we tried to evaluate the impact of T492I on the mutation type through comparative analyses of the single base substitution (SBS) spectrum in the evolved populations of different runs. The results show higher relative rates of A>T, T>G, C>A, C>G and G>A and lower relative rates of A>C, T>C and G>T in the 492I runs than the T492 runs (Figure 5A). The APOBEC enzymes presumably induce C>T and G>A if APOBEC deaminates cytosine in the antigenome ^52–56^. The ADAR enzyme induces the A>G transition and the T>C transition in the antigenome ^57^. Thus, the biased relative rates of G>A, T>C and G>T may be a result of the activation of APOBEC and inactivation of ADAR possibly induced by T492I. The higher frequencies of the nucleotide G near the 3’ end of the G>A mutations (Supplementary Figure S4K, L) suggest that the potential major target sequences of APOBEC are 5’-CC-3’ and 5’-GG-3’ in the antigenome. APOBEC3G prefers the target sequence 5’-CC-3’ ^58, 59^ and probably function predominantly in enhancing the mutation from G to A. We therefore examined the relative expression of APOBEC3A, APOBEC1, APOBEC3G and ADAR in the Calu-3 cells infected by the virus with or without T492I. The results show that infection with the T492I and Delta variants induced significant upregulation of APOBEC3A, APOBEC1 and APOBEC3G, while infection with WT and I492T variants resulted in remarkable induction of ADAR (Figure 5B-F). Similar trends were observed when using Vero-E6 cells (Figure 5G-K). Collectively, these experiments validated the capability of the NSP4 T492I to bias the mutation types by regulating APOBEC and ADAR (Figure 5L).

**Figure 5.**
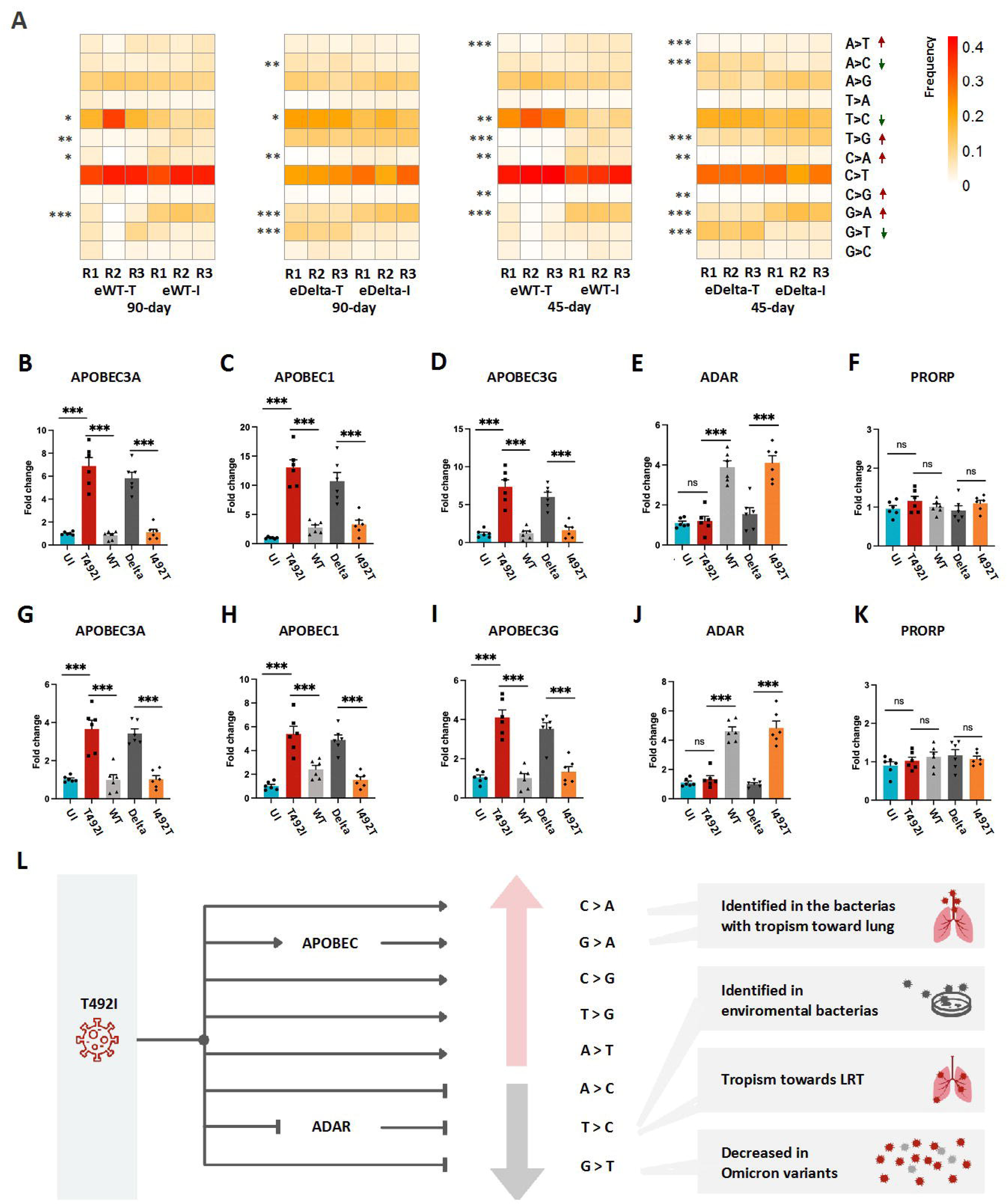
Evidence suggesting biased mutation types, regulation of mRNA editing enzymes and epistatic effects induced by T492I. (A) Heatmaps display the frequencies of different mutation types in different evolved populations. Statistics was performed via ANOVA tests for comparisons between the 492I and T492 replicates in eWT and eDelta both for 90-day and 45-day runs. Consistent or near consistent (without contradictory trends) differences are marked by red upward arrows (increase in frequency in >1 comparisons) and green downward arrows (decrease in frequency in >1 comparisons). (B-K) Calu-3 cells (B-F) or Vero E6 cells (G-K) were infected with the WT, T492I, Delta or Delta-I492T virus at a MOI of 0.01. At 24 h after infection, total RNA extracted from the cells was evaluated by real time qRT-PCR. The relative changes in the APOBEC3A (B, G), APOBEC1 (C, H), APOBEC3G (D, I) ADAR (E, J) and PRORP (Control, F, K) mRNA levels were normalized to the GAPDH mRNA level. The experiments were performed in triplicate. (L) Possible mechanism for the impacts of T492I on mutation types by regulating the expression of the APOBEC and ADAR deaminases. ‘*’ denotes P-value < 0.1, ‘**’ denotes P-value < 0.05, and ‘ns’ denotes ‘not significant’.

**Figure 6.**
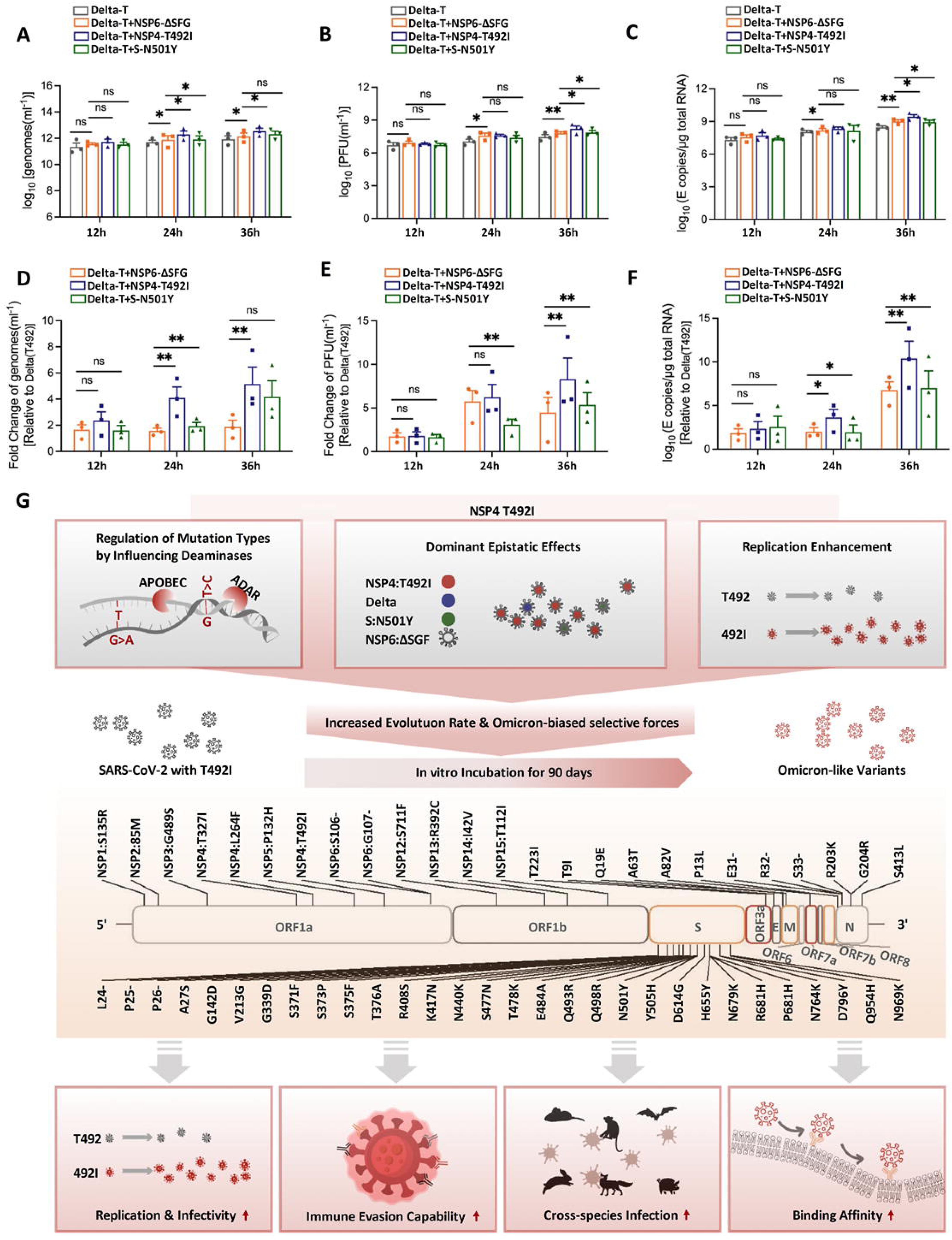
Evidence suggesting strong epistatic effects of T492I. Calu-3 cells were infected with Delta-T variants bearing the NSP4 T492I, S N501Y and NSP6 ΔSFG mutants at an MOI of 0.01. The genomic RNA levels (A), PFU titres (B), and E sgRNA loads (C) were detected via plaque assays and qRT-PCR. Fold changes in the genomic RNA levels (D), PFU titres (E), and E sgRNA loads (F) were calculated from the ratios of the NSP4 T492I, S N501Y and NSP6 ΔSFG mutants to the Delta-T strain. The experiments were performed in triplicate. (G) Diagram displays a proposed mechanism of the Omicron-biased evolution induced by T492I. ‘*’ denotes P-value < 0.1, ‘**’ denotes P-value < 0.05, and ‘ns’ denotes ‘not significant’.

### T492I confers a predominant epistatic effect

Other genomic variations, such as the S mutation N501Y and the NSP6 deletion ΔSGF, are reported to play essential roles in the increased infectivity and transmissibility of SARS-CoV-2 ^17, 23^. N501Y also has a predominant epistatic effect in S mutations ^23^. In order to further validate the advantage of the T492I mutation in the evolution of the virus, we constructed NSP4 T492I, NSP6 ΔSGF and S N501Y mutants based on the Delta 21A strain (bearing T492), respectively, and then compared the replication and infectivity of these variants in Calu-3 cells. The results showed that the T492I mutation produced significantly higher extracellular viral RNA (Figure 6A), infectious titers (Figure 6B) and E sgRNA loads (Figure 6C) than the S N501Y and NSP6 ΔSGF variants. Through normalizing the data to aDelta-T, we found that the T492I mutation contributed more to viral replication and infectivity than S N501Y and NSP6 ΔSGF (Figure 6D, E, F). These results collectively suggest a stronger epistasis by NSP4 T492I than S N501Y and NSP6 ΔSGF, and support a predominant epistatic effect of T492I. The epitasis of T492I could play a key role in the induced Omicron-biased evolution (Figure 6G).

## DISCUSSION

Our study demonstrated that the NSP4 mutation T492I is capable of inducing rapid evolution toward the emergence of Omicron-like variants via 24 independently performed replicates of evolve-and-resequence experiments for wild-type and Delta strains over incubation periods of 45 and 90 days (Figure 6G). Our evidence includes an elevated mutation rate, increased proportions of high-frequency mutations, a high ratio of Omicron/fixed mutations in T492I-promoted mutations and high coverage of T492I-promoted mutations in the mutations of historically early Omicron variants, such as BA.2 and BA.5. The acceleration of evolution toward Omicron was also confirmed by the increase in viral replication, infection, immune evasion capability and cross-species infection potentials of the viral populations that evolved from ancestors with T492I compared with those from ancestors without T492I. Consistent with previous findings in Omicron early variants, T492I-promoted mutations impair the infectivity of SARS-CoV-2 (e.g. S375F); however, the mutations that evolved in the 492I strains together show an infectivity-promoting effect (Figure 3). The T492I-promoted mutations in the spike protein also induce an accelerated increase in the RBD-hACE2 binding affinity. We concluded that the NSP4 mutation T492I is a key contributor to the emergence and spread of Omicron mutations.

Previous work revealed that the N501Y mutation is the predominant spike mutation that epistatically enables other affinity-enhancing mutations ^23^. It has also been suggested that mutations in the spike protein and NSP6 determine the function of Omicron variants ^17^. We provide experimental evidence that the NSP4 mutation T492I has a stronger epistatically enhancing effect on infectivity and transmission than the other adaptive mutations tested, spike N501Y and NSP6 ΔSGF. These findings, together with the high overlap between the Omicron mutations and T492I-promoting mutations, suggest that the NSP4 mutation T492I causes considerable epistatic shifts in the effects of mutations at other sites. The results obtained from our evolutionary analysis suggest that the epistasis of T492I alters evolution forces and causes positive selection toward mutation sites characteristic for the Omicron variant. Additionally, T492I regulates the expression of mutation-inducing host proteins, possibly resulting in biased mutation types toward the emergence of Omicron mutations. In conclusion, we found that T492I can regulate the evolution of SARS-CoV-2 by influencing mutation-inducing proteins, exerting strong epistatic effects and enhancing virus replication (Figure 6G).

The 492I runs showed a 2∼5 fold higher mutation rate than the T492 runs. The median mutation rate in eWT was 0.000027 nt^-1^ T^-1^, which is consistent with the mutation rate underlying the global diversity of SARS-CoV-2 (1 × 10^−5^ ∼ 1×10^−4^ nt^-1^ T^-1^)^60, 61^. The global historical statistics showed that the 492I variants had a lower number of mutations than the T492 variants in the spread of the VOC Alpha (Fig S1B), possibly due to the exclusion of T492I in the Alpha variants. The regulation of mutation-inducing proteins, such as the upregulation of APOBEC and the downregulation of ADAR, should promote the emergence of Omicron-biased nucleotide substitutions, such as G>A and T>C. Epistasis increases the adaptiveness of Omicron mutations and thus accelerates the increase in the frequency of the emergent Omicron mutations. Similarly, previous work revealed that C>T (the reverse-complement of G>A) and C>A prefer to occur in lung bacteria and that T>C prefers to occur in environmental bacteria ^62^. Moreover, G>T mutations seem to be generally elevated when viral replication occurs in the lower respiratory tract (LRT) ^63^ and G>T mutations were previously found to be decreased in Omicron clades ^51^. Consequently, the biases of G>A, C>A, T>C and G>T induced by T492I infer a tropism toward the upper respiration tract (URT), which agrees with the identified high ratio of Omicron mutations induced by T492I and the tropism toward URT of the Omicron variants ^12, 64^ (Figure 5L). Reactive oxygen species (ROS) are proposed to be relevant to the G >T transition ^65, 66^. The reduction in G>T changes possibly resulted from the impairment of the activity of ROS by T492I. Increased viral replication also accelerates the increase in the frequency of Omicron mutations. The strong epistatic effects of T492I may be a result of the effects of T492I on the regulation of other SARS-CoV-2 proteins, as we reported previously ^18^.

Our experiments demonstrated the *in vitro* emergence of Omicron-like variants from the ancestors with T492I within 90 days. This is in accordance with global epidemiological statistics that show the emergence of Omicron (December 2021) occurred three months after the global identified sample frequency (IF) of T492I reached 80% (August 2021). The rapid emergence of Omicron mutations in the 492I strains may partly explain the sudden emergence and rapid spread of Omicron variants at the end of 2021. Our results show that eDelta-I has a greater fraction of high frequency mutations than eWT-I does, both for the 45-day and the 90-day runs. This possibly results from the effects of replication-enhancing Delta mutations, such as S D614G ^20, 67^ and N R203M ^68^. However, the biases toward high-frequency mutations in eDelta-I do not support the hypothesis that Omicron variants evolved from Delta ancestors. Compared with the WT-to-Omicron process, nearly twenty Delta mutations need to be reversely mutated for a Delta-to-Omicron evolution process. Moreover, reverse mutations mostly emerged later than other Delta-to-Omicron mutations did, and few strains with an intermediate Delta-to-Omicron state were identified. Previous work revealed that the Omicron variants originated from early SARS-CoV-2 variants ^69, 70^ and that the nearest outgroup could have been a progeny from the B.1.1.519 (20B) lineage ^71^. According to the data provided by Nextstrain ^72^, the B.1.1.519 variant also harbors S 614 and N 203 mutations. Together with already available replication-enhancing mutations, the emergence of NSP4 T492I in the strains of B.1.1.519 may have induced a fast evolution toward the emergence of Omicron variants. The impact of T492I on the historical evolution progress of SARS-CoV-2 is surprising and is similar to a wormhole-tunnel-like effect that has shortened the evolution time from one state to another state. In Astrocosmology, the wormhole tunnel refers to a hypothetical structure connecting disparate points in spacetime in theoretical physics ^73^. Considering the possibility of missing historical samples ^15^, further work is still needed to identify the origins of Omicron.

## LIMITATIONS OF STUDY

We did not perform animal experiments, such as the Syrian golden hamsters, to evaluate and compare the viral replication, infectivity, immune evasion capability, virulence of virus and tropism for the evolved populations *in vivo*. Moreover, the effects of T492I on the expression of APOBEC and ADARp, the epistatic effects of T492I on other identified adaptive mutations and the effects of T492I on SARS-CoV-2 in evolution are needed to be evaluated by animal experiments in the future.

## ACKNOWLEDGMENTS

We gratefully acknowledge the submitting and the originating laboratories where genetic sequence data were generated and shared via NCBI and the GISAID Initiative. This work was supported by grants from the National Natural Science Foundation of China (92369115, 32170661), the SGC’s Rapid Response Funding for COVID-19 (C-0002), and the Fundamental Research Funds for the Central Universities (2023CDJXY-009). The funders had no role in study design, data collection and analysis, decision to publish, or preparation of the manuscript.

## AUTHOR CONTRIBUTIONS

Z.Z., C.Z., M.T. and C.J. performed the *in silico* analyses. H.W., J.T., Z.S., J.A., R.V., C.L., B.F., Y.X., and D.K performed the experiments. X.L., H.W., H.Z., X.Z., Q.L., J.Y., X.X.L., S.W and X.M. performed the protein structure analysis. Z.Z. conceived the idea. Z.Z., H.W., J.T. and Q.Z. managed the project. Z.Z., Q.Z., H.W., J.T. and X.L. wrote the manuscript and coordinated the project.

## METHODS

### Cell culture and infection

The key resources used in this study is listed in Table 1. Human lung epithelial Calu-3 cells (HTB-55, ATCC, MD, USA) and African green monkey kidney epithelial Vero E6 cells (CRL-1586, ATCC) were maintained at 37 °C with 5% CO2 in high-glucose Dulbecco’s modified Eagle’s medium (DMEM, Gibco, CA, USA) supplemented with 10% FBS (Gibco). The wild-type (USA_WA1/2020 SARS-CoV-2 sequence, GenBank accession No. MT020880) and Delta B.1.617.2 (EVAg: 009V-04187, European Virus Archive, www.european-virus-archive.com) SARS-CoV-2 viruses were generated via using a reverse genetic method as previously described ^18, 19^. NSP4 T492I mutation or I492T reverse mutation was introduced into the backbone virus via overlap-extension PCR as previously described ^18^. Briefly, a large bacterial artificial chromosome (BAC) construct established in our laboratory was used as a template, and the recovery of the viral mutants was performed by transfecting BAC DNA from the mutants into Vero-E6 cells with Lipofectamine 3000 (Thermo Fisher Scientific). The cells were infected at a multiplicity of infection (MOI) of 0.01 at the indicated time points. All SARS-CoV-2 live virus infection experiments were performed under biosafety conditions in the BSL-3 facility at the Institut für Virologie, Freie Universität Berlin, Germany in compliance with relevant institutional, national, and international guidelines.

**Table 1.**
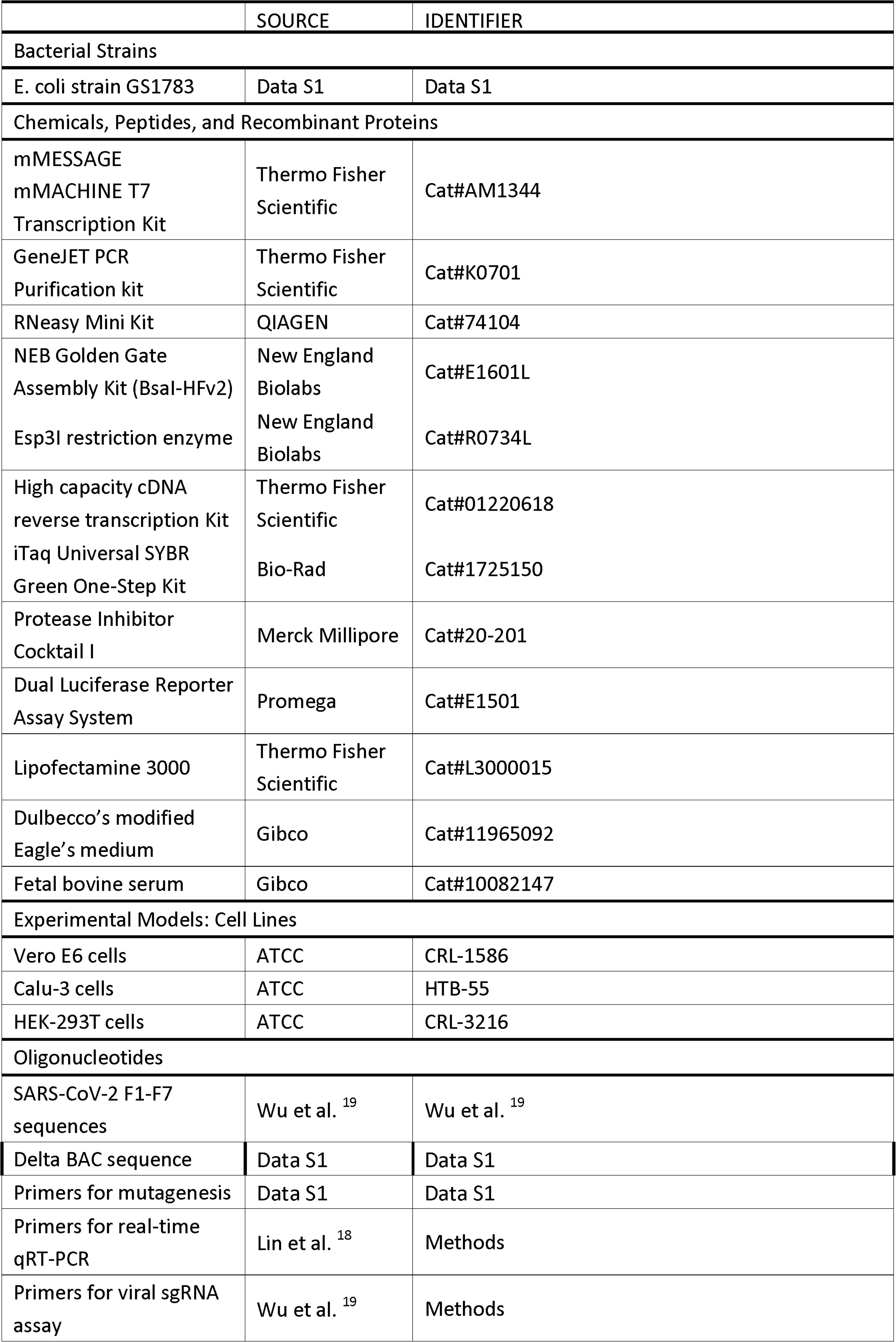
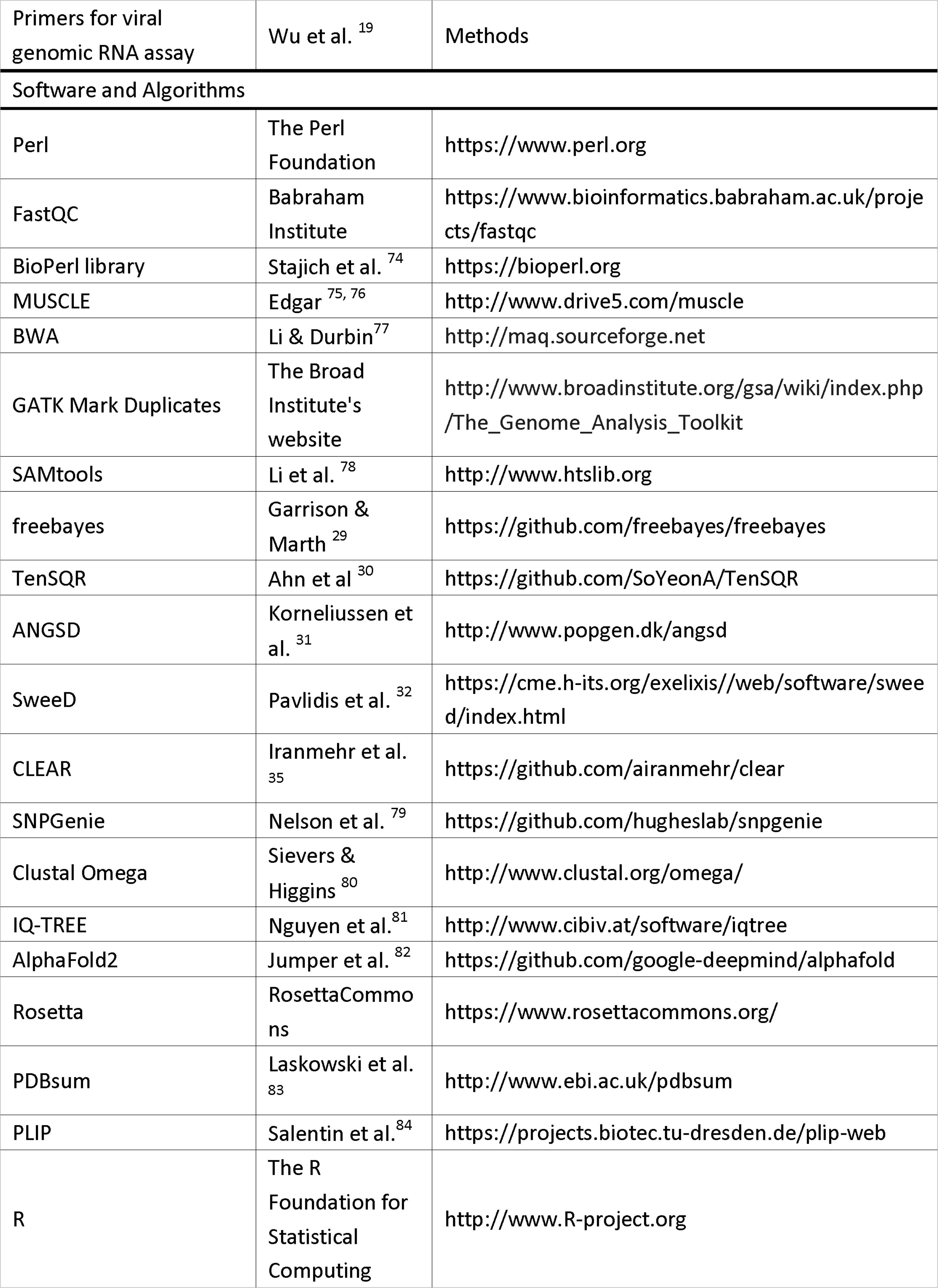
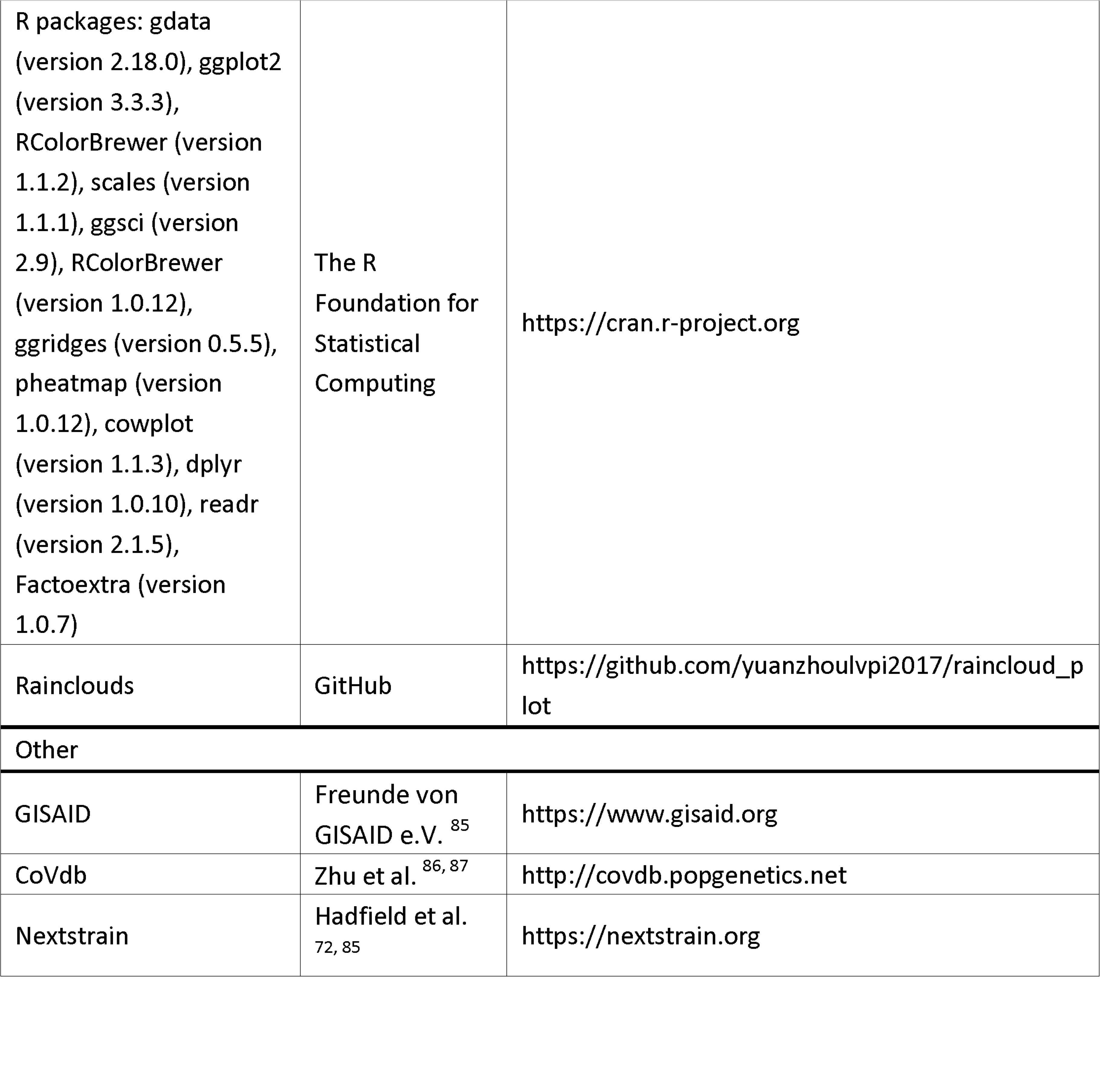
The key resources used in this study.

### Plaque assay

Approximately 5×10^5^ cells were seeded into each well of 12-well plates and cultured at 37 °C under 5% CO_2_ for 12 h. eWT-T, eWT-I, eDelta-T, and eDelta-I viruses were serially diluted in DMEM with 2% FBS, and 100 μL aliquots were transferred to cell monolayers. After 1 h at 37 °C and 5% CO2, the inoculum was removed, and the cells were overlaid with 2X Eaglés minimun essential medium (EMEM; Lonza™ BioWittaker™) containing 1.5% microcrystalline cellulose and carboxymethyl cellulose sodium (Vivapur 611p; JRS Pharma) or MEM containing 1.5% carboxymethyl cellulose sodium (Sigma Aldrich). Forty-eight hours after infection the plates were washed with 1X PBS, fixed with 4% PBS-buffered formaldehyde and stained with 0.75% crystal violet. Visualization of the plaques was performed using a light box.

### Generation of SARS-CoV-2 mutants

The genome of the SARS-CoV-2 WT/Delta strain was cloned into a bacterial artificial chromosome-yeast artificial chromosome (BAC-YAC) using TAR cloning in yeast, as previously described ^88, 89^. Subsequently, the nsp4 mutation T492I/I492T was then introduced into cloned virus genomes by scarless mutagenesis ^90^, using the primers listed in Data S1. The correct introduction of the mutation was verified by commercial nanopore sequencing (Eurofins Genomics).

To recover the WT or mutant viruses, BAC DNA was isolated from E. coli using the Plasmid Midi Kit (Qiagen) and transfected into Vero E6 cells with Lipofectamine 3000 (Thermo Fisher Scientific) as previously described ^91^.

### Sequencing, identification of mutations and relevant analyses

The virus in the wells of each run was extracted. cDNA synthesis and whole-genome amplification were subsequently performed. Pair-end genomic sequencing was performed via an Illumina NextSeq 2000 apparatus and sequence read quality analyses were performed via FastQC (www.bioinformatics.babraham.ac.uk/projects/fastqc/). Quality improvement was performed via FastP ^92^. We mapped the reads of eWT-I and eWT-T to the ancestor genome (GenBank accession No. MT020880 ^87^) via the Burrows-Wheeler Aligner (BWA) ^77^ and marked duplicated reads via GATK Mark Duplicates ^93, 94^. Similarly, we mapped the reads of eDelta-I and eDelta-T to the ancestor genome (EVAg: 009V-04187). The depths of coverage throughout the genome for these samples were mostly near 8000, both for the 90-day and 45-day runs (Figure S5, Data S1). The change in format from SAM to BAM was performed via SAMtools ^78^. We performed mutation calling on the BAM files via freebayes ^29^. The parameters were “-p 1 –C 1 –F 0.01 –-pooled-continuous”. We counted the frequency of mutations in different runs on the basis of output VCF files and wrote Perl scripts to perform the format change and annotation, such as classifying mutation types and attributing nucleotide mutations to corresponding amino acid mutations. The Perl module BioPerl ^74^ was used to translate nucleotides into amino acids. Comprehensive manual checks were also performed in the annotation work. For the difference between the ancestor Delta and WT, we identified Delta-specific reverse mutations (e.g. NSP13 L77P in eDelta) and WT-specific mutations (e.g. S D614G). The premutation state of NSP13 L77P in eDelta is not available in the ancestor of eWT. The postmutation state of S D614G is already available in the ancestor of eDelta. Moreover, convergent mutations were identified, whose postmutation state was shared by eDelta and eWT, but premutation states were different between eDelta and eWT (e.g. N M203K in eDelta and N R203K in eWT, Figure 1H).

For the identification of T492I-promoted mutations, we performed ANOVA tests of the frequencies for the high frequency mutations (with a >0.5 frequency in one or more replicates) between the 492I and T492 runs. The cases with a significant higher frequency in 492I runs than in T492 runs were considered T492I-promoted mutations. We evaluated the historical IFs of the T492I-promoted mutations on the basis of the global genomic epidemiology data of SARS-CoV-2 provided by NextStrain ^72^. The Omicron mutations with a > 0.9 identified IF on April 2024 were considered as fixed Omicron mutations. Using the VOC information provided by NextStrain and the genomic sequences provided by GISAID ^85^ (Table S6), we built a dataset of mutations characteristic for the Alpha, Delta and Omicron sublineages (VOC mutations). According to the dataset, we evaluated the coverage of T492I-promoted mutations in VOC mutations and the coverage of VOC mutations in T492I-promoted mutations. To evaluate the dominant strain in different evolved populations, we used TenSQR ^30^ to reconstruct viral quasispecies from the sequencing data, and then curated the output from the results from freebayes. Based on these, we evaluated the fractions of fixed Omicron mutations in the reconstructed dominant strains of different populations.

For the statistics of the mutation counts in the strains with T492 and those with 492I, we downloaded the protein sequences of 9836814 SARS-CoV-2 samples from GISAID. The collection dates of these samples ranged from December 2019 to March 2022. Following the pipeline we previously used to perform epidemiological analyses ^18, 19, 21^, we performed alignments of proteins between these SARS-CoV-2 strains and the wild-type reference (MT020880) via MUSCLE ^75, 76^, based on the global SARS-CoV-2 genomic data. Using the alignments and information of VOC from NextStrain ^72^, we identified mutations and calculated the IFs of mutations and VOCs (Figure S1B). To evaluate the association between T492I and the mutation rate, we compared the number of mutations (amino acid substitutions) identified early in the global pandemic of SARS-CoV-2 (from April 2020 to November 2020, Figure S1B) between the T492 and 492I strains. The selection of this time course is to avoid the effect from contemporary VOCs. For example, the VOC Alpha began to spread throughout the world after November 2020 (Figure S1E).

### Evolutionary analyses

Based on previous efforts ^42^, we counted the evolution rate (*r*) according the formula as shown below.

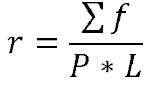

Here *f* is the frequency of some mutation and ∑*f* is the sum of the freqeuncies of all mutations in the genomic region, *P* is the number of transmission events, and *L* is the length of the genome. On the basis of the identified mutations after our evolve-and-resequence experiments, we performed statistics on the ratios of all possible nucleotide mutation types and evaluated the bias between the T492 and 492I runs via ANOVA tests. For the detection of evolutionary signatures,we used ANGSD ^31^ to perform sliding window calculations of the genetic differentiation between the T492 and 492I runs. We also calculated the nucleotide diversity (π) and the values of Tajima’s D for all runs. The window size was 50 and the step size was 20. We piled the BAM files of all runs and performed sliding window analyses of the composite likelihood ratio (CLR) with a grid size of 300 via SweeD ^32^. Furthermore, we utilized the CLEAR tool ^35^ to estimate the population size, selection strength, and likelihood (*H*). In the calculation by CLEAR, we assumed that the evolved populations of the 45-day runs were the midway of those of the 90-day runs. Considering that the ancestral virus of each run was ∼10^6^ cells/ml in 500 µl and that the MOI was 0.01, we used 10^6^*0.5*0.01=5000 as the initial viral population size and constructed the configure files for model parameter estimation.

For the identification of the positions (windows) with mutations characteristic for Omicron sublineages, we counted the number of Omicron mutations in the vicinity (<100 bp) of the middle of a window. If one or more Omicron mutations were identified, the position was considered to have Omicron mutations. In this way, we estimated the associations between Omicron mutations and selection parameters, such as *H*, CLR and the selection strength. To evaluate the selection force on the SARS-CoV-2 proteins in different populations, we used the tool SNPGenie ^79^ to infer the nonsynonymous Pi and synonymous Pi (PiN/PiS) of the proteins.

To estimate the emergence times of the mutations in the dominant strains reconstructed, we performed multiple alignments of all reconstructed strains via Clustal Omega ^80^. Then, maximum-likelihood trees were built via the GTR model and approximate Bayes tests ^95^ were performed via IQ-TREE ^81^. According to the split time of the dominant strain and other strains in the phylogenetic trees (Data S1), we estimated the emergence times of the mutations. Specifically, the total incubation course was normalized to 1. If a dominant strain A (with mutations X1, X2 and X3) had outgroups B, C and D from near to far subsequently, X1 was not available in B, C and D; X2 was available in B but not available in C and D; and X3 was available in A, B and C but not available in D; then the estimated emergence time of X1, X2 and X3 are 1/4, 2/4, and 3/4, respectively. We wrote PERL scripts to evaluate the function of T492I-promoted mutations on the basis of published records and performed the statistical analyses by R.

### Viral subgenomic RNA assay and genomic RNA assay

Approximately 1×10^6^ cells were seeded into 6-well plates and cultured in 5% CO at 371°C for 12 h. The virus was serially diluted in DMEM containing 2% FBS, and 200 μL aliquots were added to the cells. After infection, total RNA from the infectious cell lysate was extracted via an RNeasy Mini Kit (QIAGEN, Hilden, Germany). RT-PCR was performed via an iTaq Universal SYBR Green One-Step Kit (Bio-Rad) and an ABI StepOnePlus PCR system (Thermo Fisher Scientific, CA, USA) according to the manufacturer’s instructions. The viral subgenomic RNA assay was performed with primers that target envelope protein (E) gene and ORF1ab sequences as previously described ^18^ (Data S1).

### RNA isolation and qRT-PCR

Total SARS-CoV-2 RNA was extracted by using the Analytik Jena Kit (QIAGEN, Hilden, Germany), followed by reverse transcription into cDNA with a high capacity cDNA reverse transcription kit (Thermo Fisher Scientific). The quantification of mRNA levels was conducted via an iTaq Universal SYBR Green One-Step Kit (Bio-Rad). The assay was performed on an ABI StepOnePlus PCR system (Thermo Fisher Scientific). The primers used were listed in Data S1.

### Validation of the infection performance of the evolved viruses in different species

The Calu-3 cell lines stably expressing ACE2 orthologs and SARS-CoV-2 variant-bearing LVpp were developed as previously described ^44^. The cDNA of ACE2 orthologs (human, monkey, hamster, rabbit, cat, pig, dog, horse, camel, ferret, bat and mouse) were synthesized, cloned and inserted into the pCDH-CMV-MCS-EF1-RFP-T2A-Puro vector. The lentiviruses carrying ACE2 orthologs were produced in HEK-293T cells and were harvested to infect the Calu-3 cell lines. The stably-transduced cells were enriched via Puromycin selection. The spike-expressing plasmid of evolved viruses, the packing plasmid, and the mNeonGreen reporter vector were cotransfected into HEK-293T cells to generate SARS-CoV-2 variant-bearing LVpp. The p24 concentrations of viral stocks were determined via a p24 Rapid Titer Kit (Takara). For the LVpp infection assay, the Calu-3 cell lines stably expressing ACE2 orthologs were seeded into each well of 96-well plates and cultured at 37 °C under 5% CO2. After 16 h of culture, the cells were incubated with a virus inoculum of 10 ng p24. After 2 days of infection, the number of mNeonGreen-activated cells in each well was determined and expressed as the number of green-fluorescent units per well (GFU/well).

### ELISA

The binding ability of the SARS-CoV-2 RBD to human ACE2 (hACE2) was detected by ELISA using the RayBio COVID-19 Spike-ACE2 Binding Assay Kit II. Different recombinant RBD proteins were added to hACE2-coated plates and incubated overnight at 4 °C with gentle shaking. The solution was subsequently discarded and 100 μL of 1×HRP-conjugated IgG antibody was added to the plates. The reaction mixture was allowed to react for 1 h at room temperature. Finally, 100 µL of TMB One-Step Substrate Reagent was added to the plates. The mixture was incubated at room temperature for an additional 30 min, and then 50 μL of Stop Solution was added. The absorbance was immediately read at 405 nm.

### Surface plasmon resonance and protein complex structure analysis

Recombinant hACE2 protein was immobilized on a CM5 Chip and different concentrations of SARS-CoV-2 RBD proteins were injected into the hACE2-immobilized flow cell. The K_D_ was calculated via the steady-state affinity obtained for each concentration. The flow rate was 20 mL/min for 200 s and dissociation for 400 s. The structures of the hACE2 protein and SARS-CoV-2 RBD proteins were predicted by AlphaFold2 ^82^, and the inactive sites were removed. ZDOCK was subsequently used to achieve protein-protein rigid docking, and RosettaDock ^96^ was used to achieve protein-protein flexible docking. The preliminary conformation of the global docking was optimized with Rosetta ^97^ in two rounds to obtain the final model of the hACE2/RBD complex. The binding interface of the complex was comprehensively characterized and systematically analyzed via the interaction analysis platforms PDBsum ^83^ and PLIP ^84^.

For the phylogenetic-based clustering analyses, on the basis of the OD and KD values from the ELLSA and SPR experiments, we used the dynamic clustering k-means clustering method. Specifically, we used the “elbow method” as the standard. We used the function “fviz_clust”, the contour coefficients and the WSS (sum of squared errors within clusters) to determine the optimal number of clusters. We used the function “kmeans” with the parameters “centers=4, start=50” to calculate the distance between each object and the cluster center, assign each object to the nearest cluster center, cluster the surrounding points, and then calculate the average value of each class. The calculated results were used as the classification points, and the above process was repeated continuously until the classification results converged. The clustering results were visualized via the R package Factoextra.

### Analyses of the expression of APOBEC and ADAR enzymes

Calu-3 and Vero-E6 cells were seeded into each well of 6-well plates and cultured at 37 °C under 5% CO2. SARS-CoV-2 variants (WT, T492I, Delta and I492T) were inoculated into a culture at an MOI of 5. After 12 h of infection, the inoculum was removed, and the culture was washed three times with PBS. Infectious cell lysates were harvested, and total RNA was subsequently extracted via a PureLink RNA Mini Kit (Thermo Fisher Scientific Inc., CA, USA). RT-PCR was performed via a SYBR PrimeScript RT-PCR Kit (Takara, Otsu, Shiga, Japan) and qRT-PCR was performed via a TB Green Premix ExTaq II Kit (Takara) on a Bio-Rad CFX-96 system (Bio-Rad, Hercules, CA, USA). Thermal cycling was performed at 95 °C for 30 s, followed by 39 cycles of 95° C for 5 s, and 60°C for 30 s. The sequences of primers used were listed in Data S1.

### Quantitative and statistical analysis

Chi-square tests, Wilcoxon tests, correlation tests and binomial tests were performed in R. We wrote Perl scripts to classify the strains into lineages and quantified the IFs of these lineages. The source data are from GISAID ^85^ and Nextstrain ^72^. Heatmaps, box-plots, scatter-plots and raincloud plots were generated via the R libraries “gdata”, “ggplot2”, “RColorBrewer”, “scales”, “ggsci”, “RColorBrewer”, “ggridges”, “pheatmap”, “cowplot”, “dplyr”, “readr” and “Factoextra”.

## SUPPLEMENTARY INFORMATION

### Supplementary Figures S1-S5

Figure S1. Additional evidence suggesting Omicron-biased evolution induced by T492I, related to Figure 1.

Figure S2. Results of evolutionary analysis and function prediction, related to Figure 2.

Figure S3. Results of the surface plasmon resonance (SPR) experiments for measuring the hACE2 affinities of the estimated dominant strains after experimental evolution in the 90-day runs, related to Figure 4.

Figure S4. Molecular docking results of the hACE2/evolved RBD complex for the estimated dominant strains of the 90-day 492I populations and the control strains (A-J), and the frequencies of nucleotide types near the 3’ end of G > A mutations for the 90-day and 45-day runs (K and L).

Figure S5. Distribution of the sequencing depths throughout the genome.

### Supplemental Tables S1-S6

Table S1. Table S1. Information on high-frequency nonsynonymous mutations, related to Figure 1.

Table S2. Table S2. Information on high-frequency synonymous mutations, related to Figure 1.

Table S3. Table S3. Information on high-frequency noncoding region mutations, related to Figure 1.

Table S4. Table S4. Mutations in the spike RBD region of the predicted predominant strains of the 90-day runs, related to Figure 4.

Table S5. Table S5. Protein-protein interaction types in different groups, related to Figure 4.

Table S6. Table S6. The GISAID IDs of the samples selected to identify the mutations of VOCs, related to Figure 1.

### Data S1

The Supplementary ZIP File () includes the sliding window views showing the genetic differentiation, nucleotide diversity and Tajima’s D of the 90-day and 45-day runs; phylogenetic trees of the constructed virus strains in the replicates of the 90-day and 45-day runs; pSARS-CoV-2 Delta (BAC clone); the primers used in the study; and the quality control reports of the source data. The contents are related to Figures 1-6.

