## Supplementary Figure for "NSP4 mutation T492I drives rapid evolution of SARS-CoV-2 toward Omicron"

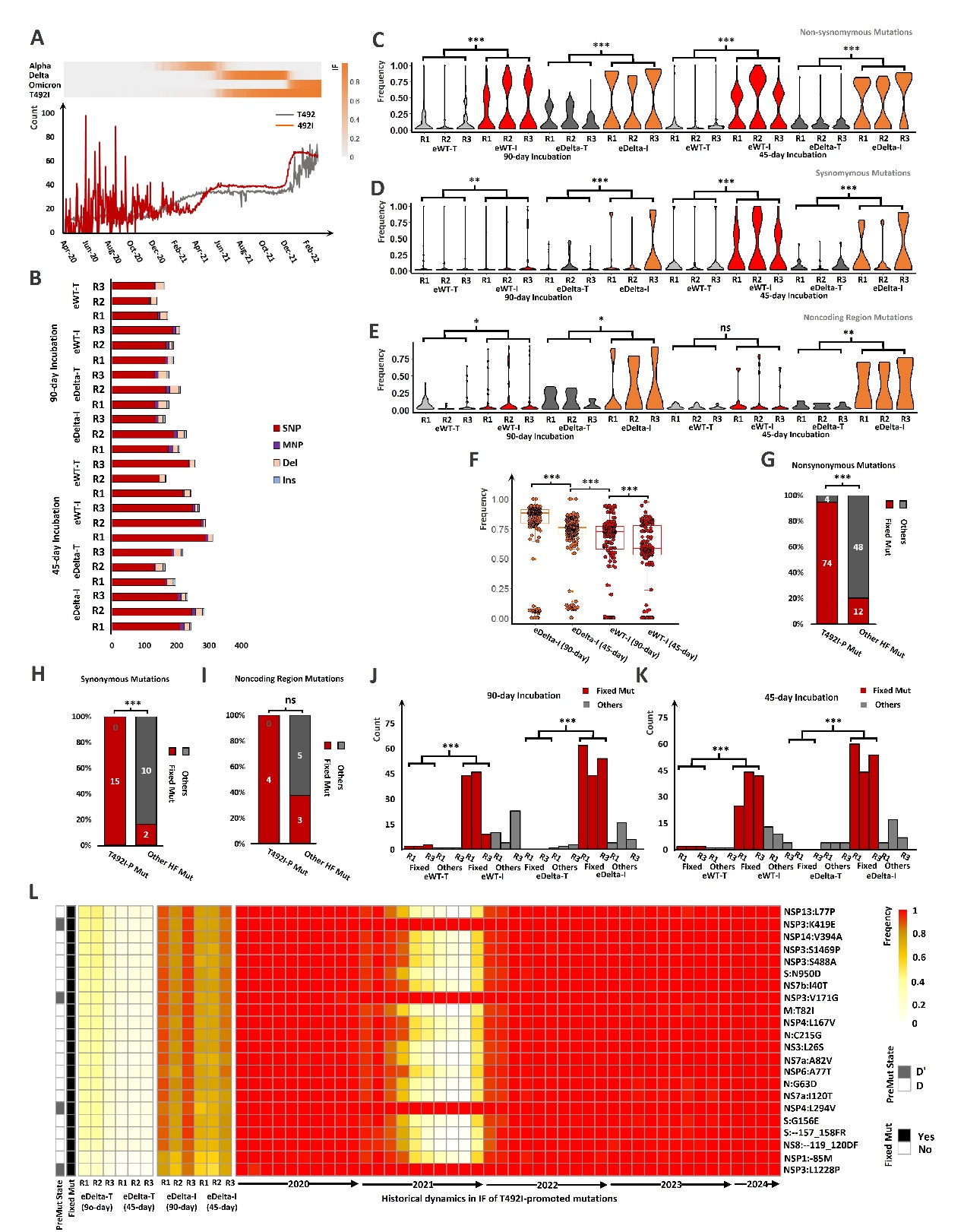

Figure S1. Additional evidence suggesting Omicron-biased evolution induced by T492I, related to Figure 1.

(A) Global dynamics of the median mutation counts in the strains bearing T492 and those bearing 492I. The changes in IF of historical dominant VOCs and T492I were shown at the top. (B) The counts of single nucleotide polymorphism (SNP), multiple nucleotide polymorphism (MNP), deletion (Del) and insertion (Ins) in different evolved populations. (C-E) Distributions of the mutation frequencies in the replicates of different runs, for the nonsynonymous (C), synonymous (D) and noncoding region mutations (E) respectively. Statistics were performed via Kolmogorov-Smirnov tests. (F) Distributions of the mutations in the evolved strains of the 492I runs. (G-I) Comparisons of the fractions of fixed Omicron mutations (Fixed Mut) in T492I-promoted mutations (T492I-P Mut) and other high-frequency mutations (Other HF Mut) for nonsynonymous (G), synonymous (H) and noncoding region (I) mutations. (J, K) Comparisons of the counts of fixed Omicron mutations (Fixed Mut) and other mutations (Others) in the reconstructed dominant strain of the 90-day runs (J) and the 45-day runs (K). (L) Heatmap displays the frequencies of the Delta-specific reverse mutations in the T492I-promoted mutations. The column ‘PreMut State’ shows the amino acid types of premutation states. D denotes the premutation state is shared by most Delta strains, and D’ denotes the premutation state is not sheared by most Delta strains, according to the statistics of the epidemiological data provided by Nextstrain. Other legends in this figure follow those in Figure 1.

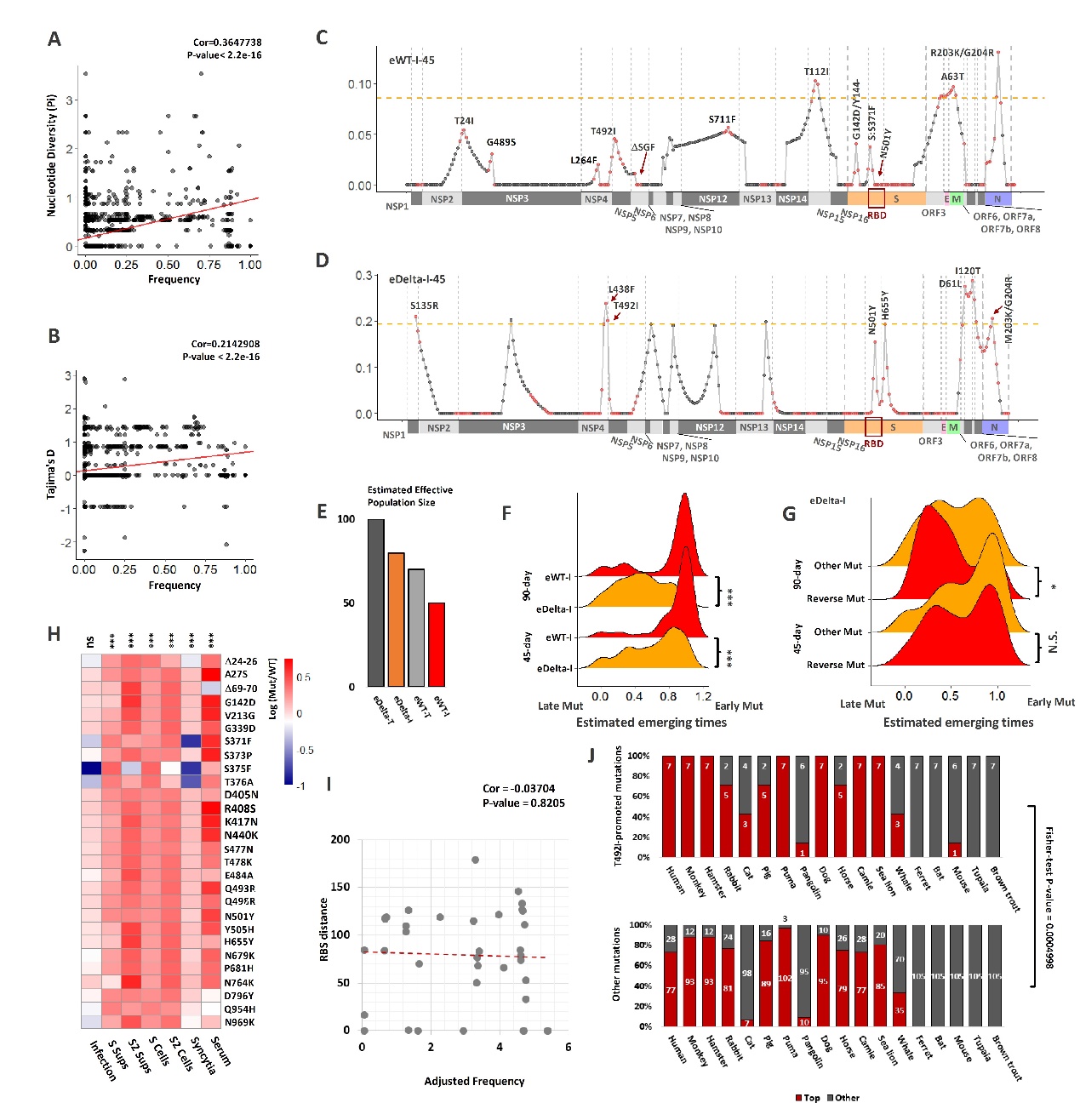

Figure S2. Results of evolutionary analysis and function prediction, related to Figure 2.

(A, B) Correlation analysis results between the frequencies of the mutations in all runs and the population genetics parameters, including the nucleotide diversity (Pi) and the values of Tajima’s D (B). (C, D) Sliding window views showing the CLR peaks and the thresholds (orange dotted line), for eWT-I (C) and eDelta-I of the 45-day runs (D). (E) The estimated effective population size across different populations. (F) Comparisons of the distributions of the estimated emergence times of mutations across different 492I populations. (G) Comparisons of the distributions of the estimated emergence times between the Delta-specific reverse mutations and other mutations in eDelta-I for both the 90-day and 45-day populations. In (F) and (G), the statistics here were performed via Kolmogorov-Smirnov tests. (H) Impacts of T492I-promoted mutations on normalized pseudo particle infection of CaCo-2 cells (Infection), levels of full-length spike in supernatants (S Sups) and cells (S Cells), levels of S2 Spike subunit in the supernatants (S2 Sups) and cells (S2 Cells), binding of spike to ACE2 (ACE2 interaction), automated quantification of syncytia formation in HEK293T cells expressing the indicated mutant S proteins and human ACE2 (Syncytia), and average TCID50 values obtained for neutralization of the indicated mutant S proteins by sera from five vaccinated individuals relative to those obtained for Hu-1 S (Serum) on the basis of published data ^1^. Log (Mut/WT) refers to a log transformation of the ratio of the corresponding values of the premutation state and the postmutation state. Binomial tests were performed to evaluate the ratio of the mutations with a Log > 0 to those with a Log < 0. (I) Correlation analysis between RBD distances (the distances of mutations to the receptor binding domain) and the adjusted frequencies (the sum of the frequencies of mutations in the runs of eDelta-I-90 and eWT-I-90) for identified mutations. The RBD distance information is based on published investigation results ^2^. (J) Comparison of the infection performance of T492I-promoted mutations (upper panel) and other mutations (lower pannel) in 112 lentiviral-pseudotyped particles bearing the single-site RBD-mutated spike in H1299-expressing ACE2 orthologs, on the basis of published data ^3^. ‘Top’ and ‘Other’ denote infection performances of ≥ 7500 GFU/well and < 7500 GFU/well, respectively. ’*’ denotes P-value < 0.1, ‘**’ denotes P-value <0.05, and ‘***’ denotes P-value <0.01. ‘ns’ denotes no significance.

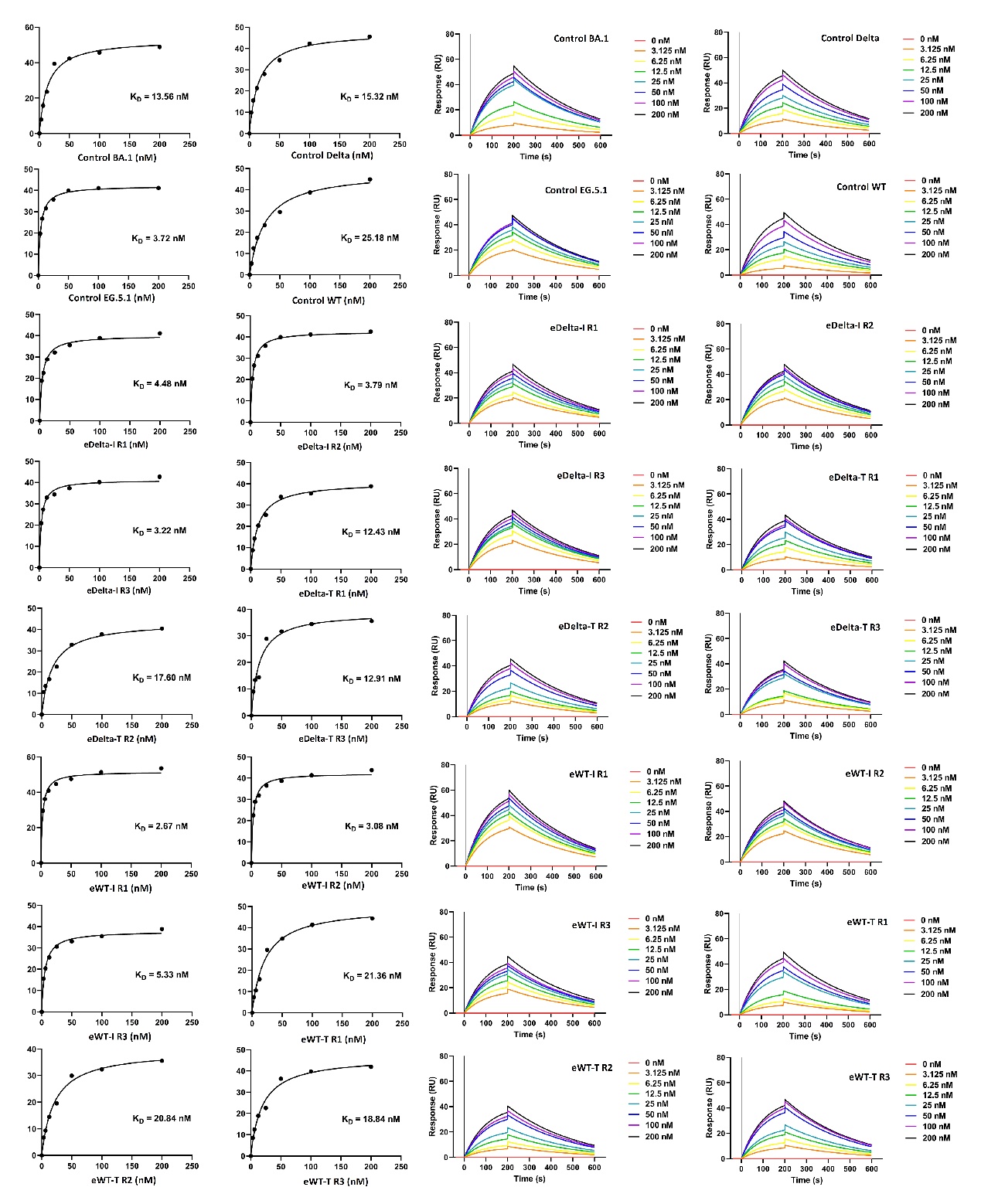

Figure S3. Results of the surface plasmon resonance (SPR) experiments for measuring the hACE2 affinities of the estimated dominant strains after experimental evolution in the 90-day runs, related to Figure 4.

The plots on the left are the results of measuring the affinity of the ligand for the receptor (K_D_), and the plots on the right are the results of measuring the response in resonance units (RUs).

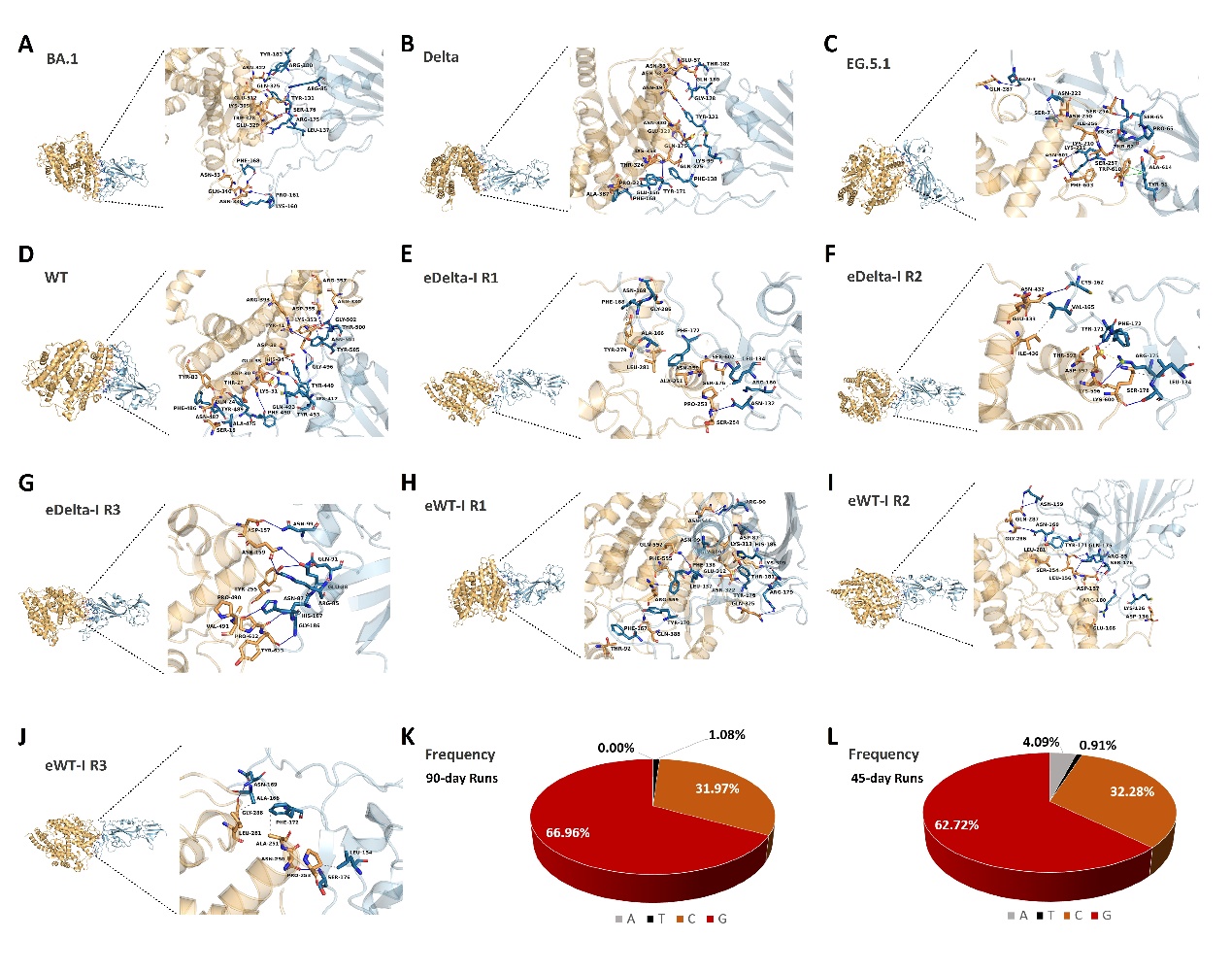

Figure S4. Molecular docking results of the hACE2/evolved RBD complex for the estimated dominant strains of the 90-day 492I populations and the control strains (A-J), and the frequencies of nucleotide types near the 3’ end of G > A mutations for the 90-day and 45-day runs (K and L). These figures are related to Figures 4 and 5. The legends follow those in Figure 4.

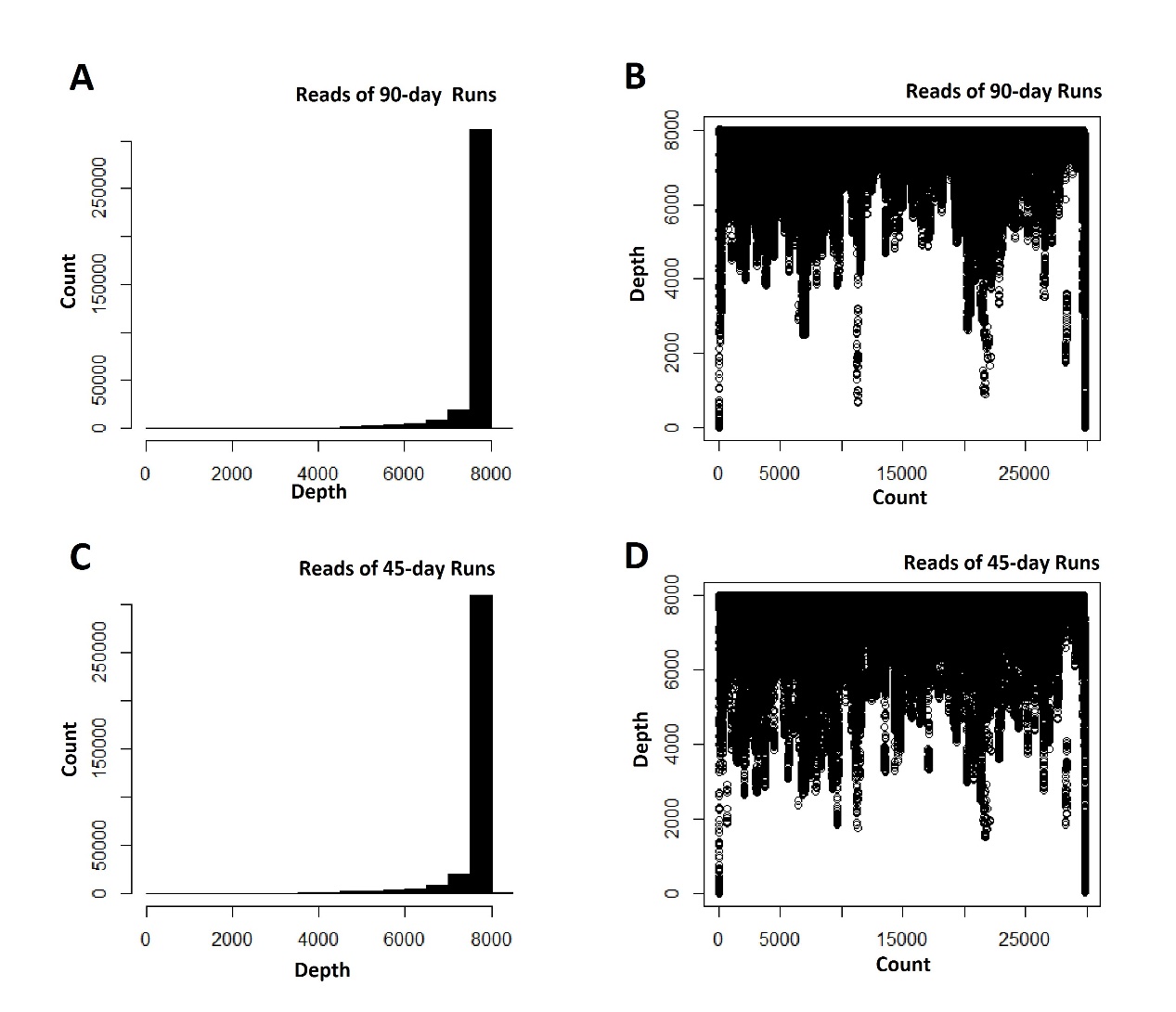

Figure S5. Distribution of the sequencing depths throughout the genome.

(A) Distribution of the sequencing depth for the reads of all 90-day runs and (B) Sequencing depths along the genome for the 90-day runs. (C) and (D) are those for the 45-day runs. This figure is related to Figures 1 and 2.
