## Supplementary material for "NSP4 mutation T492I drives rapid evolution of SARS-CoV-2 toward Omicron": Data S1: Description.pdf

**File Description**

Figures (Fig*.jpg or Fig*.pdf files)

Fig 1. Sliding window views display the genetic differentiation (Fst, A), nucleotide diversity (Pi, B) and Tajima's D (C) of the evolved populations in the 90-day runs. The points colored in red denote the regions with high-frequency mutations.

Fig 2. Sliding window views display the genetic differentiation (Fst, A), nucleotide diversity (Pi, B) and Tajima's D (C) of the 45-day runs. The legends follow those in Fig 1.

Fig 3. Phylogenetic trees of the reconstructed virus strains in the populations of the 90-day runs. eBayes denotes the likelihoods calculated via approximate Bayes test. The frequency of the dominant strain in different populations are shown on the right and colored in red.

Fig 4. Phylogenetic trees of the reconstructed virus strains in the populations of the 90-day runs. Legends follow those in Fig 3.

Fig 5. pSARS-CoV-2 Delta (BAC clone).

Tables (Tables.pdf)

Table 1. Mutagenesis primers used in the generation of SARS-CoV-2 mutants.

Table 2. Primers used for the amplification of the overlapping fragments of SARS-CoV-2 Delta.

Table 3. Primers that target envelope protein (E) gene and Orf1ab sequences.

Table 4. Primers used in qRT-PCR.

Table 5. Primers used for the analyses of the expression of APOBEC and ADAR enzymes.

The file "FASTQC.zip" contains the quality control reports of all runs.
