## Supplementary material for "NSP4 mutation T492I drives rapid evolution of SARS-CoV-2 toward Omicron": Data S1: Fig 5.pdf

>pSARS-CoV-2 Delta
TCGAGTGAGCGAGGAAGCACCAGGGAACAGCACTTATATATTCTGCTTACACACGATGCCTGAAAAAACTTCCCTTGGGG
TTATCCACTTATCCACGGGGATATTTTTATAATTATTTTTTTTATAGTTTTTAGATCTTCTTTTTTAGAGCGCCTTGTAG
GCCTTTATCCATGCTGGTTCTAGAGAAGGTGTTGTGACAAATTGCCCTTTCAGTGTGACAAATCACCCTCAAATGACAGT
CCTGTCTGTGACAAATTGCCCTTAACCCTGTGACAAATTGCCCTCAGAAGAAGCTGTTTTTTCACAAAGTTATCCCTGCT
TATTGACTCTTTTTTATTTAGTGTGACAATCTAAAAACTTGTCACACTTCACATGGATCTGTCATGGCGGAAACAGCGGT
TATCAATCACAAGAAACGTAAAAATAGCCCGCGAATCGTCCAGTCAAACGACCTCACTGAGGCGGCATATAGTCTCTCCC
GGGATCAAAAACGTATGCTGTATCTGTTCGTTGACCAGATCAGAAAATCTGATGGCACCCTACAGGAACATGACGGTATC
TGCGAGATCCATGTTGCTAAATATGCTGAAATATTCGGATTGACCTCTGCGGAAGCCAGTAAGGATATACGGCAGGCATT
GAAGAGTTTCGCGGGGAAGGAAGTGGTTTTTTATCGCCCTGAAGAGGATGCCGGCGATGAAAAAGGCTATGAATCTTTTC
CTTGGTTTATCAAACGTGCGCACAGTCCATCCAGAGGGCTTTACAGTGTACATATCAACCCATATCTCATTCCCTTCTTT
ATCGGGTTACAGAACCGGTTTACGCAGTTTCGGCTTAGTGAAACAAAAGAAATCACCAATCCGTATGCCATGCGTTTATA
CGAATCCCTGTGTCAGTATCGTAAGCCGGATGGCTCAGGCATCGTCTCTCTGAAAATCGACTGGATCATAGAGCGTTACC
AGCTGCCTCAAAGTTACCAGCGTATGCCTGACTTCCGCCGCCGCTTCCTGCAGGTCTGTGTTAATGAGATCAACAGCAGA
ACTCCAATGCGCCTCTCATACATTGAGAAAAAGAAAGGCCGCCAGACGACTCATATCGTATTTTCCTTCCGCGATATCAC
TTCCATGACGACAGGATAGTCTGAGGGTTATCTGTCACAGATTTGAGGGTGGTTCGTCACATTTGTTCTGACCTACTGAG
GGTAATTTGTCACAGTTTTGCTGTTTCCTTCAGCCTGCATGGATTTTCTCATACTTTTTGAACTGTAATTTTTAAGGAAG
CCAAATTTGAGGGCAGTTTGTCACAGTTGATTTCCTTCTCTTTCCCTTCGTCATGTGACCTGATATCGGGGGTTAGTTCG
TCATCATTGATGAGGGTTGATTATCACAGTTTATTACTCTGAATTGGCTATCCGCGTGTGTACCTCTACCTGGAGTTTTT
CCCACGGTGGATATTTCTTCTTGCGCTGAGCGTAAGAGCTATCTGACAGAACAGTTCTTCTTTGCTTCCTCGCCAGTTCG
CTCGCTATGCTCGGTTACACGGCTGCGGCGAGCATCACGTGCTATAAAAATAATTATAATTTAAATTTTTTAATATAAAT
ATATAAATTAAAAATAGAAAGTAAAAAAAGAAATTAAAGAAAAAATAGTTTTTGTTTTCCGAAGATGTAAAAGACTCTAG
GGGGATCGCCAACAAATACTACCTTTTACCTTGCTCTTCCTGCTCTCAGGTATTAATGCCGAATTGTTTCATCTTGTCTG
TGTAGAAGACCACACACGAAAATCCTGTGATTTTACATTTTACTTATCGTTAATCGAATGTATATCTATTTAATCTGCTT
TTCTTGTCTAATAAATATATATGTAAAGTACGCTTTTTGTTGAAATTTTTTAAACCTTTGTTTATTTTTTTTTCTTCATT
CCGTAACTCTTCTACCTTCTTTATTTACTTTCTAAAATCCAAATACAAAACATAAAAATAAATAAACACAGAGTAAATTC
CCAAATTATTCCATCATTAAAAGATACGAGGCGCGTGTAAGTTACAGGCAAGCGATCCTAGTACACTCTATATTTTTTTA
TGCCTCGGTAATGATTTTCATTTTTTTTTTTCCACCTAGCGGATGACTCTTTTTTTTTCTTAGCGATTGGCATTATCACAT
AATGAATTATACATTATATAAAGTAATGTGATTTCTTCGAAGAATATACTAAAAAATGAGCAGGCAAGATAAACGAAGGC
AAAGATGACAGAGCAGAAAGCCCTAGTAAAGCGTATTACAAATGAAACCAAGATTCAGATTGCGATCTCTTTAAAGGGTG
GTCCCCTAGCGATAGAGCACTCGATCTTCCCAGAAAAAGAGGCAGAAGCAGTAGCAGAACAGGCCACACAATCGCAAGTG
ATTAACGTCCACACAGGTATAGGGTTTCTGGACCATATGATACATGCTCTGGCCAAGCATTCCGGCTGGTCGCTAATCGT
TGAGTGCATTGGTGACTTACACATAGACGACCATCACACCACTGAAGACTGCGGGATTGCTCTCGGTCAAGCTTTTAAAG
AGGCCCTAGGGGCCGTGCGTGGAGTAAAAAGGTTTGGATCAGGATTTGCGCCTTTGGATGAGGCACTTTCCAGAGCGGTG
GTAGATCTTTCGAACAGGCCGTACGCAGTTGTCGAACTTGGTTTGCAAAGGGAGAAAGTAGGAGATCTCTCTTGCGAGAT
GATCCCGCATTTTCTTGAAAGCTTTGCAGAGGCTAGCAGAATTACCCTCCACGTTGATTGTCTGCGAGGCAAGAATGATC
ATCACCGTAGTGAGAGTGCGTTCAAGGCTCTTGCGGTTGCCATAAGAGAAGCCACCTCGCCCAATGGTACCAACGATGTT
CCCTCCACCAAAGGTGTTCTTATGTAGTTTTACACAGGAGTCTGGACTTGACGCTAGTGATAATAAGTGACTGAGGTATG
TGCTCTTCTTATCTCCTTTTGTAGTGTTGCTCTTATTTTAAACAACTTTGCGGTTTTTTGATGACTTTGCGATTTTGTTG
TTGCTTTGCAGTAAATTGCAAGATTTAATAAAAAAACGCAAAGCAATGATTAAAGGATGTTCAGAATGAAACTCATGGAA
ACACTTAACCAGTGCATAAACGCTGGTCATGAAATGACGAAGGCTATCGCCATTGCACAGTTTAATGATGACAGCCCGGA
AGCGAGGAAAATAACCCGGCGCTGGAGAATAGGTGAAGCAGCGGATTTAGTTGGGGTTTCTTCTCAGGCTATCAGAGATG
CCGAGAAAGCAGGGCGACTACCGCACCCGGATATGGAAATTCGAGGACGGGTTGAGCAACGTGTTGGTTATACAATTGAA
CAAATTAATCATATGCGTGATGTGTTTGGTACGCGATTGCGACGTGCTGAAGACGTATTTCCACCGGTGATCGGGGTTGC
TGCCCATAAAGGTGGCGTTTACAAAACCTCAGTTTCTGTTCATCTTGCTCAGGATCTGGCTCTGAAGGGGCTACGTGTTT
TGCTCGTGGAAGGTAACGACCCCCAGGGAACAGCCTCAATGTATCACGGATGGGTACCAGATCTTCATATTCATGCAGAA
GACACTCTCCTGCCTTTCTATCTTGGGGAAAAGGACGATGTCACTTATGCAATAAAGCCCACTTGCTGGCCGGGGCTTGA
CATTATTCCTTCCTGTCTGGCTCTGCACCGTATTGAAACTGAGTTAATGGGCAAATTTGATGAAGGTAAACTGCCCACCG
ATCCACACCTGATGCTCCGACTGGCCATTGAAACTGTTGCTCATGACTATGATGTCATAGTTATTGACAGCGCGCCTAAC
CTGGGTATCGGCACGATTAATGTCGTATGTGCTGCTGATGTGCTGATTGTTCCCACGCCTGCTGAGTTGTTTGACTACAC
CTCCGCACTGCAGTTTTTCGATATGCTTCGTGATCTGCTCAAGAACGTTGATCTTAAAGGGTTCGAGCCTGATGTACGTA
TTTTGCTTACCAAATACAGCAATAGCAATGGCTCTCAGTCCCCGTGGATGGAGGAGCAAATTCGGGATGCCTGGGGAAGC
ATGGTTCTAAAAAATGTTGTACGTGAAACGGATGAAGTTGGTAAAGGTCAGATCCGGATGAGAACTGTTTTTGAACAGGC
CATTGATCAACGCTCTTCAACTGGTGCCTGGAGAAATGCTCTTTCTATTTGGGAACCTGTCTGCAATGAAATTTTCGATC
GTCTGATTAAACCACGCTGGGAGATTAGATAATGAAGCGTGCGCCTGTTATTCCAAAACATACGCTCAATACTCAACCGG
TTGAAGATACTTCGTTATCGACACCAGCTGCCCCGATGGTGGATTCGTTAATTGCGCGCGTAGGAGTAATGGCTCGCGGT
AATGCCATTACTTTGCCTGTATGTGGTCGGGATGTGAAGTTTACTCTTGAAGTGCTCCGGGGTGATAGTGTTGAGAAGAC
CTCTCGGGTATGGTCAGGTAATGAACGTGACCAGGAGCTGCTTACTGAGGACGCACTGGATGATCTCATCCCTTCTTTTC
TACTGACTGGTCAACAGACACCGGCGTTCGGTCGAAGAGTATCTGGTGTCATAGAAATTGCCGATGGGAGTCGCCGTCGT

AAAGCTGCTGCACTTACCGAAAGTGATTATCGTGTTCTGGTTGGCGAGCTGGATGATGAGCAGATGGCTGCATTATCCAG
ATTGGGTAACGATTATCGCCCAACAAGTGCTTATGAACGTGGTCAGCGTTATGCAAGCCGATTGCAGAATGAATTTGCTG
GAAATATTTCTGCGCTGGCTGATGCGGAAAATATTTCACGTAAGATTATTACCCGCTGTATCAACACCGCCAAATTGCCT
AAATCAGTTGTTGCTCTTTTTCTCACCCCGGTGAACTATCTGCCCGGTCAGGTGATGCACTTCAAAAAGCCTTTACAGA
TAAAGAGGAATTACTTAAGCAGCAGGCATCTAACCTTCATGAGCAGAAAAAAGCTGGGGTGATATTTGAAGCTGAAGAAG
TTATCACTCTTTTAACTTCTGTGCTTAAAACGTCATCTGCATCAAGAACTAGTTTAAGCTCACGACATCAGTTTGCTCCT
GGAGCGACAGTATTGTATAAGGGCGATAAAATGGTGCTTAACCTGGACAGGTCTCGTGTTCCAACTGAGTGTATAGAGAA
AATTGAGGCCATTCTTAAGGAACTTGAAAAGCCAGCACCCTGATGCGACCTCGTTTTAGTCTACGTTTATCTGTCTTTAC
TTAATGTCCTTTGTTACAGGCCAGAAAGCATAACTGGCCTGAATATTCTCTCTGGGCCCACTGTTCCACTTGTATCGTCG
GTCTGATAATCAGACTGGGACCACGGTCCCACTCGTATCGTCGGTCTGATTATTAGTCTGGGACCACGGTCCCACTCGTA
TCGTCGGTCTGATTATTAGTCTGGGACCACGGTCCCACTCGTATCGTCGGTCTGATAATCAGACTGGGACCACGGTCCCA
CTCGTATCGTCGGTCTGATTATTAGTCTGGGACCATGGTCCCACTCGTATCGTCGGTCTGATTATTAGTCTGGGACCACG
GTCCCACTCGTATCGTCGGTCTGATTATTAGTCTGGAACCACGGTCCCACTCGTATCGTCGGTCTGATTATTAGTCTGGG
ACCACGGTCCCACTCGTATCGTCGGTCTGATTATTAGTCTGGGACCACGATCCCACTCGTGTTGTCGGTCTGATTATCGG
TCTGGGACCACGGTCCCACTTGTATTGTCGATCAGACTATCAGCGTGAGACTACGATTCCATCAATGCCTGTCAAGGGCA
AGTATTGACATGTCGTCGTAACCTGTAGAACGGAGTAACCTCGGTGTGCGGTTGTATGCCTGCTGTGGATTGCTGCTGTG
TCCTGCTTATCCACAACATTTTGCGCACGGTTATGTGGACAAAATACCTGGTTACCCAGGCCGTGCCGGCACGTTAACCG
GGCTGCATCCGATGCAAGTGTGTCGCTGTCGACGAGCTCGCGAGCTCGCTCAATATTGGCCATTAGCCATATTATTCATT
GGTTATATAGCATAAATCAATATTGGCTATTGGCCATTGCATACGTTGTATCTATATCATAATATGTACATTTATATTGG
CTCATGTCCAATATGACCGCCATGTTGGCATTGATTATTGACTAGTTATTAATAGTAATCAATTACGGGGTCATTAGTTC
ATAGCCCATATATGGAGTTCCGCGTTACATAACTTACGGTAAATGGCCCGCCTGGCTGACCGCCCAACGACCCCCGCCCA
TTGACGTCAATAATGACGTATGTTCCCATAGTAACGCCAATAGGGACTTTCCATTGACGTCAATGGGTGGAGTATTTACG
GTAAACTGCCCACTTGGCAGTACATCAAGTGTATCATATGCCAAGTACGCCCCCTATTGACGTCAATGACGGTAAATGGC
CCGCCTGGCATTATGCCCAGTACATGACCTTATGGGACTTTCCTACTTGGCAGTACATCTACGTATTAGTCATCGCTATT
ACCATGGTGATGCGGTTTTGGCAGTACATCAATGGGCGTGGATAGCGGTTTGACTCACGGGGATTTCCAAGTCTCCACCC
CATTGACGTCAATGGGAGTTTGTTTTGGCACCAAAATCAACGGGACTTTCCAAAATGTCGTAACAACTCCGCCCCATTGA
CGCAAATGGGCGGTAGGCGTGTACGGTGGGAGGTCTATATAAGCAGAGCTCGTTTAGTGAACCGTATATTAGGTTTATAC
CTTCCCAGGTAACAAACCAACCAACTTTCGATCTCTTGTAGATCTGTTCTCTAAACGAACTTTAAAATCTGTGTGGCTGT
CACTCGGCTGCATGCTTAGTGCACTCACGCAGTATAATTAATAACTAATTACTGTCGTTGACAGGACACGAGTAACTCGT
CTATCTTCTGCAGGCTGCTTACGGTTTCGTCCGTGTTGCAGCCGATCATCAGCACATCTAGGTTTCGTCCGGGTGTGACC
GAAAGGTAAGATGGAGAGCCTTGTCCCTGGTTTCAACGAGAAAACACACGTCCAACTCAGTTTGCCTGTTTTACAGGTTC
GCGACGTGCTCGTACGTGGCTTTGGAGACTCCGTGGAGGAGGTCTTATCAGAGGCACGTCAACATCTTAAAGATGGCACT
TGTGGCTTAGTAGAAGTTGAAAAAGGCGTTTTGCCTCAACTTGAACAGCCCTATGTGTTCATCAAACGTTCGGATGCTCG
AACTGCACCTCATGGTCATGTGGTTGAGCTGGTAGCAGAACTCGAAGGCATTCAGTACGGTCGTAGTGGTGAGACACTTG
GTGTCCTTGTCCCTCATGTGGGCGAAATACCAGTGGCTTACCGCAAGGTTCTTCTTCGTAAGAACGGTAATAAAGGAGCT
GGTGGCCATAGTTACGGCGCCGATCTAAAGTCATTTGACTTAGGCGACGAGCTTGGCACTGATCCTTATGAAGATTTTCA
AGAAAACTGGAACACTAAACATAGCAGTGGTGTTACCCGTGAACTCATGCGTGAGCTTAACGGAGGGGCATACACTCGCT
ATGTCGATAACAACTTCTGTGGCCCTGATGGCTACCCTCTTGAGTGCATTAAAGACCTTCTAGCACGTGCTGGTAAAGCT
TCATGCACTTTGTCCGAACAACTGGACTTTATTGACACTAAGAGGGGTGTATACTGCTGCCGTGAACATGAGCATGAAAT
TGCTTGGTACACGGAACGTTCTGAAAAGAGCTATGAATTGCAGACACCTTTTGAAATTAAATTGGCAAAGAAATTTGACA
CCTTCAATGGGGAATGTCCAAATTTTGTATTTCCCTTAAATTCCATAATCAAGACTATTCAACCAAGGGTTGAAAAGAAA
AAGCTTGATGGCTTTATGGGTAGAATTCGATCTGTCTATCCAGTTGCGTCACCAAATGAATGCAACCAAATGTGCCTTTC
AACTCTCATGAAGTGTGATCATTGTGGTGAAACTTCATGGCAGACGGGCGATTTTGTTAAAGCCACTTGCGAATTTTGTG
GCACTGAGAATTTGACTAAAGAAGGTGCCACTACTTGTGGTTACTTACCCCAAAATGCTGTTGTTAAAATTTATTGTCCA
GCATGTCACAATTCAGAAGTAGGACCTGAGCATAGTCTTGCCGAATACCATAATGAATCTGGCTTGAAAACCATTCTTCG
TAAGGGTGGTCGCACTATTGCCTTTGGAGGCTGTGTGTTCTCTTATGTTGGTTGCCATAACAAGTGTGCCTATTGGGTTC
CACGTGCTAGCGCTAACATAGGTTGTAACCATACAGGTGTTGTTGGAGAAGGTTCCGAAGGTCTTAATGACAACCTTCTT
GAAATACTCCAAAAAGAGAAAGTCAACATCAATATTGTTGGTGACTTTAAACTTAATGAAGAGATCGCCATTATTTTGGC
ATCTTTTTCTGCTTCCACAAGTGCTTTTGTGGAAACTGTGAAAGGTTTGGATTATAAAGCATTCAAACAAATTGTTGAAT
CCTGTGGTAATTTTAAAGTTACAAAAGGAAAAGCTAAAAAAGGTGCCTGGAATATTGGTGAACAGAAATCAATACTGAGT
CCTCTTTATGCATTTGCATCAGAGGCTGCTCGTGTTGTACGATCAATTTTCTCCCGCACTCTTGAAACTGCTCAAAATTC
TGTGCGTGTTTTACAGAAGGCCGCTATAACAATACTAGATGGAATTTCACAGTATTCACTGAGACTCATTGATGCTATGA
TGTTCACATCTGATTTGGCTACTAACAATCTAGTTGTAATGGCCTACATTACAGGTGGTGTTGTTCAGTTGACTTCGCAG
TGGCTAACTAACATCTTTGGCACTGTTTATGAAAAACTCAAACCCGTCCTTGATTGGCTTGAAGAGAAGTTTAAGGAAGG
TGTAGAGTTTCTTAGAGACGGTTGGGAAATTGTTAAATTTATCTCAACCTGTGCTTGTGAAATTGTCGGTGGACAAATTG
TCACCTGTGCAAAGGAAATTAAGGAGAGTGTTCAGACATTCTTTAAGCTTGTAAATAAATTTTTGGCTTTGTGTGCTGAC
TCTATCATTATTGGTGGAGCTAAACTTAAAGCCTTGAATTTAGGTGAAACATTTGTCACGCACTCAAAGGGATTGTACAG
AAAGTGTGTTAAATCCAGAGAAGAAACTGGCCTACTCATGCCTCTAAAAGCCCCAAAAGAAATTATCTTCTTAGAGGGAG
AAACACTTCCCACAGAAGTGTTAACAGAGGAAGTTGTCTTGAAAACTGGTGATTTACAACCATTAGAACAACCTACTAGT

GAAGCTGTTGAAGCTCCATTGGTTGGTACACCAGTTTGTATTAACGGGCTTATGTTGCTCGAAATCAAAGACACAGAAAA
GTACTGTGCCCTTGCACCTAATATGATGGTAACAAACAATACCTTCACACTCAAAGGCGGTGCACCAACAAAGGTTACTT
TTGGTGATGACACTGTGATAGAAGTGCAAGGTTACAAGAGTGTGAATATCACTTTTGAACTTGATGAAAGGATTGATAAA
GTACTTAATGAGAAGTGCTCTGCCTATACAGTTGAACTCGGTACAGAAGTAAATGAGTTCGCCTGTGTTGTGGCAGATGC
TGTCATAAAAACTTTGCAACCAGTATCTGAATTACTTACACCACTGGGCATTGATTTAGATGAGTGGAGTATGGCTACAT
ACTACTTATTTGATGAGTCTGGTGAGTTTAAATTGGCTTCACATATGTATTGTTCTTTTTACCCTCCAGATGAGGATGAA
GAAGAAGGTGATTGTGAAGAAGAAGAGTTTGAGCCATCAACTCAATATGAGTATGGTACTGAAGATGATTACCAAGGTAA
ACCTTTGGAATTTGGTGCCACTTCTGCTGCTCTTCAACCTGAAGAAGAGCAAGAAGAAGATTGGTTAGATGATGATAGTC
AACAAACTGTTGTTCAACAAGACGGCAGTGAGGACAATCAGACAACTACTATTCAAACAATTGTTGAGGTTCAACCTCAA
TTAGAGATGGAACTTACACCAGTTGTTCAGACTATTGAAGTGAATAGTTTTAGTGGTTATTTAAAACTTACTGACAATGT
ATACATTAAAAATGCAGACATTGTGGAAGAAGCTAAAAAGGTAAAACCAACAGTGGTTGTTAATGCAGCCAATGTTTACC
TTAAACATGGAGGAGGTGTTGCAGGAGCCTTAAATAAGGCTACTAACAATGCCATGCAAGTTGAATCTGATGATTACATA
GCTACTAATGGACCACTTAAAGTGGGTGGTAGTTGTGTTTTAAGCGGACACAATCTTGCTAAACACTGTCTTCATGTTGT
CGGCCCAAATGTTAACAAAGGTGAAGACATTCAACTTCTTAAGAGTGCTTATGAAAATTTTAATCAGCACGAAGTTCTAC
TTGCACCATTATTATCAGCTGGTATTTTTGGTGCTGACCCTATACATTCTTTAAGAGTTTGTGTAGATACTGTTCGCACA
AATGTCTACTTAGCTGTCTTTGATAAAAATCTCTATGACAAACTTGTTTCAAGCTTTTTGGAAATGAAGAGTGAAAAGCA
AGTTGAACAAAAGATCGCTGAGATTCCTAAAGAGGAAGTTAAGCCATTTATAACTGAAAGTAAACCTTCAGTTGAACAGA
GAAAACAAGATGATAAGAAAATCAAAGCTTGTGTTAAAGAAGTTACAACAACTCTGGAAGAAACTAAGTTCCTCACAGAA
AACTTGTTACTTTATATTGACATTAATGGCAATCTTCATCCAGATTCTGCCACTCTTGTTAGTGACATTGACATCACTTT
CTTAAAGAAAGATGCTCCATATATAGTGGGTGATGTTGTTCAAGAGGGTGTTTTAACTGCTGTGGTTATACCTACTAAAA
AGTCTGGTGGCACTACTGAAATGCTAGCGAAAGCTTTGAGAAAAGTGCCAACAGACAATTATATAACCACTTACCCGGGT
CAGGGTTTAAATGGTTACACTGTAGAGGAGGCAAAGACAGTGCTTAAAAAGTGTAAAAGTGCCTTTTACATTCTACCATC
TATTATCTCTAATGAGAAGCAAGAAATTCTTGGAACTGTTTCTTGGAATTTGCGAGAAATGCTTGCACATGCAGAAGAAA
CACGCAAATTAATGCCTGTCTGTGTGGAAACTAAAGCCATAGTTTCAACTATACAGCGTAAATATAAGGGTATTAAAATA
CAAGAGGGTGTGGTTGATTATGGTGCTAGATTTTACTTTTACACCAGTAAAACAACTGTAGCGTCACTTATCAACACACT
TAACGATCTAAATGAAACTCTTGTTACAATGCCACTTGGCTATGTAACACATGGCTTAAATTTGGAAGAAGCTGCTCGGT
ATATGAGATCTCTCAAAGTGCCAGCTACAGTTTCTGTTTCTTCACCTGATGCTGTTACAGCGTATAATGGTTATCTTACT
TCTTCTTCTAAAACACCTGAAGAACATTTTATTGAAACCATCTCACTTGCTGGTTCCTATAAAGATTGGTCCTATTCTGG
ACAATCTACACAACTAGGTATAGAATTTCTTAAGAGAGGTGATAAAAGTGTATATTACACTAGTAATCCTACCACATTCC
ACCTAGATGGTGAAGTTATCACCTTTGACAATCTTAAGACACTTCTTTCTTTGAGAGAAGTGAGGACTATTAAGGTGTTT
ACAACAGTAGACAACATTAACCTCCACACGCAAGTTGTGGACATGTCAATGACATATGGACAACAGTTTGGTCCAACTTA
TTTGGATGGAGCTGATGTTACTAAAATAAAACCTCATAATTCACATGAAGGTAAAACATTTTATGTTTTACCTAATGATG
ACACTCTACGTGTTGAGGCTTTTGAGTACTACCACACAACTGATCCTAGTTTTCTGGGTAGGTACATGTCAGCATTAAAT
CACACTAAAAAGTGGAAATACCCACAAGTTAATGGTTTAACTTCTATTAAATGGGCAGATAACAACTGTTATCTTGCCAC
TGCATTGTTAACACTCCAACAAATAGAGTTGAAGTTTAATCCACCTGCTCTACAAGATGCTTATTACAGAGCAAGGGCTG
GTGAAGCTGCTAACTTTTGTGCACTTATCTTAGCCTACTGTAATAAGACAGTAGGTGAGTTAGGTGATGTTAGAGAAACA
ATGAGTTACTTGTTTCAACATGCCAATTTAGATTCTTGCAAAAGAGTCTTGAACGTGGTGTGTAAAACTTGTGGACAACA
GCAGACAACCCTTAAGGGTGTAGAAGCTGTTATGTACATGGGCACACTTTCTTATGAACAATTTAAGAAAGGTGTTCAGA
TACCTTGTACGTGTGGTAAACAAGCTACAAAATATCTAGTACAACAGGAGTCACCTTTTGTTATGATGTCAGCACCACCT
GCTCAGTATGAACTTAAGCATGGTACATTTACTTGTGCTAGTGAGTACACTGGTAATTACCAGTGTGGTCACTATAAACA
TATAACTTCTAAAGAAACTTTGTATTGCATAGACGGTGCTTTACTTACAAAGTCCTCAGAATACAAAGGTCCTATTACGG
ATGTTTTCTACAAAGAAAACAGTTACACAACAACCATAAAACCAGTTACTTATAAATTGGATGGTGTTGTTTGTACAGAA
ATTGACCCTAAGTTGGACAATTATTATAAGAAAGACAATTCTTATTTCACAGAGCAACCAATTGATCTTGTACCAAACCA
ACCATATCCAAACGCAAGCTTCGATAATTTTAAGTTTGTATGTGATAATATCAAATTTGCTGATGATTTAAACCAGTTAA
CTGGTTATAAGAAACCTGCTTCAAGAGAGCTTAAAGTTACATTTTTCCCTGACTTAAATGGTGATGTGGTGGCTATTGAT
TATAAACACTACACACCCTCTTTTAAGAAAGGAGCTAAATTGTTACATAAACCTATTGTTTGGCATGTTAACAATGCAAC
TAATAAAGCCACGTATAAACCAAATACCTGGTGTATACGTTGTCTTTGGAGCACAAAACCAGTTGAAACATCAAATTCGT
TTGATGTACTGAAGTCAGAGGACGCGCAGGGAATGGATAATCTTGCCTGCGAAGATCTAAAACTAGTCTCTGAAGAAGTA
GTGGAAAATCCTACCATACAGAAAGACGTTCTTGAGTGTAATGTGAAAACTACCGAAGTTGTAGGAGACATTATACTTAA
ACCAGCAAATAATAGTTTAAAAATTACAGAAGAGGTTGGCCACACAGATCTAATGGCTGCTTATGTAGACAATTCTAGTC
TTACTATTAAGAAACCTAATGAATTATCTAGAGTATTAGGTTTGAAAACCCTTGCTACTCATGGTTTAGCTGCTGTTAAT
AGTGTCCCTTGGGATACTATAGCTAATTATGCTAAGCCTTTTCTTAACAAAGTTGTTAGTACAACTACTAACATAGTTAC
ACGGTGTTTAAACCGTGTTTGTACTAATTATATGCCTTATTTCTTTACTTTATTGCTACAATTGTGTACTTTTACTAGAA
GTACAAATTCTAGAATTAAAGCATCTATGCCGACTACTATAGCAAAGAATACTGTTAAGAGTGTCGGTAAATTTTGTCTA
GAGGCTTCATTTAATTATTTGAAGTCACCTAATTTTTCTAAACTGATAAATATTATAATTTGGTTTTTACTATTAAGTGT
TTGCCTAGGTTCTTTAATCTACTCAACCGCTGCTTTAGGTGTTTTAATGTCTAATTTAGGCATGCCTTCTTACTGTACTG
GTTACAGAGAAGGCTATTTGAACTCTACTAATGTCACTATTGCAACCTACTGTACTGGTTCTATATCTTGTAGTGTTTGT
CTTAGTGGTTTAGATTCTTTAGACACCTATCCTTCTTTAGAAACTATACAAATTACCATTTCATCTTTTAAATGGGATTT
AACTGCTTTTGGCTTAGTTGCAGAGTGGTTTTTGGCATATATTCTTTTCACTAGGTTTTTCTATGTACTTGGATTGGCTG

CAATCATGCAATTGTTTTTCAGCTATTTTGCAGTACATTTTATTAGTAATTCTTGGCTTATGTGGTTAATAATTAATCTT
GTACAAATGGCCCCGATTTCAGCTATGGTTAGAATGTACATCTTCTTTGCATCATTTTATTATGTATGGAAAAGTTATGT
GCATGTTGTAGACGGTTGTAATTCATCAACTTGTATGATGTGTTACAAACGTAATAGAGCAACAAGAGTCGAATGTACAA
CTATTGTTAATGGTGTTAGAAGGTCCTTTTATGTCTATGCTAATGGAGGTAAAGGCTTTTGCAAACTACACAATTGGAAT
TGTGTTAATTGTGATACATTCTGTGCTGGTAGTACATTTATTAGTGATGAAGTTGCGAGAGACTTGTCACTACAGTTTAA
AAGACCAATAAATCCTACTGACCAGTCTTCTTACATCGTTGATAGTGTTACAGTGAAGAATGGTTCCATCCATCTTTACT
TTGATAAAGCTGGTCAAAAGACTTATGAAAGACATTCTCTCTCTCATTTTGTTAACTTAGACAACCTGAGAGCTAATAAC
ACTAAAGGTTCATTGCCTATTAATGTTATAGTTTTTGATGGTAAATCAAAATGTGAAGAATCATCTGCAAAATCAGCGTC
TGTTTACTACAGTCAGCTTATGTGTCAACCTATACTGTTACTAGATCAGGCATTAGTGTCTGATGTTGGTGATAGTGCGG
AAGTTGCAGTTAAAATGTTTGATGCTTACGTTAATACGTTTTCATCAACTTTTAACGTACCAATGGAAAAACTCAAAACA
CTAGTTGCAACTGCAGAAGCTGAACTTGCAAAGAATGTGTCCTTAGACAATGTCTTATCTACTTTTATTTCAGCAGCTCG
GCAAGGGTTTGTTGATTCAGATGTAGAAACTAAAGATGTTGTTGAATGTCTTAAATTGTCACATCAATCTGACATAGAAG
TTACTGGCGATAGTTGTAATAACTATATGCTCACCTATAACAAAGTTGAAAACATGACACCCCGTGACCTTGGTGCTTGT
ATTGACTGTAGTGCGCGTCATATTAATGCGCAGGTAGCAAAAAGTCACAACATTGCTTTGATATGGAACGTTAAAGATTT
CATGTCATTGTCTGAACAACTACGAAAACAAATACGTAGTGCTGCTAAAAAGAATAACTTACCTTTTAAGTTGACATGTG
CAACTACTAGACAAGTTGTTAATGTTGTAACAACAAAGATAGCACTTAAGGGTGGTAAAATTGTTAATAATTGGTTGAAG
CAGTTAATTAAAGTTACACTTGTGTTCCTTTTTGTTGCTGCTATTTTCTATTTAATAACACCTGTTCATGTCATGTCTAA
ACATACTGACTTTTCAAGTGAAATCATAGGATACAAGGCTATTGATGGTGGTGTCACTCGTGACATAGCATCTACAGATA
CTTGTTTTGCTAACAAACATGCTGATTTTGACACATGGTTTAGCCAGCGTGGTGGTAGTTATACTAATGACAAAGCTTGC
CCATTGATTGCTGCAGTCATAACAAGAGAAGTGGGTTTTGTCGTGCCTGGTTTGCCTGGCACGATATTACGCACAACTAA
TGGTGACTTTTTGCATTTCTTACCTAGAGTTTTTAGTGCAGTTGGTAACATCTGTTACACACCATCAAAACTTATAGAGT
ACACTGATTTTGCAACATCAGCTTGTGTTTTGGCTGCTGAATGTACAATTTTTAAAGATGCTTCTGGTAAGCCATTACCA
TATTGTTATGATACCAATGTACTAGAAGGTTCTGTTGCTTATGAAAGTTTACGCCCTGACACACGTTATGTGCTCATGGA
TGGCTCTATTATTCAATTTCCTAACACCTACCTTGAAGGTTCTGTTAGAGTGGTAACAACTTTTGATTCTGAGTACTGTA
GGCACGGCACTTGTGAAAGATCAGAAGCTGGTGTTTGTGTATCTACTAGTGGTAGATGGGTACTTAACAATGATTATTAC
AGATCTTTACCAGGAGTTTTCTGTGGTGTAGATGCTGTAAATTTACTTACTAATATGTTTACACCACTAATTCAACCTAT
TGGTGCTTTGGACATATCAGCATCTATAGTAGCTGGTGGTATTGTAGCTATCGTATTAACATGCCTTGCCTACTATTTTA
TGAGGTTTAGAAGAGCTTTTGGTGAATACAGTCATGTAGTTGCCTTTAATACTTTACTATTCCTTATGTCATTCACTGTA
CTCTGTTTAACACCAGTTTACTCATTCTTACCTGGTGTTTATTCTGTTATTTACTTGTACTTGACATTTTATCTTACTAA
TGATGTTTCTTTTTTAGCACATATTCAGTGGATGGTTATGTTCACACCTTTAGTACCTTTCTGGATAACAATTGCTTATA
TCATTTGTATTTCCACAAAGCATTTCTATTGGTTCTTTAGTAATTACCTAAAGAGACGTGTAGTCTTTAATGGTGTTTCC
TTTAGTACTTTTGAAGAAGCTGCGCTGTGCACCTTTTTGTTAAATAAAGAAATGTATCTAAAGTTGCGTAGTGATGTGCT
ATTACCTCTTACGCAATATAATAGATACTTAGCTCTTTATAATAAGTACAAGTATTTTAGTGGAGCAATGGATACAACTA
GCTACAGAGAAGCTGCTTGTTGTCATCTCGCAAAGGCTCTCAATGACTTCAGTAACTCAGGTTCTGATGTTCTTTACCAA
CCACCACAAATCTCTATCACCTCAGCTGTTTTGCAGAGTGGTTTTAGAAAAATGGCATTCCCATCTGGTAAAGTTGAGGG
TTGTATGGTACAAGTAACTTGTGGTACAACTACACTTAACGGTCTTTGGCTTGATGACGTAGTTTACTGTCCAAGACATG
TGATCTGCACCTCTGAAGACATGCTTAACCCTAATTATGAAGATTTACTCATTCGTAAGTCTAATCATAATTTCTTGGTA
CAGGCTGGTAATGTTCAACTCAGGGTTATTGGACATTCTATGCAAAATTGTGTACTTAAGCTTAAGGTTGATACAGCCAA
TCCTAAGACACCTAAGTATAAGTTTGTTCGCATTCAACCAGGACAGACTTTTTCAGTGTTAGCTTGTTACAATGGTTCAC
CATCTGGTGTTTACCAATGTGCTATGAGGCCCAATTTCACTATTAAGGGTTCATTCCTTAATGGTTCATGTGGTAGTGTT
GGTTTTAACATAGATTATGACTGTGTCTCTTTTTGTTACATGCACCATATGGAATTACCAACTGGAGTTCATGCTGGCAC
AGACTTAGAAGGTAACTTTTATGGACCTTTTGTTGACAGGCAAACAGCACAAGCAGCTGGTACGGACACAACTATTACAG
TTAATGTTTTAGCTTGGTTGTACGCTGCTGTTATAAATGGAGACAGGTGGTTTCTCAATCGATTTACCACAACTCTTAAT
GACTTTAACCTTGTGGCTATGAAGTACAATTATGAACCTCTAACACAAGACCATGTTGACATACTAGGACCTCTTTCTGC
TCAAACTGGAATTGCCGTTTTAGATATGTGTGCTTCATTAAAAGAATTACTGCAAAATGGTATGAATGGACGTACCATAT
TGGGTAGTGCTTTATTAGAAGATGAATTTACACCTTTTGATGTTGTTAGACAATGCTCAGGTGTTACTTTCCAAAGTGCA
GTGAAAAGAACAATCAAGGGTACACACCACTGGTTGTTACTCACAATTTTGACTTCACTTTTAGTTTTAGTCCAGAGTAC
TCAATGGTCTTTGTTCTTTTTTTTGTATGAAAATGCCTTTTTACCTTTTGCTATGGGTATTATTGCTATGTCTGCTTTTG
CAATGATGTTTGTCAAACATAAGCATGCATTTCTCTGTTTGTTTTTGTTACCTTCTCTTGCCGCTGTAGCTTATTTTAAT
ATGGTCTATATGCCTGCTAGTTGGGTGATGCGTATTATGACATGGTTGGATATGGTTGATACTAGTTTGTCTGGTTTTAA
GCTAAAAGACTGTGTTATGTATGCATCAGCTGTGGTGTTACTAATCCTTATGACAGCAAGAACTGTGTATGATGATGGTG
CTAGGAGAGTGTGGACACTTATGAATGTCTTGACACTCGTTTATAAAGTTTATTATGGTAATGCTTTAGATCAAGCCATT
TCCATGTGGGCTCTTATAATCTCTGTTACTTCTAACTACTCAGGTGTAGTTACAACTGTCATGTTTTTGGCCAGAGGTAT
TGTTTTTATGTGTGTTGAGTATTGCCCTATTTTCTTCATAACTGGTAATACACTTCAGTGTATAATGCTAGTTTATTGTT
TCTTAGGCTATTTTTGTACTTGTTACTTTGGCCTCTTTTGTTTACTCAACCGCTACTTTAGACTGACTCTTGGTGTTTAT
GATTACTTAGTTTCTACACAGGAGTTTAGATATATGAATTCACAGGGACTACTCCCACCCAAGAATAGCATAGATGCCTT
CAAACTCAACATTAAATTGTTGGGTGTTGGTGGCAAACCTTGTATCAAAGTAGCCACTGTACAGTCTAAAATGTCAGATG
TAAAGTGCACATCAGTAGTCTTACTCTCAGTTTTGCAACAACTCAGAGTAGAATCATCATCTAAATTGTGGGCTCAATGT
GTCCAGTTACACAATGACATTCTCTTAGCTAAAGATACTACTGAAGCCTTTGAAAAAATGGTTTCACTACTTTCTGTTTT

GCTTTCCATGCAGGGTGCTGTAGACATAAACAAGCTTTGTGAAGAAATGCTGGACAACAGGGCAACCTTACAAGCTATAG
CCTCAGAGTTTAGTTCCCTTCCATCATATGCAGCTTTTGCTACTGCTCAAGAAGCTTATGAGCAGGCTGTTGCTAATGGT
GATTCTGAAGTTGTTCTTAAAAAGTTGAAGAAGTCTTTGAATGTGGCTAAATCTGAATTTGACCGTGATGCAGCCATGCA
ACGTAAGTTGGAAAAGATGGCTGATCAAGCTATGACCCAAATGTATAAACAGGCTAGATCTGAGGACAAGAGGGCAAAAG
TTACTAGTGCTATGCAGACAATGCTTTTCACTATGCTTAGAAAGTTGGATAATGATGCACTCAACAACATTATCAACAAT
GCAAGAGATGGTTGTGTTCCCTTGAACATAATACCTCTTACAACAGCAGCCAAACTAATGGTTGTCATACCAGACTATAA
CACATATAAAAATACGTGTGATGGTACAACATTTACTTATGCATCAGCATTGTGGGAAATCCAACAGGTTGTAGATGCAG
ATAGTAAAATTGTTCAACTTAGTGAAATTAGTATGGACAATTCACCTAATTTAGCATGGCCTCTTATTGTAACAGCTTTA
AGGGCCAATTCTGCTGTCAAATTACAGAATAATGAGCTTAGTCCTGTTGCACTACGACAGATGTCTTGTGCTGCCGGTAC
TACACAAACTGCTTGCACTGATGACAATGCGTTAGCTTACTACAACACAACAAAGGGAGGTAGGTTTGTACTTGCACTGT
TATCCGATTTACAGGATTTGAAATGGGCTAGATTCCCTAAGAGTGATGGAACTGGTACTATCTATACAGAACTGGAACCA
CCTTGTAGGTTTGTTACAGACACACCTAAAGGTCCTAAAGTGAAGTATTTATACTTTATTAAAGGATTAAACAACCTAAA
TAGAGGTATGGTACTTGGTAGTTTAGCTGCCACAGTACGTCTACAAGCTGGTAATGCAACAGAAGTGCCTGCCAATTCAA
CTGTATTATCTTTCTGTGCTTTTGCTGTAGATGCTGCTAAAGCTTACAAAGATTATCTAGCTAGTGGGGGACAACCAATC
ACTAATTGTGTTAAGATGTTGTGTACACACACTGGTACTGGTCAGGCAATAACAGTTACACCGGAAGCCAATATGGATCA
AGAATCCTTTGGTGGTGCATCGTGTTGTCTGTACTGCCGTTGCCACATAGATCATCCAAATCCTAAAGGATTTTGTGACT
TAAAAGGTAAGTATGTACAAATACCTACAACTTGTGCTAATGACCCTGTGGGTTTTACACTTAAAAACACAGTCTGTACC
GTCTGCGGTATGTGGAAAGGTTATGGCTGTAGTTGTGATCAACTCCGCGAACCCATGCTTCAGTCAGCTGATGCACAATC
GTTTTTAAACGGGTTTGCGGTGTAAGTGCAGCCCGTCTTACACCGTGCGGCACAGGCACTAGTACTGATGTCGTATACAG
GGCTTTTGACATCTACAATGATAAAGTAGCTGGTTTTGCTAAATTCCTAAAAACTAATTGTTGTCGCTTCCAAGAAAAGG
ACGAAGATGACAATTTAATTGATTCTTACTTTGTAGTTAAGAGACACACTTTCTCTAACTACCAACATGAAGAAACAATT
TATAATTTACTTAAGGATTGTCCAGCTGTTGCTAAACATGACTTCTTTAAGTTTAGAATAGACGGTGACATGGTACCACA
TATATCACGTCAACGTCTTACTAAATACACAATGGCAGACCTCGTCTATGCTTTAAGGCATTTTGATGAAGGTAATTGTG
ACACATTAAAAGAAATACTTGTCACATACAATTGTTGTGATGATGATTATTTCAATAAAAAGGACTGGTATGATTTTGTA
GAAAACCCAGATATATTACGCGTATACGCCAACTTAGGTGAACGTGTACGCCAAGCTTTGTTAAAAACAGTACAATTCTG
TGATGCCATGCGAAATGCTGGTATTGTTGGTGTACTGACATTAGATAATCAAGATCTCAATGGTAACTGGTATGATTTCG
GTGATTTCATACAAACCACGCCAGGTAGTGGAGTTCCTGTTGTAGATTCTTATTATTCATTGTTAATGCCTATATTAACC
TTGACCAGGGCTTTAACTGCAGAGTCACATGTTGACACTGACTTAACAAAGCCTTACATTAAGTGGGATTTGTTAAAATA
TGACTTCACGGAAGAGAGGTTAAAACTCTTTGACCGTTATTTTAAATATTGGGATCAGACATACCACCCAAATTGTGTTA
ACTGTTTGGATGACAGATGCATTCTGCATTGTGCAAACTTTAATGTTTTATTCTCTACAGTGTTCCCACTTACAAGTTTT
GGACCACTAGTGAGAAAAATATTTGTTGATGGTGTTCCATTTGTAGTTTCAACTGGATACCACTTCAGAGAGCTAGGTGT
TGTACATAATCAGGATGTAAACTTACATAGCTCTAGACTTAGTTTTAAGGAATTACTTGTGTATGCTGCTGACCCTGCTA
TGCACGCTGCTTCTGGTAATCTATTACTAGATAAACGCACTACGTGCTTTTCAGTAGCTGCACTTACTAACAATGTTGCT
TTTCAAACTGTCAAACCCGGTAATTTTAACAAAGACTTCTATGACTTTGCTGTGTCTAAGGGTTTCTTTAAGGAAGGAAG
TTCTGTTGAATTAAAACACTTCTTCTTTGCTCAGGATGGTAATGCTGCTATCAGCGATTATGACTACTATCGTTATAATC
TACCAACAATGTGTGATATCAGACAACTACTATTTGTAGTTGAAGTTGTTGATAAGTACTTTGATTGTTACGATGGTGGC
TGTATTAATGCTAACCAAGTCATCGTCAACAACCTAGACAAATCAGCTGGTTTTCCATTTAATAAATGGGGTAAGGCTAG
ACTTTATTATGATTCAATGAGTTATGAGGATCAAGATGCACTTTTCGCATATACAAAACGTAATGTCATCCCTACTATAA
CTCAAATGAATCTTAAGTATGCCATTAGTGCAAAGAATAGAGCTCGCACCGTAGCTGGTGTCTCTATCTGTAGTACTATG
ACCAATAGACAGTTTCATCAAAAATTATTGAAATCAATAGCCGCCACTAGAGGAGCTACTGTAGTAATTGGAACAAGCAA
ATTCTATGGTGGTTGGCACAACATGTTAAAAACTGTTTATAGTGATGTAGAAAACCCTCACCTTATGGGTTGGGATTATC
CTAAATGTGATAGAGCCATGCCTAACATGCTTAGAATTATGGCCTCACTTGTTCTTGCTCGCAAACATACAACGTGTTGT
AGCTTGTCACACCGTTTCTATAGATTAGCTAATGAGTGTGCTCAAGTATTGAGTGAAATGGTCATGTGTGGCAGTTCACT
ATATGTTAAACCAGGTGGAACCTCATCAGGAGATGCCACAACTGCTTATGCTAATAGTGTTTTTAACATTTGTCAAGCTG
TCACGGCCAATGTTAATGCACTTTTATCTACTGATGGTAACAAAATTGCCGATAAGTATGTCCGCAATTTACAACACAGA
CTTTATGAGTGTCTCTATAGAAATAGAGATGTTGACACAGACTTTGTGAATGAGTTTTACGCATATTTGCGTAAACATTT
CTCAATGATGATACTCTCTGACGATGCTGTTGTGTGTTTCAATAGCACTTATGCATCTCAAGGTCTAGTGGCTAGCATAA
AGAACTTTAAGTCAGTTCTTTATTATCAAAACAATGTTTTTATGTCTGAAGCAAAATGTTGGACTGAGACTGACCTTACT
AAAGGACCTCATGAATTTTGCTCTCAACATACAATGCTAGTTAAACAGGGTGATGATTATGTGTACCTTCCTTACCCAGA
TCCATCAAGAATCCTAGGGGCCGGCTGTTTTGTAGATGATATCGTAAAAACAGATGGTACACTTATGATTGAACGGTTCG
TGTCTTTAGCTATAGATGCTTACCCACTTACTAAACATCCTAATCAGGAGTATGCTGATGTCTTTCATTTGTACTTACAA
TACATAAGAAAGCTACATGATGAGTTAACAGGACACATGTTAGACATGTATTCTGTTATGCTTACTAATGATAACACTTC
AAGGTATTGGGAACCTGAGTTTTATGAGGCTATGTACACACCGCATACAGTCTTACAGGCTGTTGGGGCTTGTGTTCTTT
GCAATTCACAGACTTCATTAAGATGTGGTGCTTGCATACGTAGACCATTCTTATGTTGTAAATGCTGTTACGACCATGTC
ATATCAACATCACATAAATTAGTCTTGTCTGTTAATCCGTATGTTTGCAATGCTCCAGGTTGTGATGTCACAGATGTGAC
TCAACTTTACTTAGGAGGTATGAGCTATTATTGTAAATCACATAAACTACCCATTAGTTTTCCATTGTGTGCTAATGGAC
AAGTTTTTGGTTTATATAAAAATACATGTGTTGGTAGCGATAATGTTACTGACTTTAATGCAATTGCAACATGTGACTGG
ACAAATGCTGGTGATTACATTTTAGCTAACACCTGTACTGAAAGACTCAAGCTTTTTGCAGCAGAAACGCTCAAAGCTAC
TGAGGAGACATTTAAACTGTCTTATGGTATTGCTACTGTACGTGAAGTGCTGTCTGACAGAGAATTACATCTTTCATGGG

AAGTTGGTAAACCTAGACCACCACTTAACCGAAATTATGTCTTTACTGGTTATCGTGTAACTAAAAACAGTAAAGTACAA
ATAGGAGAGTACACCTTTGAAAAAGGTGACTATGGTGATGCTGTTGTTTACCGAGGTACAACAACTTACAAATTAAATGT
TGGTGATTATTTTGTGCTGACATCACATACAGTAATGCCATTAAGTGCACCTACACTAGTGCCACAAGAGCACTATGTTA
GAATTACTGGCTTATACCCAACACTCAATATCTCAGATGAGTTTTCTAGCAATGTTGCAAATTATCAAAAGGTTGGTATG
CAAAAGTATTCTACACTCCAGGGACCACCTGGTACTGGTAAGAGTCATTTTGCTATTGGCCTAGCTCTCTACTACCCTTC
TGCTCGCATAGTGTATACAGCTTGCTCTCATGCCGCTGTTGATGCACTATGTGAGAAGGCATTAAAATATTTGCCTATAG
ATAAATGTAGTAGAATTATACCTGCACGTGCTCGTGTAGAGTGTTTTGATAAATTCAAAGTGAATTCAACATTAGAACAG
TATGTCTTTTGTACTGTAAATGCATTGCCTGAGACGACAGCAGATATAGTTGTCTTTGATGAAATTTCAATGGCCACAAA
TTATGATTTGAGTGTTGTCAATGCCAGATTACGTGCTAAGCACTATGTGTACATTGGCGACCCTGCTCAATTACCTGCAC
CACGCACATTGCTAACTAAGGGCACACTAGAACCAGAATATTTCAATTCAGTGTGTAGACTTATGAAAACTATAGGTCCA
GACATGTTCCTCGGAACTTGTCGGCGTTGTCCTGCTGAAATTGTTGACACTGTGAGTGCTTTGGTTTATGATAATAAGCT
TAAAGCACATAAAGACAAATCAGCTCAATGCTTTAAAATGTTTTATAAGGGTGTTATCACGCATGATGTTTCATCTGCAA
TTAACAGGCCACAAATAGGCGTGGTAAGAGAATTCCTTACACGTAACCCTGCTTGGAGAAAAGCTGTCTTTATTTCACCT
TATAATTCACAGAATGCTGTAGCCTCAAAGATTTTGGGACTACCAACTCAAACTGTTGATTCATCACAGGGCTCAGAATA
TGACTATGTCATATTCACTCAAACCACTGAAACAGCTCACTCTTGTAATGTAAACAGATTTAATGTTGCTATTACCAGAG
CAAAAGTAGGCATACTTTGCATAATGTCTGATAGAGACCTTTATGACAAGTTGCAATTTACAAGTCTTGAAATTCCACGT
AGGAATGTGGCAACTTTACAAGCTGAAAATGTAACAGGACTCTTTAAAGATTGTAGTAAGGTAATCACTGGGTTACATCC
TACACAGGCACCTACACACCTCAGTGTTGACACTAAATTCAAAACTGAAGGTTTATGTGTTGACATACCTGGCATACCTA
AGGACATGACCTATAGAAGACTCATCTCTATGATGGGTTTTAAAATGAATTATCAAGTTAATGGTTACCCTAACATGTTT
ATCACCCGCGAAGAAGCTATAAGACATGTACGTGCATGGATTGGCTTCGATGTCGAGGGGTGTCATGCTACTAGAGAAGC
TGTTGGTACCAATTTACCTTTACAGCTAGGTTTTTCTACAGGTGTTAACCTAGTTGCTGTACCTACAGGTTATGTTGATA
CACCTAATAATACAGATTTTTCCAGAGTTAGTGCTAAACCACCGCCTGGAGATCAATTTAAACACCTCATACCACTTATG
TACAAAGGACTTCCTTGGAATGTAGTGCGTATAAAGATTGTACAAATGTTAAGTGACACACTTAAAAATCTCTCTGACAG
AGTCGTATTTGTCTTATGGGCACATGGCTTTGAGTTGACATCTATGAAGTATTTTGTGAAAATAGGACCTGAGCGCACCT
GTTGTCTATGTGATAGACGTGCCACATGCTTTTCCACTGCTTCAGACACTTATGCCTGTTGGCATCATTCTATTGGATTT
GATTACGTCTATAATCCGTTTATGATTGATGTTCAACAATGGGGTTTTACAGGTAACCTACAAAGCAACCATGATCTGTA
TTGTCAAGTCCATGGTAATGCACATGTAGCTAGTTGTGATGCAATCATGACTAGGTGTCTAGCTGTCCACGAGTGCTTTG
TTAAGCGTGTTGACTGGACTATTGAATATCCTATAATTGGTGATGAACTGAAGATTAATGCGGCTTGTAGAAAGGTTCAA
CACATGGTTGTTAAAGCTGCATTATTAGCAGACAAATTCCCAGTTCTTCACGACATTGGTAACCCTAAAGCTATTAAGTG
TGTACCTCAAGCTGATGTAGAATGGAAGTTCTATGATGCACAGCCTTGTAGTGACAAAGCTTATAAAATAGAAGAATTAT
TCTATTCTTATGCCACACATTCTGACAAATTCACAGATGGTGTATGCCTATTTTGGAATTGCAATGTCGATAGATATCCT
GTTAATTCCATTGTTTGTAGATTTGACACTAGAGTGCTATCTAACCTTAACTTGCCTGGTTGTGATGGTGGCAGTTTGTA
TGTAAATAAACATGCATTCCACACACCAGCTTTTGATAAAAGTGCTTTTGTTAATTTAAAACAATTACCATTTTTCTATT
ACTCTGACAGTCCATGTGAGTCTCATGGAAAACAAGTAGTGTCAGATATAGATTATGTACCACTAAAGTCTGCTACGTGT
ATAACACGTTGCAATTTAGGTGGTGCTGTCTGTAGACATCATGCTAATGAGTACAGATTGTATCTCGATGCTTATAACAT
GATGATCTCAGCTGGCTTTAGCTTGTGGGTTTACAAACAATTTGATACTTATAACCTCTGGAACACTTTTACAAGACTTC
AGAGTTTAGAAAATGTGGCTTTTAATGTTGTAAATAAGGGACACTTTGATGGACAACAGGGTGAAGTACCAGTTTCTATC
ATTAATAACACTGTTTACACAAAAGTTGATGGTGTTGATGTAGAATTGTTTGAAAATAAAACAACATTACCTGTTAATGT
AGCATTTGAGCTTTGGGCTAAGCGCAACATTAAACCAGTACCAGAGGTGAAAATACTCAATAATTTGGGTGTGGACATTG
CTGCTAATACTGTGATCTGGGACTACAAAAGAGATGCTCCAGCACATATATCTACTATTGGTGTTTGTTCTATGACTGAC
ATAGCCAAGAAACCAACTGAAACGATTTGTGCACCACTCACTGTCTTTTTTGATGGTAGAGTTGATGGTCAAGTAGACTT
ATTTAGAAATGCCCGTAATGGTGTTCTTATTACAGAAGGTAGTGTTAAAGGTTTACAACCATCTGTAGGTCCCAAACAAG
CTAGTCTTAATGGAGTCACATTAATTGGAGAAGCCGTAAAAACACAGTTCAATTATTATAAGAAAGTTGATGGTGTTGTC
CAACAATTACCTGAAACTTACTTTACTCAGAGTAGAAATTTACAAGAATTTAAACCCAGGAGTCAAATGGAAATTGATTT
CTTAGAATTAGCTATGGATGAATTCATTGAACGGTATAAATTAGAAGGCTATGCCTTCGAACATATCGTTTATGGAGATT
TTAGTCATAGTCAGTTAGGTGGTTTACATCTACTGATTGGACTAGCTAAACGTTTTAAGGAATCACCTTTTGAATTAGAA
GATTTTATTCCTATGGACAGTACAGTTAAAAACTATTTCATAACAGATGCGCAAACAGGTTCATCTAAGTGTGTGTGTTC
TGTTATTGATTTATTACTTGATGATTTTGTTGAAATAATAAAATCCCAAGATTTATCTGTAGTTTCTAAGGTTGTCAAAG
TGACTATTGACTATACAGAAATTTCATTTATGCTTTGGTGTAAAGATGGCCATGTAGAAACATTTTACCCAAAATTACAA
TCTAGTCAAGCGTGGCAACCGGGTGTTGCTATGCCTAATCTTTACAAAATGCAAAGAATGCTATTAGAAAAGTGTGACCT
TCAAAATTATGGTGATAGTGCAACATTACCTAAAGGCATAATGATGAATGTCGCAAAATATACTCAACTGTGTCAATATT
TAAACACATTAACATTAGCTGTACCCTATAATATGAGAGTTATACATTTTGGTGCTGGTTCTGATAAAGGAGTTGCACCA
GGTACAGCTGTTTTAAGACAGTGGTTGCCTACGGGTACGCTGCTTGTCGATTCAGATCTTAATGACTTTGTCTCTGATGC
AGATTCAACTTTGATTGGTGATTGTGCAACTGTACATACAGCTAATAAATGGGATCTCATTATTAGTGATATGTACGACC
CTAAGACTAAAAATGTTACAAAAGAAAATGACTCTAAAGAGGGTTTTTTCACTTACATTTGTGGGTTTATACAACAAAAG
CTAGCTCTTGGAGGTTCCGTGGCTATAAAGATAACAGAACATTCTTGGAATGCTGATCTTTATAAGCTCATGGGACACTT
CGCATGGTGGACAGCCTTTGTTACTAATGTGAATGCGTCATCATCTGAAGCATTTTTAATTGGATGTAATTATCTTGGCA
AACCACGCGAACAAATAGATGGTTATGTCATGCATGCAAATTACATATTTTGGAGGAATACAAATCCAATTCAGTTGTCT
TCCTATTCTTTATTTGACATGAGTAAATTTCCCCTTAAATTAAGGGGTACTGCTGTTATGTCTTTAAAAGAAGGTCAAAT

CAATGATATGATTTTATCTCTTCTTAGTAAAGGTAGACTTATAATTAGAGAAAACAACAGAGTTGTTATTTCTAGTGATG
TTCTTGTTAACAACTAAACGAACAATGTTTGTTTTTCTTGTTTTATTGCCACTAGTCTCTAGTCAGTGTGTTAATCTTAG
AACCAGAACTCAATTACCCCCTGCATACACTAATTCTTTCACACGTGGTGTTTATTACCCTGACAAAGTTTTCAGATCCT
CAGTTTTACATTCAACTCAGGACTTGTTCTTACCTTTCTTTTCCAATGTTACTTGGTTCCATGCTATACATGTCTCTGGG
ACCAATGGTACTAAGAGGTTTGATAACCCTGTCCTACCATTTAATGATGGTGTTTATTTTGCTTCCATTGAGAAGTCTAA
CATAATAAGAGGCTGGATTTTTGGTACTACTTTAGATTCGAAGACCCAGTCCCTACTTATTGTTAATAACGCTACTAATG
TTGTTATTAAAGTCTGTGAATTTCAATTTTGTAATGATCCATTTTTGGATGTTTATTACCACAAAAACAACAAAAGTTGG
ATGGAAAGTGGAGTTTATTCTAGTGCGAATAATTGCACTTTTGAATATGTCTCTCAGCCTTTTCTTATGGACCTTGAAGG
AAAACAGGGTAATTTCAAAAATCTTAGGGAATTTGTGTTTAAGAATATTGATGGTTATTTTAAAATATATTCTAAGCACA
CGCCTATTAATTTAGTGCGTGATCTCCCTCAGGGTTTTTCGGCTTTAGAACCATTGGTAGATTTGCCAATAGGTATTAAC
ATCACTAGGTTTCAAACTTTACTTGCTTTACATAGAAGTTATTTGACTCCTGGTGATTCTTCTTCAGGTTGGACAGCTGG
TGCTGCAGCTTATTATGTGGGTTATCTTCAACCTAGGACTTTTCTATTAAAATATAATGAAAATGGAACCATTACAGATG
CTGTAGACTGTGCACTTGACCCTCTCTCAGAAACAAAGTGTACGTTGAAATCCTTCACTGTAGAAAAAGGAATCTATCAA
ACTTCTAACTTTAGAGTCCAACCAACAGAATCTATTGTTAGATTTCCTAATATTACAAACTTGTGCCCTTTTGGTGAAGT
TTTTAACGCCACCAGATTTGCATCTGTTTATGCTTGGAACAGGAAGAGAATCAGCAACTGTGTTGCTGATTATTCTGTCC
TATATAATTCCGCATCATTTTCCACTTTTAAGTGTTATGGAGTGTCTCCTACTAAATTAAATGATCTCTGCTTTACTAAT
GTCTATGCAGATTCATTTGTAATTAGAGGTGATGAAGTCAGACAAATCGCTCCAGGGCAAACTGGAAAGATTGCTGATTA
TAATTATAAATTACCAGATGATTTTACAGGCTGCGTTATAGCTTGGAATTCTAACAATCTTGATTCTAAGGTTGGTGGTA
ATTATAATTACCGGTATAGATTGTTTAGGAAGTCTAATCTCAAACCTTTTGAGAGAGATATTTCAACTGAAATCTATCAG
GCCGGTAGCAAACCTTGTAATGGTGTTGAAGGTTTTAATTGTTACTTTCCTTTACAATCATATGGTTTCCAACCCACTAA
TGGTGTTGGTTACCAACCATACAGAGTAGTAGTACTTTCTTTTGAACTTCTACATGCACCAGCAACTGTTTGTGGACCTA
AAAAGTCTACTAATTTGGTTAAAAACAAATGTGTCAATTTCAACTTCAATGGTTTAACAGGCACAGGTGTTCTTACTGAG
TCTAACAAAAAGTTTCTGCCTTTCCAACAATTTGGCAGAGACATTGCTGACACTACTGATGCTGTCCGTGATCCACAGAC
ACTTGAGATTCTTGACATTACACCATGTTCTTTTGGTGGTGTCAGTGTTATAACACCAGGAACAAATACTTCTAACCAGG
TTGCTGTTCTTTATCAGGGTGTTAACTGCACAGAAGTCCCTGTTGCTATTCATGCAGATCAACTTACTCCTACTTGGCGT
GTTTATTCTACAGGTTCTAATGTTTTTCAAACACGTGCAGGCTGTTTAATAGGGGCTGAACATGTCAACAACTCATATGA
GTGTGACATACCCATTGGTGCAGGTATATGCGCTAGTTATCAGACTCAGACTAATTCTCGTCGGCGGGCACGTAGTGTAG
CTAGTCAATCCATCATTGCCTACACTATGTCACTTGGTGCAGAAAATTCAGTTGCTTACTCTAATAACTCTATTGCCATA
CCCACAAATTTTACTATTAGTGTTACCACAGAAATTCTACCAGTGTCTATGACCAAGACATCAGTAGATTGTACAATGTA
CATTTGTGGTGATTCAACTGAATGCAGCAATCTTTTGTTGCAATATGGCAGTTTTTGTACACAATTAAACCGTGCTTTAA
CTGGAATAGCTGTTGAACAAGACAAAAACACCCAAGAAGTTTTTGCACAAGTCAAACAAATTTACAAAACACCACCAATT
AAAGATTTTGGTGGTTTTAATTTTTCACAAATATTACCAGATCCATCAAAACCAAGCAAGAGGTCATTTATTGAAGATCT
ACTTTTCAACAAAGTGACACTTGCAGATGCTGGCTTCATCAAACAATATGGTGATTGCCTTGGTGATATTGCTGCTAGAG
ACCTCATTTGTGCACAAAAGTTTAACGGCCTTACTGTTTTGCCACCTTTGCTCACAGATGAAATGATTGCTCAATACACT
TCTGCACTGTTAGCGGGTACAATCACTTCTGGTTGGACCTTTGGTGCAGGTGCTGCATTACAAATACCATTTGCTATGCA
AATGGCTTATAGGTTTAATGGTATTGGAGTTACACAGAATGTTCTCTATGAGAACCAAAAATTGATTGCCAACCAATTTA
ATAGTGCTATTGGCAAAATTCAAGACTCACTTTCTTCCACAGCAAGTGCACTTGGAAAACTTCAAAATGTGGTCAACCAA
AATGCACAAGCTTTAAACACGCTTGTTAAACAACTTAGCTCCAATTTTGGTGCAATTTCAAGTGTTTTAAATGATATCCT
TTCACGTCTTGACAAAGTTGAGGCTGAAGTGCAAATTGATAGGTTGATCACAGGCAGACTTCAAAGTTTGCAGACATATG
TGACTCAACAATTAATTAGAGCTGCAGAAATCAGAGCTTCTGCTAATCTTGCTGCTACTAAAATGTCAGAGTGTGTACTT
GGACAATCAAAAAGAGTTGATTTTTGTGGAAAGGGCTATCATCTTATGTCCTTCCCTCAGTCAGCACCTCATGGTGTAGT
CTTCTTGCATGTGACTTATGTCCCTGCACAAGAAAAGAACTTCACAACTGCTCCTGCCATTTGTCATGATGGAAAAGCAC
ACTTTCCTCGTGAAGGTGTCTTTGTTTCAAATGGCACACACTGGTTTGTAACACAAAGGAATTTTTATGAACCACAAATC
ATTACTACAGACAACACATTTGTGTCTGGTAACTGTGATGTTGTAATAGGAATTGTCAACAACACAGTTTATGATCCTTT
GCAACCTGAATTAGACTCATTCAAGGAGGAGTTAGATAAATATTTTAAGAATCATACATCACCAGATGTTGATTTAGGTG
ACATCTCTGGCATTAATGCTTCAGTTGTAAACATTCAAAAAGAAATTGACCGCCTCAATGAGGTTGCCAAGAATTTAAAT
GAATCTCTCATCGATCTCCAAGAACTTGGAAAGTATGAGCAGTATATAAAATGGCCATGGTACATTTGGCTAGGTTTTAT
AGCTGGCTTGATTGCCATAGTAATGGTGACAATTATGCTTTGCTGTATGACCAGTTGCTGTAGTTGTCTCAAGGGCTGTT
GTTCTTGTGGATCCTGCTGCAAATTTGATGAAGACGACTCTGAGCCAGTGCTCAAAGGAGTCAAATTACATTACACATAA
ACGAACTTATGGATTTGTTTATGAGAATCTTCACAATTGGAACTGTAACTTTGAAGCAAGGTGAAATCAAGGATGCTACT
CCTTTAGATTTTGTTCGCGCTACTGCAACGATACCGATACAAGCCTCACTCCCTTTCGGATGGCTTATTGTTGGCGTTGC
ACTTCTTGCTGTTTTTCAGAGCGCTTCCAAAATCATAACCCTCAAAAAGAGATGGCAACTAGCACTCTCCAAGGGTGTTC
ACTTTGTTTGCAACTTGCTGTTGTTGTTTGTAACAGTTTACTCACACCTTTTGCTCGTTGCTGCTGGCCTTGAAGCCCCT
TTTCTCTATCTTTATGCTTTAGTCTACTTCTTGCAGAGTATAAACTTTGTAAGAATAATAATGAGGCTTTGGCTTTGCTG
GAAATGCCGTTCCAAAAACCCATTACTTTATGATGCCAACTATTTTCTTTGCTGGCATACTAATTGTTACGACTATTGTA
TACCTTACAATAGTGTAACTTCTTCAATTGTCATTACTTCAGGTGATGGCACAACAAGTCCTATTTCTGAACATGACTAC
CAGATTGGTGGTTATACTGAAAAATGGGAATCTGGAGTAAAAGACTGTGTTGTATTACACAGTTACTTCACTTCAGACTA
TTACCAGCTGTACTCAACTCAATTGAGTACAGACACTGGTGTTGAACATGTTACCTTCTTCATCTACAATAAAATTGTTG
ATGAGCCTGAAGAACATGTCCAAATTCACACAATCGACGGTTCATCCGGAGTTGTTAATCCAGTAATGGAACCAATTTAT

GATGAACCGACGACGACTACTAGCGTGCCTTTGTAAGCACAAGCTGATGAGTACGAACTTATGTACTCATTCGTTTCGGA
AGAGACAGGTACGTTAATAGTTAATAGCGTACTTCTTTTTCTTGCTTTCGTGGTATTCTTGCTAGTTACACTAGCCATCC
TTACTGCGCTTCGATTGTGTGCGTACTGCTGCAATATTGTTAACGTGAGTCTTGTAAAACCTTCTTTTTACGTTTACTCT
CGTGTTAAAAATCTGAATTCTTCTAGAGTTCCTGATCTTCTGGTCTAAACGAACTAAATATTATATTAGTTTTTCTGTTT
GGAACTTTAATTTTAGCCATGGCAGATTCCAACGGTACTATTACCGTTGAAGAGCTTAAAAAGCTCCTTGAACAATGGAA
CCTAGTAATAGGTTTCCTATTCCTTACATGGATTTGTCTTCTACAATTTGCCTATGCCAACAGGAATAGGTTTTTGTATA
TAATTAAGTTAATTTTCCTCTGGCTGTTATGGCCAGTAACTTTAGCTTGTTTTGTGCTTGCTGCTGTTTACAGAATAAAT
TGGATCACCGGTGGAATTGCTACCGCAATGGCTTGTCTTGTAGGCTTGATGTGGCTCAGCTACTTCATTGCTTCTTTCAG
ACTGTTTGCGCGTACGCGTTCCATGTGGTCATTCAATCCAGAAACTAACATTCTTCTCAACGTGCCACTCCATGGCACTA
TTCTGACCAGACCGCTTCTAGAAAGTGAACTCGTAATCGGAGCTGTGATCCTTCGTGGACATCTTCGTATTGCTGGACAC
CATCTAGGACGCTGTGACATCAAGGACCTGCCTAAAGAAATCACTGTTGCTACATCACGAACGCTTTCTTATTACAAATT
GGGAGCTTCGCAGCGTGTAGCAGGTGACTCAGGTTTTGCTGCATACAGTCGCTACAGGATTGGCAACTATAAATTAAACA
CAGACCATTCCAGTAGCAGTGACAATATTGCTTTGCTTGTACAGTAAGTGACAACAGATGTTTCATCTCGTTGACTTTCA
GGTTACTATAGCAGAGATATTACTAATTATTATGAGGACTTTTAAAGTTTCCATTTGGAATCTTGATTACATCATAAACC
TCATAATTAAAAATTTATCTAAGTCACTAACTGAGAATAAATATTCTCAATTAGATGAAGAGCAACCAATGGAGATTGAT
TAAACGAACATGAAAATTATTCTTTTCTTGGCACTGATAACACTCGCTACTTGTGAGCTTTATCACTACCAAGAGTGTGT
TAGAGGTACAACAGTACTTTTAAAAGAACCTTGCTCTTCTGGAACATACGAGGGCAATTCACCATTTCATCCTCTAGCTG
ATAACAAATTTGCACTGACTTGCTTTAGCACTCAATTTGCTTTTGCTTGTCCTGACGGCGTAAAACACGTCTATCAGTTA
CGTGCCAGATCAGCTTCACCTAAACTGTTCATCAGACAAGAGGAAGTTCAAGAACTTTACTCTCCAATTTTTCTTATTGT
TGCGGCAATAGTGTTTATAACACTTTGCTTCACACTCAAAAGAAAGATAGAATGATTGAACTTTCATTAATTGACTTCTA
TTTGTGCTTTTTAGCCTTTCTGCTATTCCTTGTTTTAATTATGCTTATTATCTTTTGGTTCTCACTTGAACTGCAAGATC
ATAATGAAATTTGTCACGCCTAAACGAACATGAAATTTCTTGTTTTCTTAGGAATCATCACAACTGTAGCTGCATTTCAC
CAAGAATGTAGTTTACAGTCATGTACTCAACATCAACCATATGTAGTTGATGACCCGTGTCCTATTCACTTCTATTCTAA
ATGGTATATTAGAGTAGGAGCTAGAAAATCAGCACCTTTAATTGAATTGTGCGTGGATGAGGCTGGTTCTAAATCACCCA
TTCAGTACATCGATATCGGTAATTATACAGTTTCCTGTTTACCTTTTACAATTAATTGCCAGGAACCTAAATTGGGTAGT
CTTGTAGTGCGTTGTTCGTTCTATGAAGACTTTTTAGAGTATCATGACGTTCGTGTTGTTTTAATCTAAACGAACAAACT
AAATGTCTGATAATGGACCCCAAAATCAGCGAAATGCACCCCGCATTACGTTTGGTGGACCCTCAGATTCAACTGGCAGT
AACCAGAATGGAGAACGCAGTGGGGCGCGATCAAAACAACGTCGGCCCCAAGGTTTACCCAATAATACTGCGTCTTGGTT
CACCGCTCTCACTCAACATGGCAAGGAAGGCCTTAAATTCCCTCGAGGACAAGGCGTTCCAATTAACACCAATAGCAGTC
CAGATGACCAAATTGGCTACTACCGAAGAGCTACCAGACGAATTCGTGGTGGTGACGGTAAAATGAAAGATCTCAGTCCA
AGATGGTATTTCTACTACCTAGGAACTGGGCCAGAAGCTGGACTTCCCTATGGTGCTAACAAAGACGGCATCATATGGGT
TGCAACTGAGGGAGCCTTGAATACACCAAAAGATCACATTGGCACCCGCAATCTTGCTAACAATGCTGCAATCGTGCTAC
AACTTCCTCAAGGAACAACATTGCCAAAAGGCTTCTACGCAGAAGGGAGCAGAGGCGGCAGTCAAGCCTCTTCTCGTTCC
TCATCACGTAGTCGCAACAGTTCAAGAAATTCAACTCCAGGCAGCAGTATGGGAACTTCTCCTGCTAGAATGGCTGGCAA
TGGCTGTGATGCTGCTCTTGCTTTGCTGCTGCTTGACAGATTGAACCAGCTTGAGAGCAAAATGTCTGGTAAAGGCCAAC
AACAACAAGGCCAAACTGTCACTAAGAAATCTGCTGCTGAGGCTTCTAAGAAGCCTCGGCAAAAACGTACTGCCACTAAA
GCATACAATGTAACACAAGCTTTCGGCAGACGTGGTCCAGAACAAACCCAAGGAAATTTTGGGGACCAGGAACTAATCAG
ACAAGGAACTGATTACAAACATTGGCCGCAAATTGCACAATTTGCCCCCAGCGCTTCAGCGTTCTTCGGAATGTCGCGCA
TTGGCATGGAAGTCACACCTTCGGGAACGTGGTTGACCTACACAGGTGCCATCAAATTGGATGACAAAGATCCAAATTTC
AAAGATCAAGTCATTTTGCTGAATAAGCATATTGACGCATACAAAACATTCCCACCAACAGAGCCTAAAAAGGACAAAAA
GAAGAAGGCTTATGAAACTCAAGCCTTACCGCAGAGACAGAAGAAACAGCAAACTGTGACTCTTCTTCCTGCTGCAGATT
TGGATGATTTCTCCAAACAATTGCAACAATCCATGAGCAGTGCTGACTCAACTCAGGCCTAAACTCATGCAGACCACACA
AGGCAGATGGGCTATATAAACGTTTTCGCTTTTCCGTTTACGATATATAGTCTACTCTTGTGCAGAATGAATTCTCGTAA
CTACATAGCACAAGTAGATGTAGTTAACTTTAATCTCACATAGCAATCTTTAATCAGTGTGTAACATTAGGGAGGACTTG
AAAGAGCCACCACATTTTCACCGAGGCCACGCGGAGTACGATCGAGTGTACAGTGAACAATGCTAGGGAGAGCTGCCTAT
ATGGAAGAGCCCTAATGTGTAAAATTAATTTTAGTAGTGCTATCCCCATGTGATTTTAATAGCTTCTTAGGAGAATGACA
AAAAAAAAAAAAAAAAAAAAAAAAAAAAACGGCCGGGCCGGCATGGTCCCAGCCTCCTCGCTGGCGCCGGCTGGGCAACAT
TCCGAGGGGACCGTCCCCTCGGTAATGGCGAATGGGACGGGCCCTGCGATATCGCGACGAGGATCTAGATCCTCTAGAGT
CGACCTGCAGGCATGCAAGCTTGAGTATTCTATAGTCTCACCTAAATAGCTTGGCGTAATCATGGTCATAGCTGTTTCCT
GTGTGAAATTGTTATCCGCTCACAATTCCACACAACATACGAGCCGGAAGCATAAAGTGTAAAGCCTGGGGTGCCTAATG
AGTGAGCTAACTCACATTAATTGCGTTGCGCTCACTGCCCGCTTTCCAGTCGGGAAACCTGTCGTGCCAGCTGCATTAAT
GAATCGGCCAACGCGAACCCCTTGCGGCCGCCCGGGCCGTCGACCAATTCTCATGTTTGACAGCTTATCATCGAATTTCT
GCCATTCATCCGCTTATTATCACTTATTCAGGCGTAGCAACCAGGCGTTTAAGGGCACCAATAACTGCCTTAAAAAAATT
ACGCCCCGCCCTGCCACTCATCGCAGTACTGTTGTAATTCATTAAGCATTCTGCCGACATGGAAGCCATCACAAACGGCA
TGATGAACCTGAATCGCCAGCGGCATCAGCACCTTGTCGCCTTGCGTATAATATTTGCCCATGGTGAAAACGGGGGCGAA
GAAGTTGTCCATATTGGCCACGTTTAAATCAAAACTGGTGAAACTCACCCAGGGATTGGCTGAGACGAAAAACATATTCT
CAATAAACCCTTTAGGGAAATAGGCCAGGTTTTCACCGTAACACGCCACATCTTGCGAATATATGTGTAGAAACTGCCGG
AAATCGTCGTGGTATTCACTCCAGAGCGATGAAAACGTTTCAGTTTGCTCATGGAAAACGGTGTAACAAGGGTGAACACT
ATCCCATATCACCAGCTCACCGTCTTTCATTGCCATACGAAATTCCGGATGAGCATTCATCAGGCGGGCAAGAATGTGAA

TAAAGGCCGGATAAAACTTGTGCTTATTTTTCTTTACGGTCTTTAAAAAGGCCGTAATATCCAGCTGAACGGTCTGGTTA
TAGGTACATTGAGCAACTGACTGAAATGCCTCAAAATGTTCTTTACGATGCCATTGGGATATATCAACGGTGGTATATCC
AGTGATTTTTTTCTCCATTTTAGCTTCCTTAGCTCCTGAAAATCTCGATAACTCAAAAAATACGCCCGGTAGTGATCTTA
TTTCATTATGGTGAAAGTTGGAACCTCTTACGTGCCGATCAACGTCTCATTTTCGCCAAAAGTTGGCCCAGGGCTTCCCG
GTATCAACAGGGACACCAGGATTTATTTATTCTGCGAAGTGATCTTCCGTCACAGGTATTTATTCGCGATAAGCTCATGG
AGCGGCGTAACCGTCGCACAGGAAGGACAGAGAAAGCGCGGATCTGGGAAGTGACGGACAGAACGGTCAGGACCTGGATT
GGGGAGGCGGTTGCCGCCGCTGCTGCTGACGGTGTGACGTTCTCTGTTCCGGTCACACCACATACGTTCCGCCATTCCTA
TGCGATGCACATGCTGTATGCCGGTATACCGCTGAAAGTTCTGCAAAGCCTGATGGGACATAAGTCCATCAGTTCAACGG
AAGTCTACACGAAGGTTTTTGCGCTGGATGTGGCTGCCCGGCACCGGGTGCAGTTTGCGATGCCGGAGTCTGATGCGGTT
GCGATGCTGAAACAATTATCCTGAGAATAAATGCCTTGGCCTTTATATGGAAATGTGGAACTGAGTGGATATGCTGTTTT
TGTCTGTTAAACAGAGAAGCTGGCTGTTATCCACTGAGAAGCGAACGAAACAGTCGGGAAAATCTCCCATTATCGTAGAG
ATCCGCATTATTAATCTCAGGAGCCTGTGTAGCGTTTATAGGAAGTAGTGTTCTGTCATGATGCCTGCAAGCGGTAACGA
AAACGATTTGAATATGCCTTCAGGAACAATAGAAATCTTCGTGCGGTGTTACGTTGAAGTGGAGCGGATTATGTCAGCAA
TGGACAGAACAACCTAATGAACACAGAACCATGATGTGGTCTGTCCTTTTACAGCCAGTAGTGCTCGCCGCAGTCGAGCG
ACAGGGCGAAGCCCTCGAGGG
