## Supplementary material for "NSP4 mutation T492I drives rapid evolution of SARS-CoV-2 toward Omicron": Data S1: Tables.pdf

Table 1. Mutagenesis primers used in the generation of SARS-CoV-2 mutants

| Primer | Sequence (5'-3') |
|---|---|
| nsp4 T492I F | AGTAACTCAGGTTCTGATGTTCTTTACCAACCACCACAAATCTCTATCACCTCAGCTGTTTAGGGATAACAGGGTAATCG |
| nsp4 T492I R | TTTTCTAAAACCACTCTGCAAAACAGCTGAGGTGATAGAGATTTGTGGTGGTTGGTAAAGAGCCAGTGTTACAACCAATTAAC |
| nsp4 I492T F | AGTAACTCAGGTTCTGATGTTCTTTACCAACCACCACAAACCTCTATCACCTCAGCTGTTTAGGGATAACAGGGTAATCG |
| nsp4 I492T R | TTTTCTAAAACCACTCTGCAAAACAGCTGAGGTGATAGAGGTTTGTGGTGGTTGGTAAAGAGCCAGTGTTACAACCAATTAAC |

Table 2. Primers used for the amplification of the overlapping fragments of SARS-CoV-2 Delta.

| Name | Sequence (5'-3') |
|---|---|
| Delta-F1-F | CTTCCCAGGTAACAAACC |
| Delta-F1-R | CGTTTGATGAACACATAGGGCTG |
| Delta-F2-F | GTCTTATCAGAGGCACGTCAAC |
| Delta-F2-R | CTGGTGTAAGTTCCATCTCTAATTG |
| Delta-F3-F | CAGATGAGGATGAAGAAGAAGG |
| Delta-F3-R | TTTGTGCTCCAAAGACAACGTATAC |
| Delta-F4-F | ATCTTGTACCAAACCAACCATATCC |
| Delta-F4-R | GTTGCAAAGTCAGTGTACTCTATAAG |
| Delta-F5-F | GTGACATAGCATCTACAGATACTTG |
| Delta-F5-R | CTAAGAGAATGTCATTGTGTAACTGG |
| Delta-F6-F | TCAACCGCTACTTTAGACTGAC |
| Delta-F6-R | AATAGATTACCAGAAGCAGCGTG |
| Delta-F7-F | GATCAGACATACCACCCAAATTG |
| Delta-F7-R | TTGCAGATGAAACATCATGCGTG |
| Delta-F8-F | ATGCCAGATTACGTGCTAAGCAC |
| Delta-F8-R | ACCTAACTGACTATGACTAAAATCTC |
| Delta-F9-F | GGAGTCACATTAATTGGAGAAGC |
| Delta-F9-R | GCATCAGTAGTGTCAGCAATGTC |
| Delta-F10-F | AATCTATCAGGCCGGTAGCAC |
| Delta-F10-R | TCATGTTCAGAAATAGGACTTGTTG |
| Delta-F11-F | GATGGCAACTAGCACTCTCC |
| Delta-F11-R | TTTGGCAATGTTGTTCCTTGAGG |
| Delta-F12-F | ATGTCTGATAATGGACCCCAAAATC |
| Delta-F12-R | TTTTTTGTCATTCTCCTAAGAAGCT |

Table 3. Primers that target envelope protein (E) gene and Orf1ab sequences.

| Target | Sequence |
|---|---|
| E_Sarbeco_F | 5’-ACAGGTACGTTAATAGTTAATAGCGT-3’ |
| E_Sarbeco_R | 5’-ATATTGCAGCAGTACGCACACA-3’ |
| E_Sarbeco_P1 (Probe) | 5’-FAM-ACACTAGCCATCCTTACTGCGCTTCG-BHQ1-3’ |
| SARS-CoV2.ORF1ab.F | 5’-GGCCAATTCTGCTGTCAAATTA-3’ |
| SARS-CoV2.ORF1ab.R | 5’-CAGTGCAAGCAGTTTGTGTAG-3’ |
| SARS-CoV2.ORF1ab.P (Probe) | 5’-FAM-ACAGATGTCTTGTGCTGCCGGTA-BHQ1-3’ |

Table 4. Primers used in qRT-PCR.

| Target | Sequence |
|---|---|
| IFN-β | Forward, 5'-TAGCACTGGCTGGAATGAG-3',<br>Reverse, 5'-GTTTCGGAGGTAACCTGTAAG-3' |
| IFN-λ | Forward, 5'-TCGCTTCTGCTGAAGGACTGCA-3',<br>Reverse, 5'-CCTCCAGAACCTTCAGCGTCAG-3' |
| ISG56 | Forward, 5'-TACAGCAACCATGAGTACAA-3',<br>Reverse, 5'-TCAGGTGTTTCACATAGGC-3' |
| GAPDH | Forward, 5'-GCAAATTTCCATGGCACCGT-3',<br>Reverse, 5'-GCCCCACTTGATTTTGGAGG-3' |

Table 5. Primers used for the analyses of the expression of APOBEC and ADAR enzymes.

| Target | Sequence |
|---|---|
| APOBEC3A_F | 5’-TGGCATTGGAAGGCATAAGAC-3’ |
| APOBEC3A_R | 5’-TTAGCCTGGTTGTGTAGAAAGC-3’ |
| APOBEC1_F | 5’-GTGGATGATGTTGTACGCACT-3’ |
| APOBEC1_R | 5’-GCGGAATCGTTTGGTAATGGC-3’ |
| APOBEC3G_F | 5’-GCATCGTGACCAGGAGTATGA-3’ |
| APOBEC3G_R | 5’-GTCAGGGTAACCTTCGGGT-3’ |
| ADAR_F | 5’-CTGAGACCAAAAGAAACGCAGA-3’ |
| ADAR_R | 5’-GCCATTGTAATGAACAGGTGGTT-3’ |
| PRORP (control)_F | 5’- GACACAGTGGTGCAAACAACT-3’ |
| PRORP (control)_R | 5’- CATTTGGTTGGTACTTCACAGGA-3’ |
