## Supplementary figures and images for "NSP4 mutation T492I drives rapid evolution of SARS-CoV-2 toward Omicron"

### Fig 1.jpg

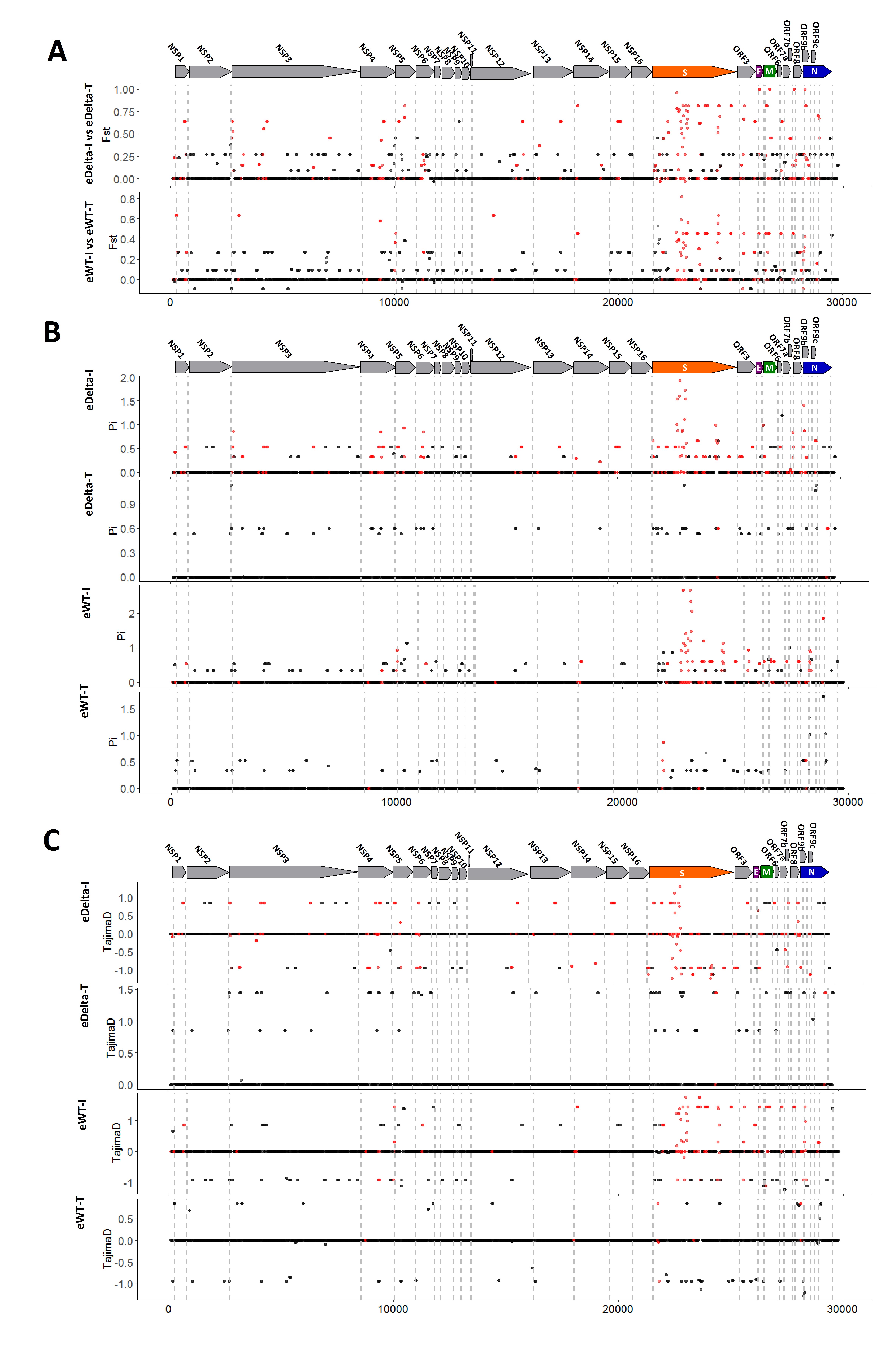

### Fig 2.jpg

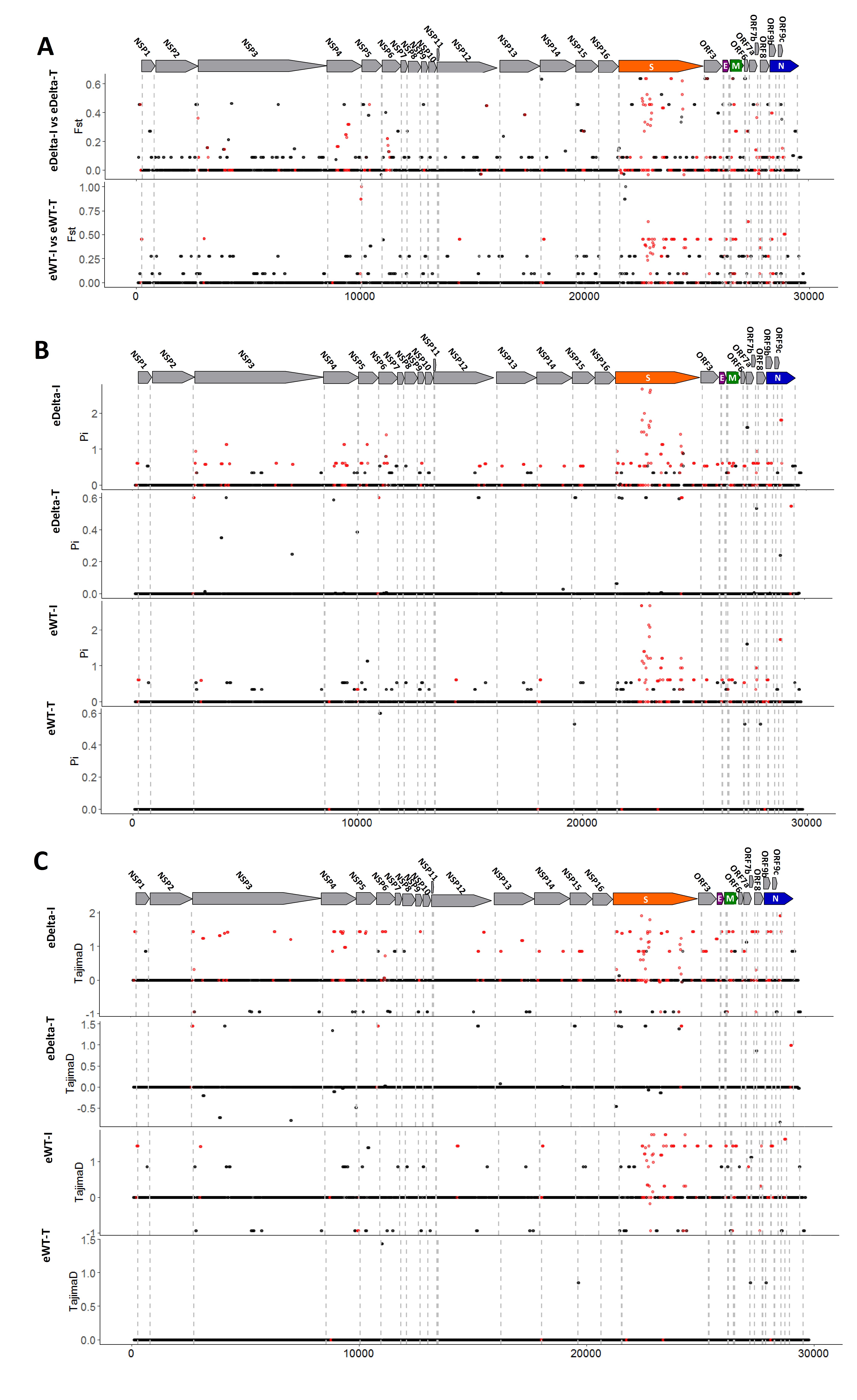

### Fig 3.jpg

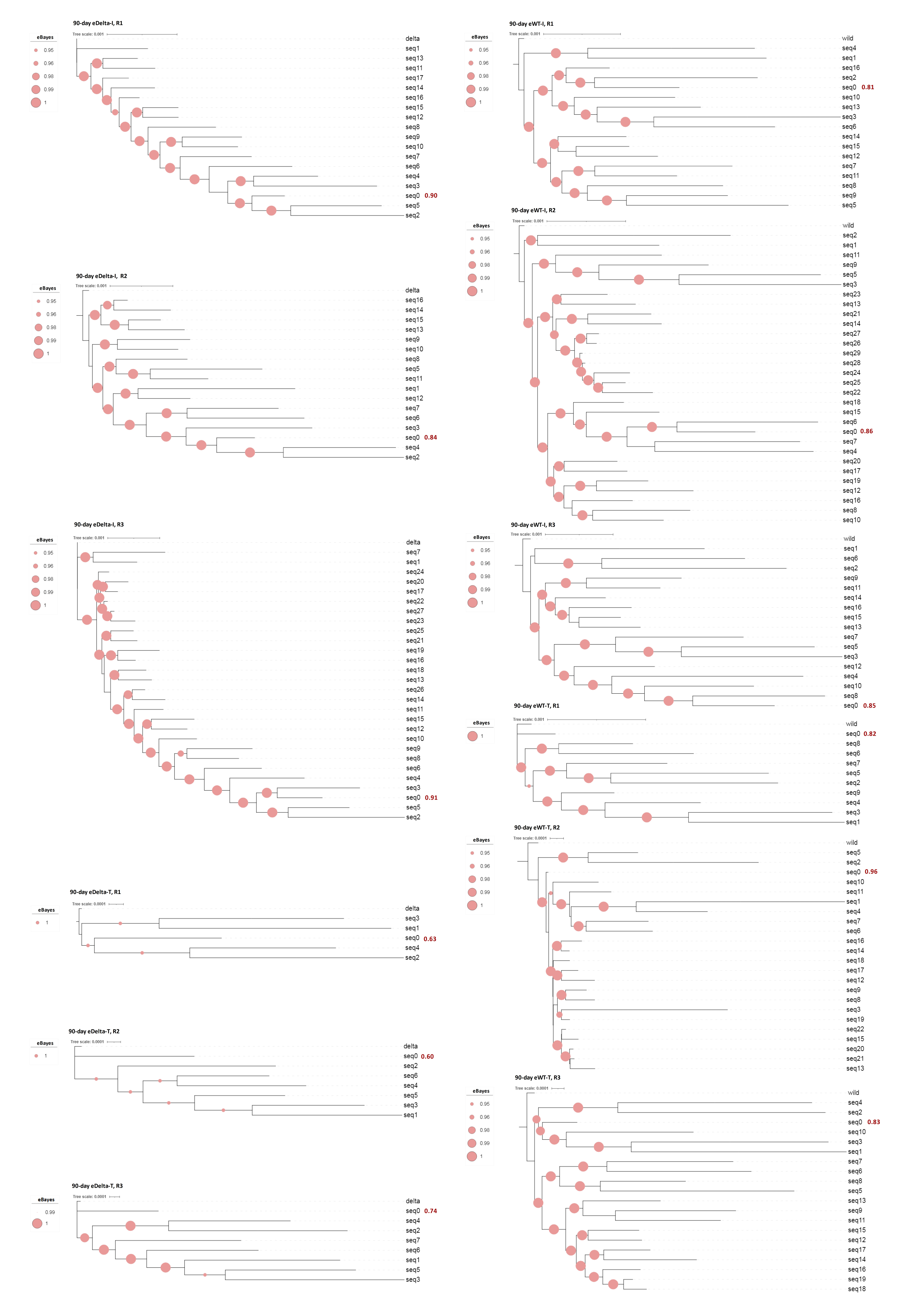

### Fig 4.jpg

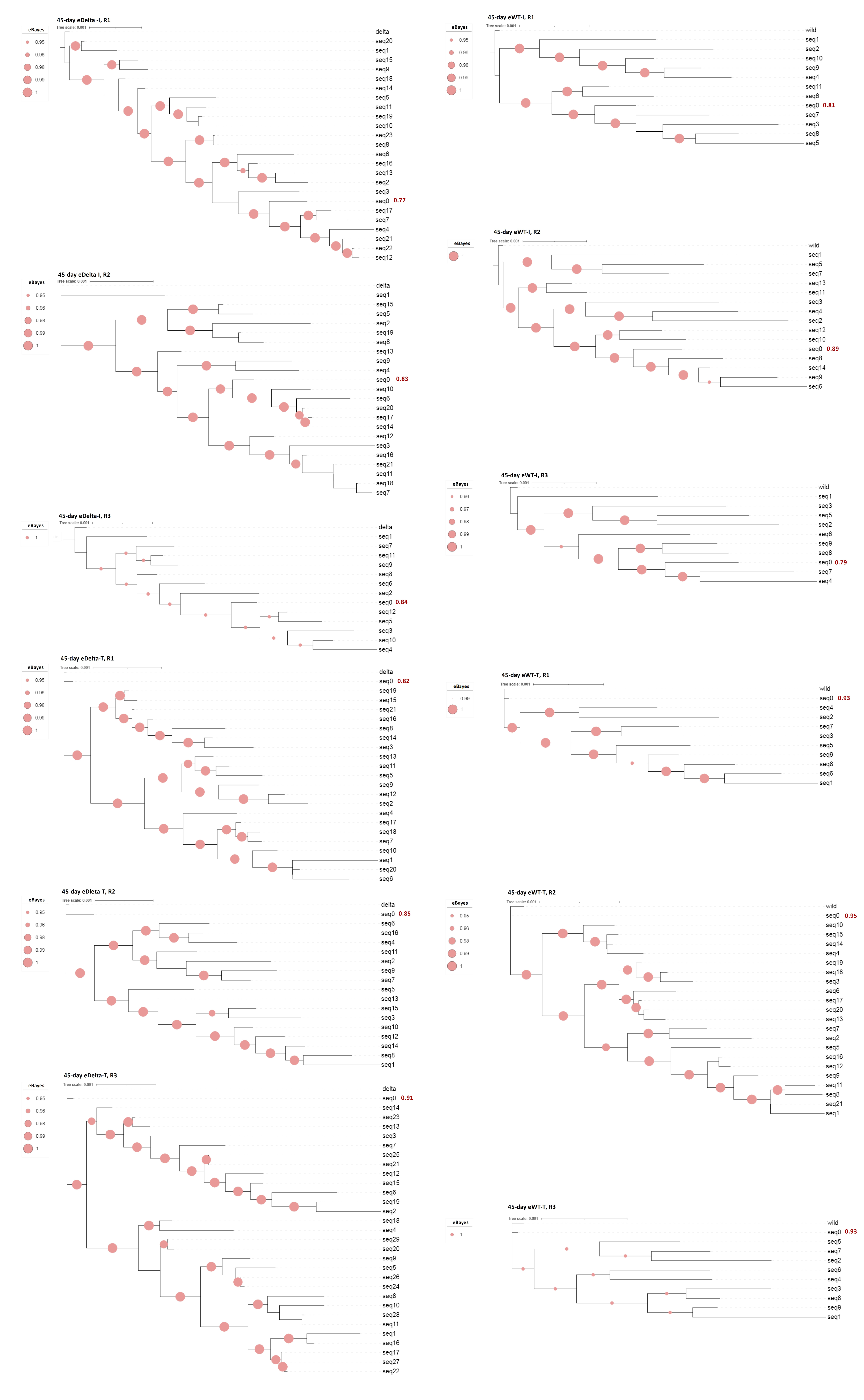
